# Demogenomic modeling of the timing and the processes of early European farmers differentiation

**DOI:** 10.1101/2020.11.23.394502

**Authors:** Nina Marchi, Laura Winkelbach, Ilektra Schulz, Maxime Brami, Zuzana Hofmanová, Jens Blöcher, Carlos S. Reyna-Blanco, Yoan Diekmann, Alexandre Thiéry, Adamandia Kapopoulou, Vivian Link, Valérie Piuz, Susanne Kreutzer, Sylwia M. Figarska, Elissavet Ganiatsou, Albert Pukaj, Travis J. Struck, Ryan N. Gutenkunst, Necmi Karul, Fokke Gerritsen, Joachim Pechtl, Joris Peters, Andrea Zeeb-Lanz, Eva Lenneis, Maria Teschler-Nicola, Sevasti Triantaphyllou, Sofija Stefanović, Christina Papageorgopoulou, Daniel Wegmann, Joachim Burger, Laurent Excoffier

## Abstract

The precise genetic origins of the first Neolithic farming populations, as well as the processes and the timing of their differentiation, remain largely unknown. Based on demogenomic modeling of high-quality ancient genomes, we show that the early farmers of Anatolia and Europe emerged from a multiphase mixing of a Near Eastern population with a strongly bottlenecked Western hunter-gatherer population after the Last Glacial Maximum. Moreover, the population branch leading to the first farmers of Europe and Anatolia is characterized by a 2,500-year period of extreme genetic drift during its westward range expansion. Based on these findings, we derive a spatially explicit model of the population history of Southwest Asia and Europe during the late Pleistocene and early Holocene.

**One-Sentence Summary:** Early European farmers emerged from multiple post LGM mixtures and experienced extreme drift during their westward expansion.

---

In recent years, genetic analyses of skeletal remains from early Neolithic farmers have fundamentally enriched our knowledge of the communities that became sedentary and invented food-production (*1, 2*). Nonetheless, the genetic origins of the first Western Eurasian farmers remain elusive. To date, palaeogenetic findings revealed that early European farmers were genetically distinct from Holocene European hunter-gatherers (HGs) (*3, 4*), with limited genetic exchange in early phases of the farming expansion, and more intense exchange at later stages (*5, 6*). Most continental European farmers likely descend from populations inhabiting the Aegean basin (*7*), but their ultimate genetic origins are still a matter of debate. Whereas early Neolithic Aegeans are clearly connected to Central Anatolians (*8*), they also show some affinities with Pre-Pottery Neolithic farmers of the southern Levant (*9*), suggesting a Near Eastern origin of these populations, potentially related to the spread of farming from the Fertile Crescent (*10*). However, Aegeans are genetically distant from early farmers from the Zagros region, the eastern wing of the Fertile Crescent, which was interpreted as evidence for the adoption of farming by genetically distinct groups of hunter-gatherers in Southwest Asia (*11*). There is however evidence of genetic continuity between Central Anatolian HGs and early farmers (*12*), indicating an earlier westwards migration from the Levant. Moreover, some Central Anatolian individuals show genetic affinities to Caucasus HGs as represented by the genome from Kotias in Georgia (*10, 13, 14*), who are thought to be closely connected to early Iranian Neolithic farmers (*9*) as well as to steppe populations (*15–17*).

These paleogenetic findings mostly stem from the interpretation of descriptive and summary statistics (e.g. Principal Component and admixture analyses, *f*-statistics), usually computed on low coverage genomes and/or on ascertained sets of SNPs (*18*) that are difficult to integrate into demogenetic analyses. In fact, previous analyses have estimated divergence times between ancestors of Neolithic and HG groups (*11, 14*), but the underlying models were very simplistic. Therefore, we still lack a detailed historical scenario of population demography, divergence and migration that is embedded in a spatially explicit and temporal framework, for the inference of which whole genomes of good sequencing quality are a prerequisite.

To reconstruct the ancestry of Neolithic populations of Southwest Asia and Europe and the processes that contributed to their differentiation from HGs, we therefore produced high-resolution genomes of early Holocene farmers and HGs (Table 1) arranged along a geographical and temporal transect from the Near East via the Aegean along the Danube to the Rhine (Fig. 1, Fig. S1A).

**Fig. 1.**
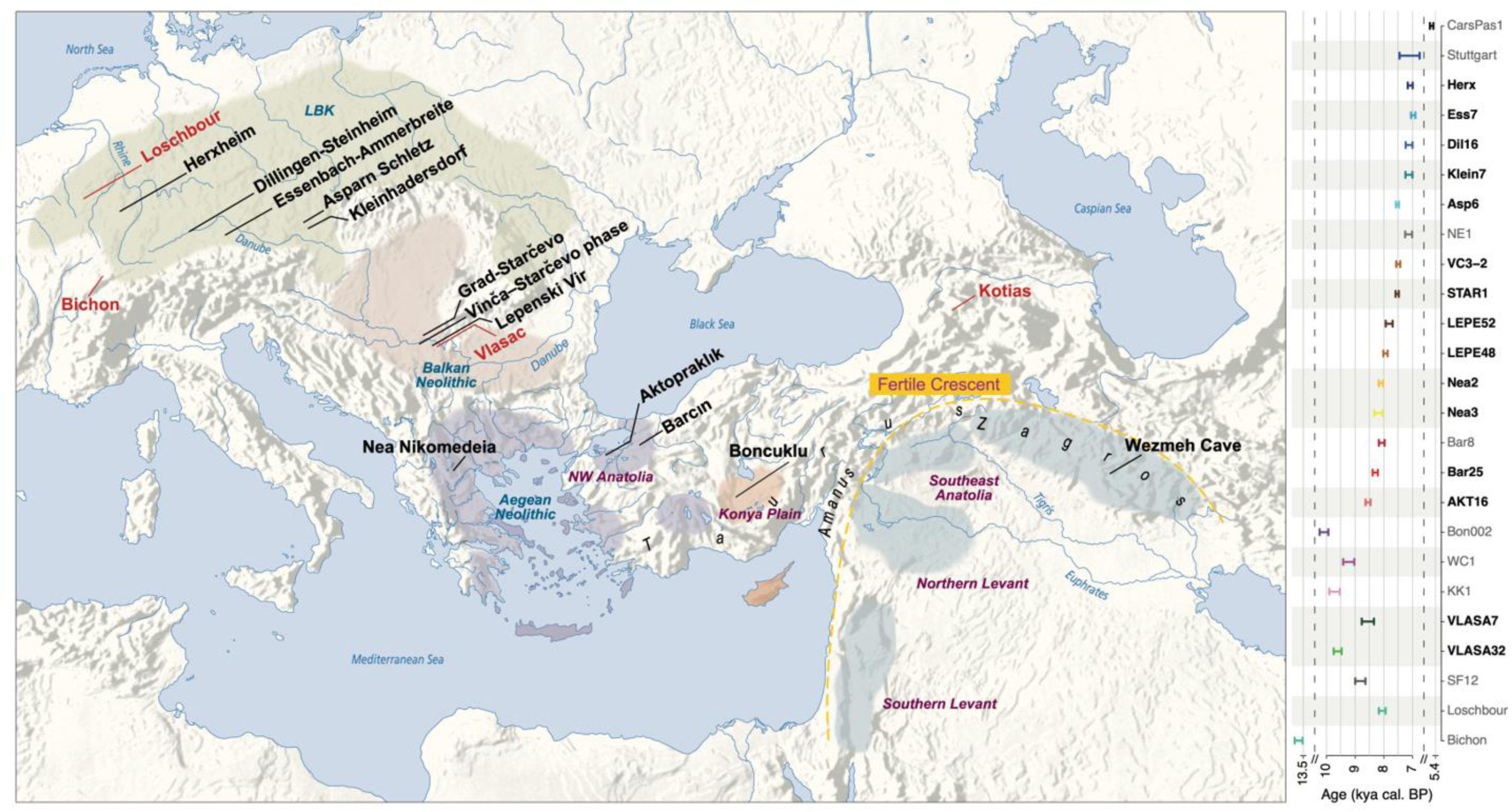
Spatial and temporal distribution of the palaeogenomes analyzed in this study. **Left**: Location of Neolithic (black), and Mesolithic or Palaeolithic (red) archaeological sites. Coloured areas represent different chronological phases of Neolithic expansion and cultures (blue) along the Danubian route of Neolithization; geographical areas are indicated in purple. **Right:** Chronological distribution of the 25 genomes analyzed in this study (see details in Table 1 and Table S2), with the 15 newly-sequenced genomes shown in bold. The chronological interval at 2 sigma (95.4% probability) is shown for each directly-^14^C dated sample, except for Stuttgart and Ess7, for which approximate dates are given based on the archaeological context.

**Table 1.**
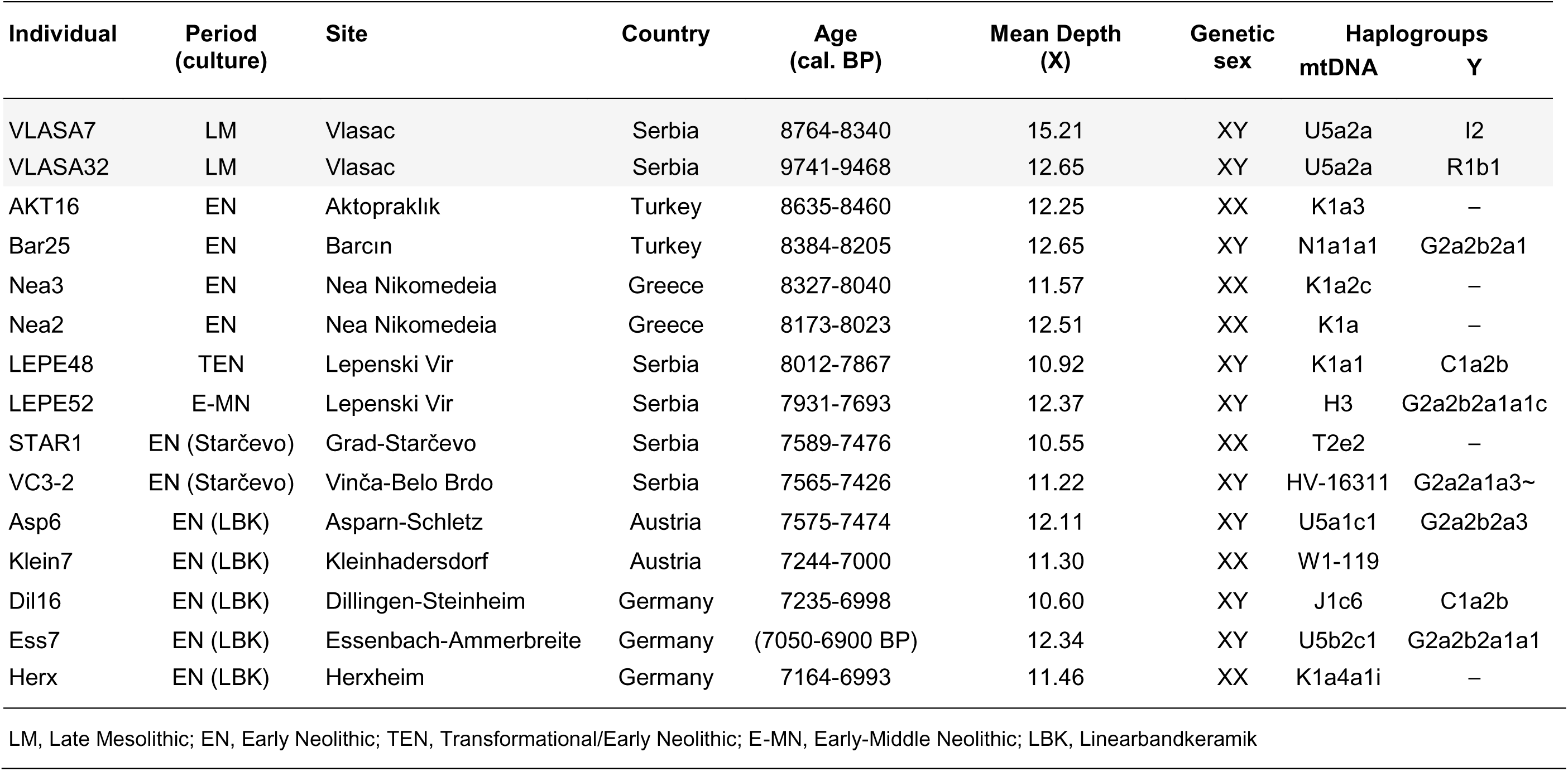
Archaeological and genetic information on the newly-sequenced genomes. Mesolithic samples are shown in grey. The samples’ ages are based on 14C dating (95.4% probability), except for Ess7 for which an approximate date is given based on the archaeological context (see Supp. Text- Archaeological context of the samples - Essenbach-Ammerbreite).

| Individual | Period<br>(culture) | Site | Country | Age<br>(cal. BP) | Mean Depth<br>(X) | Genetic<br>sex | Haplogroups<br>mtDNA | Y |
| --- | --- | --- | --- | --- | --- | --- | --- | --- |
| VLASA7 | LM | Vlasac | Serbia | 8764-8340 | 15.21 | XY | U5a2a | I2 |
| VLASA32 | LM | Vlasac | Serbia | 9741-9468 | 12.65 | XY | U5a2a | R1b1 |
| AKT16 | EN | Aktopraklık | Turkey | 8635-8460 | 12.25 | XX | K1a3 | – |
| Bar25 | EN | Barcın | Turkey | 8384-8205 | 12.65 | XY | N1a1a1 | G2a2b2a1 |
| Nea3 | EN | Nea Nikomedeia | Greece | 8327-8040 | 11.57 | XX | K1a2c | – |
| Nea2 | EN | Nea Nikomedeia | Greece | 8173-8023 | 12.51 | XX | K1a | – |
| LEPE48 | TEN | Lepenski Vir | Serbia | 8012-7867 | 10.92 | XY | K1a1 | C1a2b |
| LEPE52 | E-MN | Lepenski Vir | Serbia | 7931-7693 | 12.37 | XY | H3 | G2a2b2a1a1c |
| STAR1 | EN (Starčevo) | Grad-Starčevo | Serbia | 7589-7476 | 10.55 | XX | T2e2 | – |
| VC3-2 | EN (Starčevo) | Vinča-Belo Brdo | Serbia | 7565-7426 | 11.22 | XY | HV-16311 | G2a2a1a3~ |
| Asp6 | EN (LBK) | Asparn-Schletz | Austria | 7575-7474 | 12.11 | XY | U5a1c1 | G2a2b2a3 |
| Klein7 | EN (LBK) | Kleinhadersdorf | Austria | 7244-7000 | 11.30 | XX | W1-119 |  |
| Dil16 | EN (LBK) | Dillingen-Steinheim | Germany | 7235-6998 | 10.60 | XY | J1c6 | C1a2b |
| Ess7 | EN (LBK) | Essenbach-Ammerbreite | Germany | (7050-6900 BP) | 12.34 | XY | U5b2c1 | G2a2b2a1a1 |
| Herx | EN (LBK) | Herxheim | Germany | 7164-6993 | 11.46 | XX | K1a4a1i | – |

LM, Late Mesolithic; EN, Early Neolithic; TEN, Transformational/Early Neolithic; E-MN, Early-Middle Neolithic; LBK, Linearbandkeramik

## Results

### Genetic structure and affinities of ancient individuals

Multidimensional scaling (MDS) performed on the neutral portion of the genome (Fig. 2A) of modern and ancient individuals reveals three main ancient groups: i) a cluster of European HGs, ii) a cluster including an early Neolithic individual (WC1) from Wezmeh Cave in Iran and a Caucasus Mesolithic HG (KK1) from Western Georgia, and iii) a cluster with all other Holocene individuals. Consistent with previous analyses based on ascertained SNPs (*4, 19*), European and Anatolian Neolithic samples show strongest affinities with modern Sardinians, with the exception of one English (CarsPas1) and two NW Anatolian (Bar8, AKT16) individuals who are found closer to modern individuals from other parts of Southern Europe. In contrast, Palaeolithic and Mesolithic HGs are distinct from all modern Western Eurasians. The Iranian early Neolithic and the Caucasus Mesolithic HG individuals appear closest to modern genomes from Iran and the Caucasus, suggesting some genetic continuity in those regions.

**Fig. 2.**
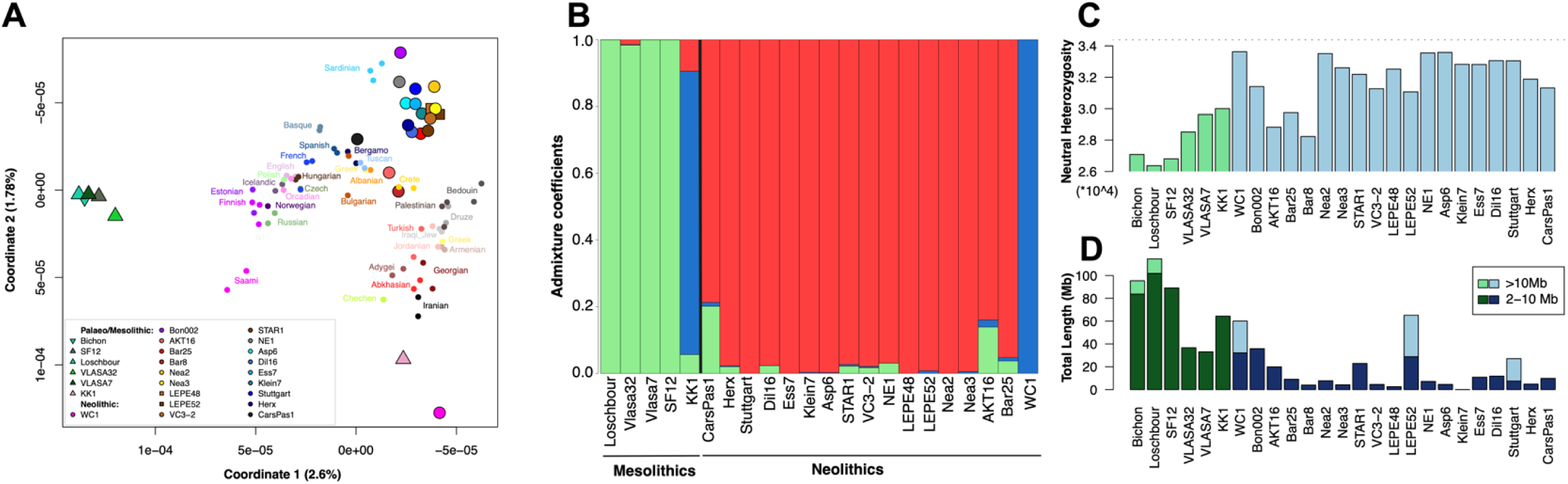
Genetic relationships and diversity of high quality ancient genomes. (**A**) Multidimensional Scaling (MDS) analysis performed on the neutrally evolving portion of ancient (n = 25) and modern (n = 65, shown as small circles) genomes from Europe and SW Asia. (**B**) Admixture analyses (K = 3) performed on 22 ancient genomes (three genomes with the lowest quality were discarded: Bichon, Bon002, Bar8). (**C**) Heterozygosity computed at neutral sites in ancient genomes (Palaeolithic and Mesolithic HGs in green, Neolithic individuals in blue); the median heterozygosity observed within modern genomes is indicated by the dashed line. (**D**) Runs of Homozygosity (ROHs) computed on imputed ancient whole genomes for short ROHs (2-10Mb, dark color) or long ROHs (>10Mb, light color).

### Demogenomic modeling assumptions

We first contrasted alternative models of population differentiation using high-resolution ancient genomes that were representative of the three main clusters described above. Sampled individuals from a given cluster were assumed to come from populations belonging to a large structured metapopulation, made up of interconnected but mostly unsampled populations, formally described as a continent-island model (Excoffier 2004; Excoffier et al. 2013). In this framework, the sampled islands receive a single pulse of gene flow from the continent shortly before sampling time. The three metapopulations were called *Western, Central* and *Eastern* and model the Western HGs, the Anatolian/Aegean/European early farmers, and the Iranian early farmers, respectively. Additional ancient individuals were then added successively to the initial model to infer their ancestry and various relationships with other individuals (see Supp. Text -Demographic inferences with *fastsimcoal2*). Thus, more complex models were progressively built, resulting in the historical scenario shown in Fig. 3A.

**Fig. 3.**
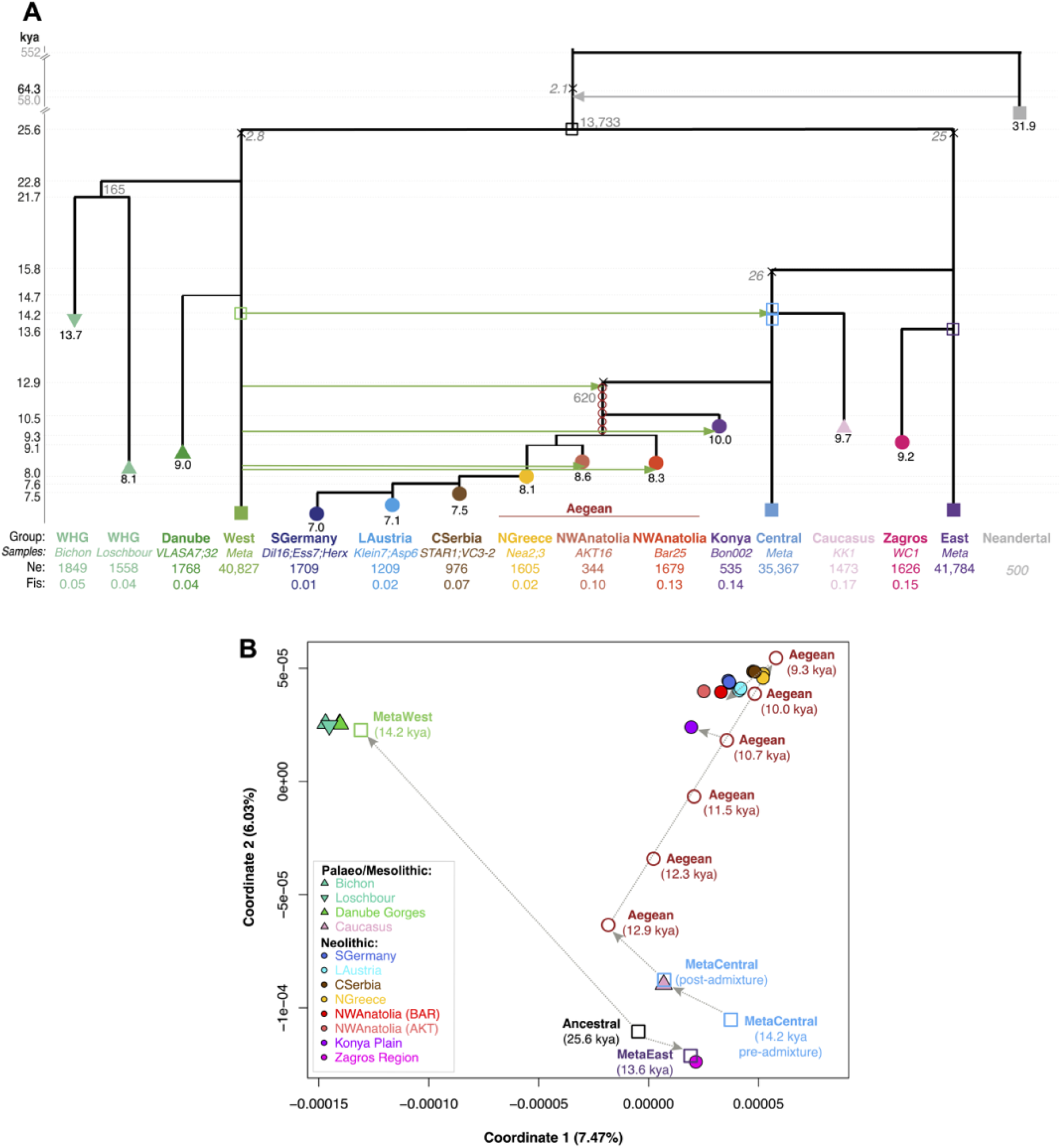
Demographic scenario inferred from the sampled genomes. (**A**) This demographic history was obtained by compiling the best models of all tested scenarios (see Supp. Text - Demographic inferences with *fastsimcoal2* - Final Model). Times of the events (y-axis) and population ages (shown below their symbol) are indicated in ky BP. Under each population name, we indicate their sampled genomes, their diploid population size and their associated inbreeding coefficient. Empty symbols indicate ancestral populations we simulated after or before key events (admixture or split). The X symbols indicate bottlenecks that occurred on ancestral branches, modeled as one generation bottleneck through a population of a size indicated in italic. Admixture proportions >10% from the *Western* metapopulation populations are represented with green arrows. (**B**) MDS analysis performed on data simulated according to the parameters of the scenario shown in pane A. Empty symbols are for simulated ancestral populations; grey arrows indicate the trajectory of the populations after admixture events and/or episodes of drift.

### All European and Anatolian farmers share a remote common ancestry with CHGs

Contrary to previous interpretations (*9*), we find that Caucasus HGs (represented by KK1) and early European and Anatolian farmers are related, and descend from a population ancestral to the *Central* metapopulation. This ancestral population received about 14% (95% CI 8-26%) of its gene pool from the *Western* metapopulation some 14.2 kya (95% CI 13.7-19.0, Fig. 3A, Fig. S31, Supp. Table 4). The ancestors of the Iranian Neolithic population were not affected by this initial admixture: they diverged from the *Eastern* metapopulation 13.6 kya (95% CI 11-24.6), from which the *Central* metapopulation already split ∼15.8 kya (95% CI 14.3-25.6). These analyses thus suggest that even though Caucasus HGs show closer genetic affinities with early Iranian farmers (Fig. 2A-B), they share a common ancestry with all Anatolian and European early farmers.

### Ancestors of early Anatolian and European farmers admixed twice with Western HGs

We find that the ancestors of early Anatolian and Aegean farmers, but not Caucasus HGs, received a second pulse of gene flow (15%, 95% CI 6-17) from the *Western* metapopulation ∼12.9 kya (95% CI 9.4-13.9), and all models not including this additional admixture are clearly rejected (Fig. S33, Supp. Table 4). Thus, the ancestors of early European and Anatolian farmers are the product of repeated episodes of gene flow from the *Western* metapopulation, and further diverged from Caucasus HGs due to an intense period of genetic drift between 13 and 9 kya (Fig. 3).

### Anatolian and Aegean farmers differentiation

Aegean populations from NW Anatolia (from the archeological sites of Aktopraklık and Barcın) and Northern Greece (from Nea Nikomedeia) seem to have diverged from each other at about the same time ∼9.1-9.3 kya (95% CI 9.1-12) shortly before they were sampled, potentially indicating a spatial diffusion of all Aegean populations around that time. Contrastingly, early farmers from Central Anatolia (represented by a genome from Boncuklu) have diverged earlier ∼10.5 kya (95% CI 10.5-11). Furthermore, Anatolian and Aegean populations show varying amounts of recent additional gene flow from the *Western* metapopulation, suggesting different levels of interaction with surrounding HGs. Indeed, genomes from Northern Greece show a lower degree of introgression (3%, 95% CI 1-11) than those from Boncuklu (10%, 95% CI 3-15), Barcın (12%, 95% CI 6-16), and especially Aktopraklık (17%, 95% CI 11-18) (Fig. S33-35-37, Supp. Table 4). The high level of *Western* metapopulation admixture found in Aktopraklık, a site previously described as influenced by both Epipalaeolithic and Neolithic traditions (*20*), is in line with the admixture analysis (Fig 2B) and *f*-statistics (Fig. S43).

### A stepwise, demic expansion of Neolithic farmers into Central Europe

To better understand the spread of early farmers into Europe, we modelled the genetic differentiation of three early Neolithic populations from Central Serbia, Lower Austria and Southern Germany (Fig. S38). We find that a simple model with a strict stepwise migration of early farmers originating in the wider Aegean region (NW Anatolia or Greece) and extending to Serbia along the Balkans and the so-called Danubian corridor, then to Austria, and eventually Southern Germany, is better supported than a scenario allowing for long-distance migration from the Aegean directly to Lower Austria (Fig. S39A, Supp. Table 4). We also find that early farming communities incorporated a few HG individuals (2-7%, compatible with previous estimates of 3-9% (*7, 21, 22*)) at each modeled stage of their dispersal along the Danubian corridor. Scenarios without *Western* metapopulation introgression into early farmer populations are rejected. Even though we modelled this introgression from the *Western* metapopulation, closely related to the Danube Gorges Mesolithic individuals from Vlasac, we cannot exclude that it actually occurred from other Western European HGs like those related to Loschbour and Bichon (Fig. 4H), since previous work suggested that different Mesolithic backgrounds could have introgressed into early farmer gene pools in different regions (*21*).

**Fig. 4.**
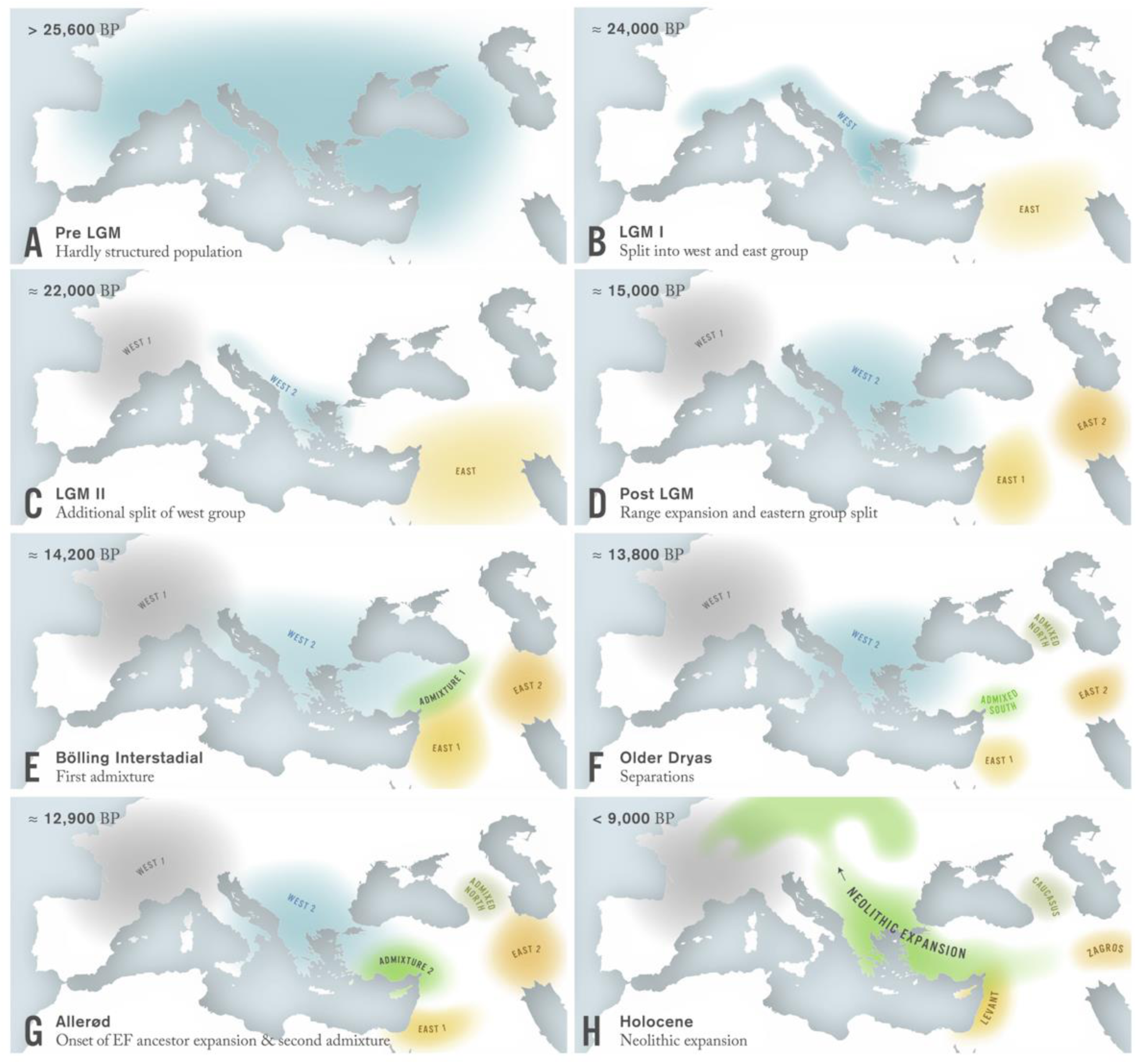
Possible scenario of the population history of SW Asia and Europe between the Last Glacial Maximum (LGM) and the early Neolithic period, i.e. approximately 26,000 to 7,000 years ago. Colored shaded areas indicate putative distributions of populations at different time points. See main text for a detailed description. Note that warmer periods (Bölling, Allerød interstadials; Holocene) correspond to population range expansions while colder periods (LGM, Older Dryas) are associated with contractions.

### An LGM divergence between Eastern and Western metapopulations

Our model also provides novel insights on the deep branching of pre-Neolithic populations. The divergence between the ancestors of the *Western* and *Eastern* metapopulations is estimated to date back to the LGM ∼25.6 kya (95% CI 17.3-31.3). This is much younger than the previously inferred divergence time between the ancestors of Iranian and Aegean early farmers (46-77 kya) (*11*) or between European early farmers and Western European HGs (27-76 kya) (*14*), which were both obtained under a simple model with constant population sizes and no admixture. We would get older divergence times (27-37 kya) between the ancestors of these two HG groups if we also used models without population size changes, but the fit to the data would be worse (Fig. S31A). Comparing two Western European HGs from the sites of Bichon and Loschbour to our newly-sequenced Mesolithic individuals from the Danube Gorges reveals that they already split during the LGM ∼22.8 kya (95% CI 16.8-24.7) (Fig. S41, Supp. Table 4), and that Bichon and Loschbour populations diverged from each other approximately a thousand years later.

### The reduced diversity in European hunter-gatherers is due to a massive LGM bottleneck

Genetic diversity as quantified by the heterozygosity at neutral sites is much lower in HGs than in early farmers, with the exception of NW Anatolians (Fig. 2C), in agreement with previous results (*10, 23*). Furthermore, HG genomes also show a generally larger proportion of short runs of homozygosity (2-10Mb ROHs, Fig. 2D, Fig. S20A) (*24*) indicative of remote inbreeding within European HGs and thought to be due to their small population size (*25*), in line with *MSMC2* analyses (Fig. S25). However, we estimate HG effective population sizes to actually be larger than those of most early farmers (Fig. 3A), particularly those from Anatolia (Bon002, AKT16 and Bar25) which show small effective population sizes in the order of a few hundred individuals in agreement with their relatively high proportion of short ROHs. The lower diversity observed among European HGs thus seems rather due to a very strong LGM bottleneck, which we estimate equivalent to a single human couple for one generation, or 20 individuals for 10 generations (Fig. 3A, Fig. S31, Supp. Table 4).

## Discussion

### Evolutionary insights gained from explicit demographic modeling

Our sequencing of ancient genomes at >10X, which triples the number of high quality whole genomes available for early Holocene in Europe, has allowed us to perform genetic analyses on an unbiased set of markers minimally impacted by selection, and thus ideally suited for reconstructing the population history of Western Eurasia from the Late Pleistocene to the Early Holocene. In addition to confirming some previous interpretations, our modeling provides several new insights about the demographic processes preceding and underlying the Neolithic transition and spread.

First, we found that European HGs had already split into two Western and Central subgroups ∼23 kya, and passed through a very severe LGM-driven bottleneck that is responsible for their low level of genetic diversity (Fig. 2C). Contrary to previous interpretations (*25, 31*), HGs are generally found to have larger population sizes than contemporary early farmer populations (Fig. 3A) which explains why they show close genetic affinities on the MDS (Fig. 2A) despite long divergence times and a wide geographic distribution. Conversely, it suggests that the Neolithic transition was linked to a reduction in local effective population sizes, potentially due to sedentarization (*32*) and restricted gene flow.

Second, we have evidenced that HGs from the Caucasus are phylogenetically related to the ancestors of early farmers from Europe and Anatolia as both show the traces of an ancestral admixture event between the *Western* and *Eastern* metapopulations (Figs. 2A-B). Despite this historical proximity, our model still predicts the observed affinity of the Caucasus HG genome to the Iranian Neolithic farmer (compare Fig. 2B and Fig. S42, as well as Fig. 2A and 3B).

Third, we can show how specific demographic processes affected the genetic divergence of past populations. Genomic data simulated under the final demographic scenario leads to population relationships very similar to those observed on an MDS (compare Fig. 2A and 3B) and a simplified admixture graph (*18*) compatible with all *f*-statistics calculated on the real data (Fig. S45-46), both providing an *a posteriori* validation of our model-based approach. By simulating additional genomes from unsampled populations or timepoints (Fig. 3), we further predict that the population ancestral to all sampled individuals was genetically close to ancestors of Iranian early farmers and Caucasus HGs; European HGs then considerably drifted after their LGM bottleneck, explaining their outlier position on the MDS. The ancestors of European and Anatolian early farmers, however, were first brought towards the center of the MDS by two consecutive admixture events with the *Western* metapopulation, and then pushed orthogonally towards the upper right by >2,500 years of intense genetic drift, possibly due to recurrent founder effects during their dispersal through Anatolia. Even though European and Anatolian early farmers were previously recognized to be genetically intermediate between other Near Eastern groups (*12*), or considered to be a mixture of other ancient populations (*6*), this initial admixture signal remained hidden to previous approaches as it was eroded by later genetic drift (Fig. S42): Whereas populations simulated shortly after the two main admixture events with the *Western* metapopulation indeed appear as admixed (i.e. MetaCentral (489g), Aegeans (445g) on Fig. 3B), this signal progressively disappears through time, and these populations eventually look like having a completely independent gene pool (e.g. in the admixture analyses of Fig. S42). Their more central position on the MDS later on is then due to admixture with the *Western* metapopulation and surrounding farmers modeled as the *Central* metapopulation.

### A spatially explicit scenario of population differentiation

Even though our demogenomic model (Fig. 3A) can explain observed population affinities, we are aware that it has temporal and geographical gaps that will only be filled by gathering further genomic data. In particular, high quality samples from the Northern and Southern Levant as well as representatives of Eastern HGs are needed to complete and confirm our conclusions.

Nonetheless, the timing and sequence of demographic events that emerge from our analyses suggest a spatially explicit scenario of population differentiation during the LGM and early Holocene (Fig. 4). Under this scenario, the very strong bottleneck in the population ancestral to European HGs that occurred after their divergence from the *Eastern* metapopulation was probably due to a contraction into a small LGM refugium, potentially located in milder Western Mediterranean coastal regions (Fig. 4B). It was followed by a rapid differentiation of these HGs in two separate refugia (Fig. 4C), corresponding to what archaeologists identified as the areas of distribution of Solutrean and Epigravettian lithic traditions in Europe (*26, 27*).

Following a period of range expansion after the LGM (Fig. 4D), representatives of the *Central* metapopulation, likely descendants of the Epigravettian refugial population, mixed ∼14.2 kya with the population ancestral to both Caucasus HGs and Anatolian/Aegean early farmers. Given the former geographical distribution of the glacial refugia, this admixture likely happened during the Bölling interstadial period in a region encompassing Southeastern Anatolia and the Northern Levant or even in neighboring regions such as Central and Eastern Anatolia or the Turkish South coast (Fig. 4E).

The subsequent demographic processes explaining the differentiation between Central Anatolian and Aegean farmers are more difficult to pinpoint and locate. The inferred low population size of the ancestors of Anatolian/Aegean farmers after their split from Caucasus HGs during the Older Dryas (∼12.9 kya; Figs. 3A and 4F) could be due to a westward range expansion and associated recurrent founder effects during the Allerød interstadial, a period with relatively favorable climatic conditions during which they would have also further admixed with Epipalaeolithic HGs in Anatolia (Fig. 4G). The fact that early farmers from Central Anatolia share the same admixture event and drift with Aegean farmers suggests they were part of the same expansion wave. However, their close genetic proximity with Epipalaeolithic Central Anatolian foragers (*12*) questions this scenario and rather indicate that admixed groups existed in Central Anatolia in pre-Neolithic times, or had moved there from the Fertile Crescent, adopting fully developed farming practices at a later stage. Indeed, early aceramic sites like Boncuklu and Aşıklı on the Anatolian Plateau show experiments in crop cultivation and caprine management ∼9.7 kya (*28, 29*).

In contrast, the migration to NW Anatolia (Fig. 4H) likely occurred at the time of the fully developed ceramic Neolithic characterized by the establishment of widespread mixed farming (*30*). Further support for such a demic diffusion scenario to NW Anatolia by a direct (coastal) route and not via the Konya plain region comes from *f*-statistics showing Levantine populations sharing more drift with Aegeans than with Central Anatolian Neolithic individuals (Fig. S48). This signal could either be due to i) some long distance gene flow between the Aegeans and a Levant-like population, ii) a higher level of *Western* metapopulation admixture observed in Boncuklu (Fig. S47), or iii) an early migration of the Boncuklu ancestors from the Fertile Crescent to Central Anatolia, combined with some later gene flow between people from the Fertile Crescent and the ancestors of Aegeans. However, *f*-statistics analyses reveal that early farmers from the Aegean are rather heterogeneous in their levels of shared drift with several populations, including Levantine HGs and early Iranian farmers (Fig. S49), suggesting that the Neolithization of the Aegean was a more complex process. Nonetheless, our results imply that even though the initial spread of the Neolithic must have been through cultural diffusion in the Fertile Crescent among genetically already well differentiated groups, its extension to other parts of Anatolia and along the Danubian corridor occurred through demic diffusion processes (Fig. 4H).

In sum, our population modelling allowed us to extract novel, unexpected, but complementary and far more detailed information on population affinities than what one could conclude from summary statistics or multivariate analysis alone. In addition, it provides a time frame for the differentiation of the major groups populating SW Asia and Europe from the LGM until the introduction of agriculture, and highlights the crucial role of climatic changes in promoting population fragmentation and secondary contacts (*33*).

## Acknowledgments

We are grateful to Martina Unterländer and Aleksandra Žegarac for help with sample preparation. Lara Cassidy and Kay Prüfer kindly provided access to unpublished raw sequencing data. We thank Ourania Palli and Franz Pieler for useful archaeological information. We thank the General Directorate of Cultural Heritage Rhineland-Palatinate, the Anthropological Department of the Natural History Museum Vienna, Michaela Harbeck and the Bavarian State Collection for Anthropology and Palaeoanatomy, Munich, for providing skeletal samples. We also acknowledge the use of the sequencing platform at the University of Berne for whole genome sequencing services and support, the IBU cluster of the University of Berne for NGS data analysis (https://www.bioinformatics.unibe.ch/), as well as the UBELIX HPC cluster of the University of Berne (http://www.id.unibe.ch/hpc) for demographic model analyses, and the supercomputer Mogon of Johannes Gutenberg University Mainz (https://hpc.uni-mainz.de).

## Funding

Swiss SNSF grant No. 310030_188883 (LE, NM)

Swiss SNSF grant No. 31003A_173062 (DW, IS, VL, CR)

Swiss SNSF grant No. <u>310030_200420</u> (DW)

German Science Foundation BU 1403/6-1 (JBu, CP, SK)

Humboldt foundation (JBu, CP)

Greek-German bilateral agreement (GSRT and BMBF) project ’BIOMUSE’ 5030121 (CP, JBu, LW, EG, YD)

Serbian Ministry of Science project ID III47001 (SS)

ERC BIRTH project 640557 (SS)

Marie Skłodowska-Curie actions ITN ’BEAN’ (JBu, SS, SF, CP, ZH)

Marie Skłodowska-Curie individual fellowship (MB: 793893 ’ODYSSEA’)

EMBO Long-Term Fellowship (ZH: ALTF 445-2017)

Seal of Excellence Fund grant from the University of Berne (NM: SELF2018-04).

Mainz University (LW)

## Author contributions

Conceptualisation: JBu, DW, LE.

Resources (samples, archaeological and anthropological context): SS, MB, CP, ST, NK, FG, AZL, JPec, JPet, EL, MT-N

Data production: LW, SF, SK

Data curation: IS, VL, AT, AK

Formal analyses, Investigation & Visualisation: NM, LE, AK, EG, TJS, RNG, VP, JBu, JBl, YD, ZH, IS, AP, CRB, DW

Writing – original draft: NM, MB, DW, JBu, LE

Writing – review & editing: All co-authors.

## Data and materials availability

Sequences will be made available at the European Nucleotide Archive. ATLAS pipeline commit 6df90e7 is available at: https://bitbucket.org/wegmannlab/atlas-pipeline/src/master/. Other bioinformatic codes will be made available upon request.

## Supplementary Materials

Materials and Methods

Supplementary Text

Supplementary Tables Legend

Supplementary Tables 1-5

