## Supplementary Information for "Demogenomic modeling of the timing and the processes of early European farmers differentiation"

**This PDF file includes:**

Materials and Methods
Supplementary Text
Figs. S1 to S49
Tables S1 to S10

**Other Supplementary Materials for this manuscript include the following:**

Supplementary Tables Legend
Supplementary Tables 1-5

### Materials and Methods

#### ***DNA preparation and sequencing***

In this study, we present new whole-genome sequences for 15 ancient human individuals (see Table 1 for an overview).

Paleogenetic analyses were conducted at the Institute of Organismic and Molecular Evolution (Johannes Gutenberg University, Mainz). All laboratory steps prior to PCR amplification (sample preparation, DNA extraction and library preparation) were carried out in the dedicated ancient DNA facilities of the Palaeogenetics Group, which are separated from post-PCR areas. Strict ancient DNA protocols to prevent and detect contamination with modern DNA as well as cross contamination between samples were applied as previously described (*3*, *34*, *35*). These ancient DNA standards include decontamination of workspace, labware and samples, independent DNA extractions and the processing of blank controls during sample pulverization, DNA extraction, library preparation and PCR reactions to monitor contaminations.

##### Sample preparation

The ancient DNA samples were extracted from petrous bones (*Pars petrosa*) which were prepared as described in Hofmanová *et al. (7)*. In detail, after documentation, the bone samples were sterilized under ultraviolet light (254 nm) from 2 sides for 45 minutes per side. In order to remove superficial contaminants, soil remains and the outer bone surface were removed using a sandblasting machine (P-G 400, Harnisch & Rieth, Winterbach, Germany) with Spezial-Edelkorund (EW60/250 my and 30B/50 my; Harnisch+Rieth). The densest, inner part of the petrous bone was then isolated and cut in cubes using a disk saw (Electer Emax IH-300, MAFRA). After irradiation with ultraviolet light (254 nm) from two sides, for 45 minutes per side, the densest bone cubes were pulverized using a mixer mill (MM200, Retsch). To control contamination, blank milling controls containing hydroxyapatite were processed in parallel.

##### DNA extraction

DNA extraction followed the protocol by Yang *et al. (36)* with the modifications described in MacHugh *et al. (37)* and Gamba *et al. (38)* as well as additional modifications described below. For each sample, 0.15-0.31 g of bone powder was used for extraction. Prior to extraction, a pre-lysis was performed by adding 1 mL of EDTA (0.5 M, pH8, Ambion/Applied Biosystems, Life technologies, Darmstadt, Germany) to the bone powder and incubating at room temperature for 10 minutes. The solution was centrifuged at maximum speed to pellet the powder and the supernatant was removed. Lysis was performed on rocking shakers at 37°C for 24 hours (900 rpm) using 1 ml of extraction buffer containing EDTA (950 µl, 0.5 M, pH8, Ambion/Applied Biosystems, Life technologies, Darmstadt, Germany), Tris-HCl (20 µl, 1 M, pH8, Life Technologies, Carlsbad, United States), N-Laurylsarcosine (17 µl, 5%, Merck Millipore, Darmstadt, Germany) and Proteinase K (13 µl; 20 mg/ml, Roche, Mannheim, Germany). After 24 hours of incubation, the samples were centrifuged for 10 minutes at 10,000 rpm, the supernatant was removed, transferred into new tubes and stored in a fridge until further processing. A second lysis step was performed following the same procedure. Following lysis, the supernatants from the two lysis steps were merged on an Amicon Filter (Amicon Ultra-4 30 kDA, 15 ml, Merck Millipore, Darmstadt, Germany) and centrifuged for 10 min at 2500 rpm. The DNA was then washed twice with 3 ml 1X Tris-EDTA, followed by centrifugation at 2500 rpm for 20 minutes and discarding of the flow-through in between. After washing, the extract was concentrated to 100 µl and subsequently purified with the QIAgen MinElute kit (Qiagen, Venlo, Netherlands) following the manufacturer's instructions but incubating for 5 minutes during elution with 44 µl elution buffer (preheated to 65°C). Blank controls were processed during DNA extraction and incorporated into all further steps of the analyses.

**Library preparation**

The libraries were sequenced on an Illumina HiSeq3000 (SE, 100 cycles or PE, 150 cycles) at the Next Generation Sequencing Platform of the University of Berne, Switzerland.

Double-indexed Libraries were prepared according to the protocol by Kircher *et al. (39)* with slight modifications. Prior to library preparation the DNA extracts were treated with USER™ enzyme: 5 µl of USER™ enzyme (New England Biolabs, Ipswich, Massachusetts, United States) was added to 16.25 µl of DNA extract (exception: 17µl of two merged extracts were used for library SL3_5 of sample AKT16, see Supp. Table 1) and the mixture was incubated for three hours at 37°C (*40*). The blunt-end repair step followed immediately and was performed using the NEBNext End Repair Module (New England Biolabs, Ipswich, Massachusetts, United States): the DNA extract was mixed with NEBNext End Repair Reaction Buffer (10X, 7 µl), NEBNext End Repair Enzyme Mix (3.5 µl) and nuclease-free water (38.25 µl; for a final reaction volume of 70 µl) and incubated for 15 minutes at 25°C followed by 5 minutes at 12°C. Deviating from this procedure, two libraries (SL2.12_MU and SL3.12_MU of sample VC3-2) were produced using 20 µl DNA extract and without USER™ enzyme treatment of the DNA extract (see Supp. Table 1). In the adapter ligation step hybridized adapters P5 and P7 (IDT, Leuven, Belgium) were used at a concentration of 0.75 µM. 3 µl of Fill-In product (total volume: 40 µl) were amplified using AccuPrime™ Pfx SuperMix (20 µl; Thermo Fisher Scientific, Waltham, Massachusetts, United States) in one PCR parallel adding unique and sample-specific index combinations to the library molecules (final reaction volume: 25 µl; final primer concentration: 200 nM or 160 nM each). Double indexing followed Kircher *et al. (39)*, but using index sequences from the NexteraXT index Kit v2 (Illumina, San Diego, California, United States; barcode length 8 bp; ordered at IDT, Leuven, Belgium). The PCR was performed in 9-13 cycles; the PCR temperature profile followed the manufacturer's recommendations but using an annealing temperature of 60°C, extending for 30 seconds during each cycle and performing a final elongation step for 5 minutes. For purification during library preparation the MinElute PCR Purification Kit (Qiagen, Hilden, Germany) was used, while amplified libraries were purified with the MSB® Spin PCRapace (Invitek, Stratec Molecular, Berlin, Germany). Libraries were quantified by Qubit® Fluorometric quantitation (dsDNA HS assay, Invitrogen, Carlsbad, California, United States) and measurement on the Agilent 2100 Bioanalyzer System (HS DNA, Agilent Technologies, Waldbronn, Germany). Blank controls as well as positive controls (nonsense hybrids) of known concentration, were processed in every library step including PCR amplification to verify the success of the library preparation and to monitor contamination. Quantification of blank controls did not indicate significant contamination during any laboratory step (pulverization, DNA extraction, library preparation and amplification).

**Sample Screening**

Using shallow shotgun sequencing on an Illumina MiSeq™ platform at StarSEQ GmbH (Mainz, Germany), all samples were screened for their endogenous DNA preservation. Libraries were equimolarly pooled, subsequently purified with magnetic beads (Agencourt® AMPure® XP beads, Beckmann Coulter) and sequenced in single-end runs with 50 bp read length. Demultiplexing was performed by the sequencing facility using the MiSeq Reporter and allowing one mismatch in the barcode. Blank controls from DNA extraction, library preparation and PCR-steps were sequenced alongside to estimate the fraction of potentially contaminating reads introduced in the lab (for details see Supp. Table 1).

**Whole-genome sequencing**

For deeper shotgun sequencing 2-5 DNA extracts and 3-9 libraries of each sample were prepared (as detailed above). The libraries were amplified in up to 13 PCR parallels, each with an individual index combination, to increase the complexity of the libraries. Subsequently, PCR parallels were purified and quantified individually as described above.

For sequencing, PCR parallels were pooled equimolarly according to their concentrations measured on Qubit® and purified with magnetic beads (Agencourt® AMPure® XP beads, Beckmann Coulter). The samples were sequenced on an Illumina HiSeq3000 (SE, 100 cycles or PE, 150 cycles) at the Next

Generation Sequencing Platform at the University of Berne, Switzerland. The lab work and the sequencing strategy for each sample are summarized in Table S1; for details see Supp. Table 1.

**Table S1 - Summary of lab work and sequencing strategy.**

| Sample ID | # Extracts | # Libraries | # PCR parallels | # HiSeq3000 Lanes |
|---|---|---|---|---|
| AKT16 | 4 (+1 merged) | 8 | 98 | 8 |
| Bar25 | 5 | 5 | 58 | 5 |
| Nea2 | 4 | 4 | 41 | 3 |
| Nea3 | 4 | 4 | 42 | 3 |
| VLASA32 | 2 | 4 | 52 | 4 |
| VLASA7 | 2 | 4 | 52 | 4 |
| LEPE48 | 2 | 3 | 35 | 3 |
| LEPE52 | 5 | 7 | 72 | 4 |
| STAR1 | 2 | 3 | 36 | 3 |
| VC3-2 | 4 | 6 | 60 | 4 |
| Asp6 | 2 | 3 | 35 | 3 |
| Klein7 | 4 | 5 | 50 | 3 |
| Ess7 | 4 | 6 | 69 | 4 |
| Dil16 | 4 | 5 | 47 | 3 |
| Herx | 5 | 9 | 110 | 6 |

***<u>Genotyping, quality checks</u>***

**Samples processed**

We analyzed a total of 102 high-depth samples, consisting of:

1. 77 modern individuals from the Simons Genome Diversity Project (SGDP, https://www.simonsfoundation.org/simons-genome-diversity-project/) (*41*), mostly from Europe and SW Asia (Fig. S1B), but we also included two Africans (Mende), six Asians (two Tubalars, two Kapus, two Hans), two Oceanians (Papuans) and two Amerindians (Karitianas) for comparison.
2. 15 ancient individuals sequenced for this study at 10X - 15X (see Table 1)
3. The following ten other high-quality ancient (Palaeolithic, Mesolithic and Neolithic) individuals previously published (see Table S2; Fig. S1A):
    a. Bichon and KK1 (*14*), demultiplexed FASTQ files downloaded from ENA with run accessions ERR1078331-ERR1078351 and ERR1078321-ERR1078325, respectively.
    b. Loschbour and Stuttgart (*6*), demultiplexed FASTQ files kindly provided by the authors (contact: Kay Prüfer).
    c. SF12 (*26*), BAM file downloaded from ENA with run accession ERR2060277.
    d. Bon002 (*10*), unaligned FASTQ file downloaded from European Nucleotide Archive (ENA) with run accession ERR1514027.
    e. WC1 (*11*), demultiplexed FASTQ file kindly provided by the authors (contact: Jens Blöcher, co-author of the study).
    f. Bar8 (*7*), demultiplexed FASTQ file kindly provided by the authors (contact: Zuzana Hofmanová, co-author of the study).
    g. NE1 (*38*), raw sequencing FASTQ file kindly provided by the authors (contact: Lara Cassidy).
    h. CarsPas1 (*42*), demultiplexed FASTQ file kindly provided by the authors (contact: Yoan Diekmann, co-author of this study).

**Table S2 - General and genetic information on the 10 ancient genomes from the literature used in our study**, Palaeolithic and Mesolithic shown in grey, Neolithic in white. Note that the mean depth was recalculated from the pipeline described in this study.

| Individual | Period (culture) | Site | Country | Age (cal. BP) | Mean Depth (X) | Genetic sex | Haplogroups mtDNA | Haplogroups Y | Phenotype (Eye; Hair) | Publication |
|---|---|---|---|---|---|---|---|---|---|---|
| Bichon | LUP | Grotte du Bichon | Switzerland | 13,770-13,560 | 7.47 | XY | U5b1h | I2 | Br; D | Jones, 2015 |
| SF12 | Meso | Stora Förvar | Sweden | 9,033–8,757 | 51.21 | XX | U4a1 | – | Br; D | Günther, 2018 |
| Loschbour | LM | Heffingen | Luxembourg | 8,170-7,940 | 22.43 | XY | U5b1a | I2a1b* | Bl; D | Lazaridis, 2014 |
| Bon002 | Neo | Boncuklu | Turkey | 10,229-9,927 | 5.45 | XX | K1a | – | – | Kilinc, 2016 |
| WC1 | EN | Wezmeh Cave | Iran | 9,405-9,032 | 12.66 | XY | J1d6 | G2b | Bl; D | Broushaki, 2016 |
| Bar8 | EN | Barcın | Turkey | 8,162-7,980 | 7.37 | XX | K1a2 | – | Br; D | Hofmanova, 2016 |
| NE1 | MN (ALP) | Polgar-Ferenci-hat | Hungary | 7,260-7,020 | 17.66 | XX | U5b2c | – | Br; D | Gamba, 2014 |
| Stuttgart | Neo (LBK) | Viesenhäuser Hof, Stuttgart-Mühlhausen | Germany | 7,050-6,750* | 15.58 | XX | T2c1d1 | – | Br; D | Lazaridis, 2014 |
| CarsPas1 | EN | Carsington Pasture Cave, Brassington | England | 5,606-5,471 | 10.14 | XY | J1c1 | I2a2a1a1a | Br; D | Brace, 2019 |

LUP, Late Upper Palaeolithic; Meso, Mesolithic; LM, Late Mesolithic; Neo, Neolithic; EN/MN, Early/Middle Neolithic; LBK, Linearbandkeramik; ALP, Alföld Linear Pottery

Br, Brown eyes; Bl, Blue eyes; D, Dark hair (listed as "Brown or Black", "Black/Dark", "Dark" in the original publications)

*approximate date based on the archaeological context

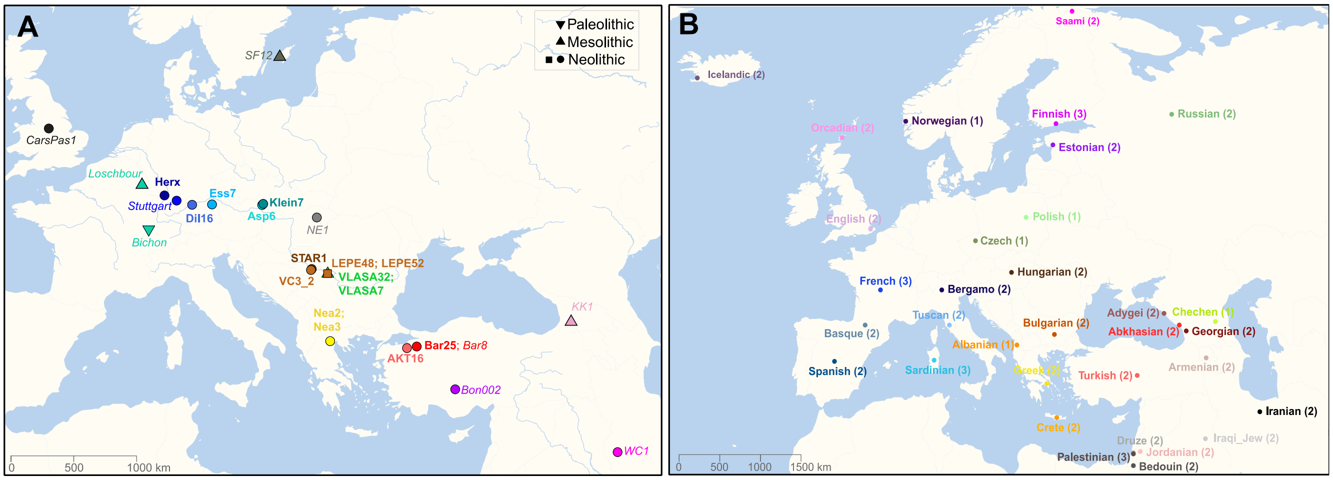


**Figure S1 - Geographical distribution of all genomes included in this study.** (**A**) 25 high-quality ancient genomes: 15 newly sequenced (bold) and 10 from the literature (italic). (**B**) 65 modern genomes from Europe and SW Asia that were selected within the SGDP panel.

### Programs used

All bioinformatic steps were conducted with the following programs and versions:

- *fastqc* - version 0.11.5, (www.bioinformatics.babraham.ac.uk/projects/fastqc/)
- *Trim Galore!* - version 0.4.3, (https://github.com/FelixKrueger/TrimGalore)
- *bwa - Burrows-Wheeler Alignment Tool* - version 0.7.15 (*43*)
- *SAMtools* - version 1.3 (*44*)
- *Picard-tools* - version 2.9, http://broadinstitute.github.io/picard/
- *GATK* - version 3.7 (*45*)
- *ATLAS* - version 1.0, commit 6bd2482 (*46*)

- *Snakemake - version 4.0 (47)*
- *seqtk* - version 1.2 (https://github.com/lh3/seqtk)
- *ContamMix - version 1.0 (48)*
- *MIA (Mapping Iterative Assembler)* - version 1.0 (https://github.com/mpieva/mapping-iterative-assembler )
- *mafft* - version 7.31 (*49*)
- *ANGSD* - version 0.917 (*50*)

Some of the steps were performed with commit 6df90e7 of the *snakemake* script *ATLAS-Pipeline* available at bitbucket.org/wegmannlab/atlas-pipeline, as indicated below.

**Bioinformatics Pipeline**

All 102 ancient and modern samples were processed with the same pipeline, albeit minor changes due to variation in how the raw data were obtained. We therefore first describe the pipeline as used on the 15 samples sequenced in this study, followed by a description of how previously published data were analyzed. The pipeline consisted of the following 8 steps.

Step 1: Raw data handling

Raw, demultiplexed sequencing data were processed per PCR-parallel as follows:

1. The sequencing quality was checked with fastqc.
2. Raw reads were trimmed using *trimgalore* with no quality filter and a length filter of 30 bp ( *-q 0 , --length 30, -a 'AGATCGGAAGAGCACACGTCTGAACTCC'* ). For paired-end libraries the additional option --retain unpaired was used and the second adapter sequence (-a2 'AGATCGGAAGAGCGTCGTGTAGGGAAAG') was provided.
3. A second fastqc analysis was performed to verify trimming and the quality of the remaining sequences.
4. Reads were then aligned to the 1000 genomes version of the human reference genome hs37d5 (ftp://ftp.1000genomes.ebi.ac.uk/vol1/ftp/technical/reference/phase2_reference_assembly_sequence/hs37d5.fa.gz (*51*)) using *bwa -mem* with options *-t 8 -M*. Reads with a mapping quality below 30 were filtered out using *SAMtools*.
5. Duplicate reads were marked but not removed using *picard-tools MarkDuplicates* with *VALIDATION_STRINGENCY=SILENT* and *AS=TRUE*.
6. Unmapped reads were removed with *SAMtools* with option *-F4*. Informative read groups were added to keep track of individual PCR-parallels with *picard-tools AddOrReplaceReadGroups*.

7. All necessary sorting and indexing in-between the steps was performed with *SAMtools*, library-files from the same samples were merged using *SAMtools merge*.

Step 2: Local Realignment

Local indel-realignment is an important but computationally expensive step in our pipeline. In order to allow for parallelization, we proceeded as follows. First, we identified potential target intervals using *GATK RealignerTargetCreator* from a set of ten ancient and ten modern samples: Asp6, Bar25, Dil16, Ess7, Klein7, LEPE48, Nea2, Nea3, STAR1, VC3, VLASA32, VLASA7, French-1, Sardinian-1, Russian-1, Spanish-1, Georgian-1, Hungarian-1, Icelandic-1, Finnish-1, English-1, Greek-1, Mende-1, Estonian-1, Polish-1 and providing known indel sites identified by the 1000 Genomes project (*ftp://gsapubftp-anonymous@ftp.broadinstitute.org/bundle/b37/1000G_phase1.indels.b37.vcf.gz, ftp://gsapubftp-anonymous@ftp.broadinstitute.org/bundle/b37/Mills_and_1000G_gold_standard.indels.b37.vcf.gz(51)*). We refer to the obtained target set as *TargetsBase.* Second, we created a set of reads by downsampling the following 5 ancient and 5 modern samples to a depth of 4X: Bar25, Klein7, STAR1, VLASA32, VLASA7, French-1, Georgian-1, Finnish-1, Greek-1, Polish-1). We refer to this set as the *GuidanceSet*. Third, we run local indel-realignment for each sample individually, proceeding as follows:

1. We identified private target positions for the sample using *GATK RealignerTargetCreator* and providing the known indel sites from the 1000 Genomes project (as above). We then unified the identified targets with *TargetsBase* to obtain a sample-specific target set.
2. We run *GATK IndelRealigner* on this sample together with the *GuidanceSet* on the sample-specific targets. We parallelized this step additionally by contig, merging the output with picard-tools.
3. After local-realignment, we used *picard-tools FilterSamReads* to filter out reads that contained soft-clipped bases (identified with *ATLAS assessSoftClipping*) or did not pass picard-tools *ValidateSamFile*. We further used *SAMtools* to keep only primary alignments and, for paired-end libraries, proper pairs.

Step 3: Sample Stats

We determined read counts, sequencing depth, endogenous DNA-content and other statistics using *ATLAS BAMDiagnostics, ATLAS depthPerSiteDist* and *SAMtools flagstat*. These statistics are given per library-parallel for all ancient samples in Supp. Table 1 and merged for the 15 samples produced here in Table S3, Fig. S2**.**

**Table S3 - Sequencing data produced in this study** (n = 15).

| Sample | Total no. of sequenced reads | No. of reads mapping to hs37d5 after filtering | % of endogenous reads without duplicates | Mean sequencing depth (X) |
|---|---|---|---|---|
| AKT16 | 3,484,002,250 | 709,352,253 | 20.36027 | 12.25 |
| Asp6 | 1,139,245,581 | 689,412,495 | 52.80776 | 12.11 |
| Bar25 | 1,991,972,802 | 1,176,984,160 | 43.04312 | 12.65 |
| Dil16 | 1,164,789,725 | 652,044,979 | 48.95333 | 10.60 |
| Ess7 | 1,562,375,278 | 749,310,530 | 41.48627 | 12.34 |
| Herx | 2,356,604,701 | 698,260,906 | 25.51949 | 11.46 |
| Klein | 1,174,296,074 | 679,244,954 | 50.60461 | 11.30 |
| LEPE48 | 1,148,525,245 | 622,825,286 | 48.02362 | 10.92 |
| LEPE52 | 1,541,932,690 | 902,777,757 | 47.21246 | 12.37 |
| Nea2 | 1,168,563,125 | 723,747,890 | 52.05976 | 12.51 |
| Nea3 | 1,193,896,217 | 751,809,835 | 52.52959 | 11.57 |
| STAR1 | 1,167,689,785 | 673,467,996 | 49.30335 | 10.55 |
| VC3-2 | 1,537,245,748 | 765,893,269 | 42.79341 | 11.22 |
| VLASA32 | 1,603,082,543 | 899,098,270 | 46.05057 | 12.65 |
| VLASA7 | 1,601,828,507 | 989,180,954 | 53.10986 | 15.21 |

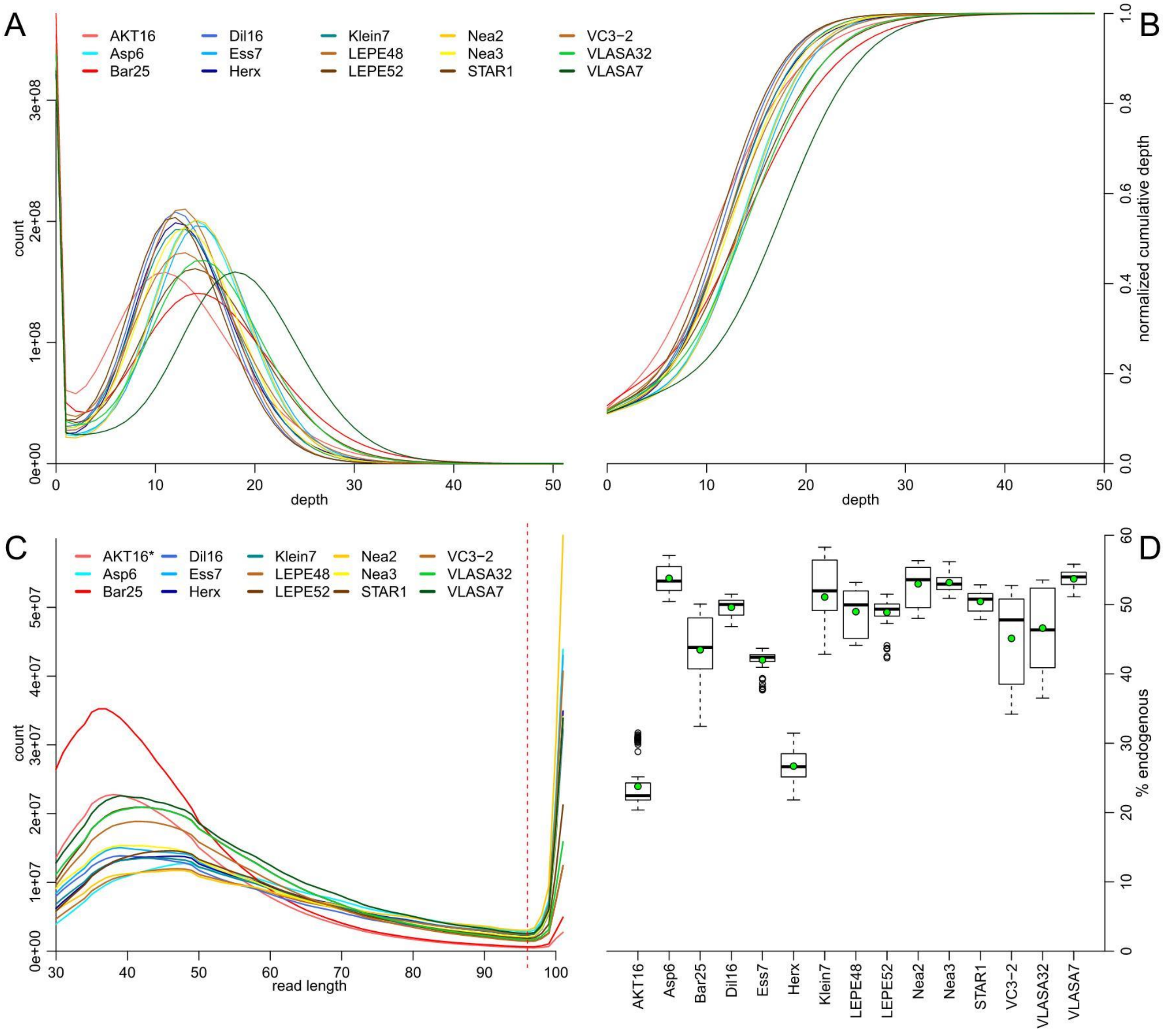


**Figure S2 - Sample Statistics.** (**A**) Distribution of read depth. (**B**) normalized cumulative depth distribution for all samples produced in this study. (**C**) Length distribution of sequenced single-end samples. The vertical dashed red line indicates where the samples were split for post-mortem damage estimation; reads above a length of 95 bp are assumed to not have been sequenced to their full length and may therefore miss the typical PMD-pattern on their 3' side. (*only single-end libraries were plotted for AKT16). (**D**) Percentage of endogenous reads among all sequencing reads per library-parallel (black) and for whole samples (green points) for all samples produced in this study.

Steps 4-7: ATLAS-Pipeline

We used commit 6df90e7 of the *snakemake* script *ATLAS-Pipeline* available at bitbucket.org/wegmannlab/atlas-pipeline with the following entries in the config-file:

- *sample_file:* Input-table with sample paths
  *atlas:* location of ATLAS executable
  *ref:* location of reference genome (hs37d5)
  *UCNE:* location of ultraconserved sites for recal
- *sequence: single, split=T, cutoff:5* for single-end libraries
  *sequence: single, split=F* for pre-merged reads from reference-data
  *sequence: paired* for paired-end libraries
- *recal: NO-file*
- *PMD: separate*

Step 4: Post-Mortem-Damage (PMD)

This step was performed with the *snakemake* script *ATLAS-Pipeline*.

We proceeded as follows to deal with post-mortem damage (PMD) affecting the ancient samples:

1. Since PMD is most prevalent at read ends, the PMD pattern is different if a read spans the entire fragment or not. Following (*23*), we therefore split single-end reads by length using ATLAS task=splitRGbyLength with the option *allowForLarger*. We identified reads that span the entire fragment as those shorter than the maximum read length minus 5 bases to account for variation introduced by adapter-trimming (Fig. S2).
2. We identified PMD patterns for each read group (PCR-parallel) using *ATLAS task=estimatePMD*, providing the reference genome with *fasta=hs37d5.fasta* and limiting the analysis to the chromosomes 1-22, X and Y with option *chr*.

As shown in Fig. S3, all samples show typical post-mortem damage patterns. VC3-2 contains libraries that were not UDG-treated and therefore shows higher PMD-patterns than the other samples from this study.

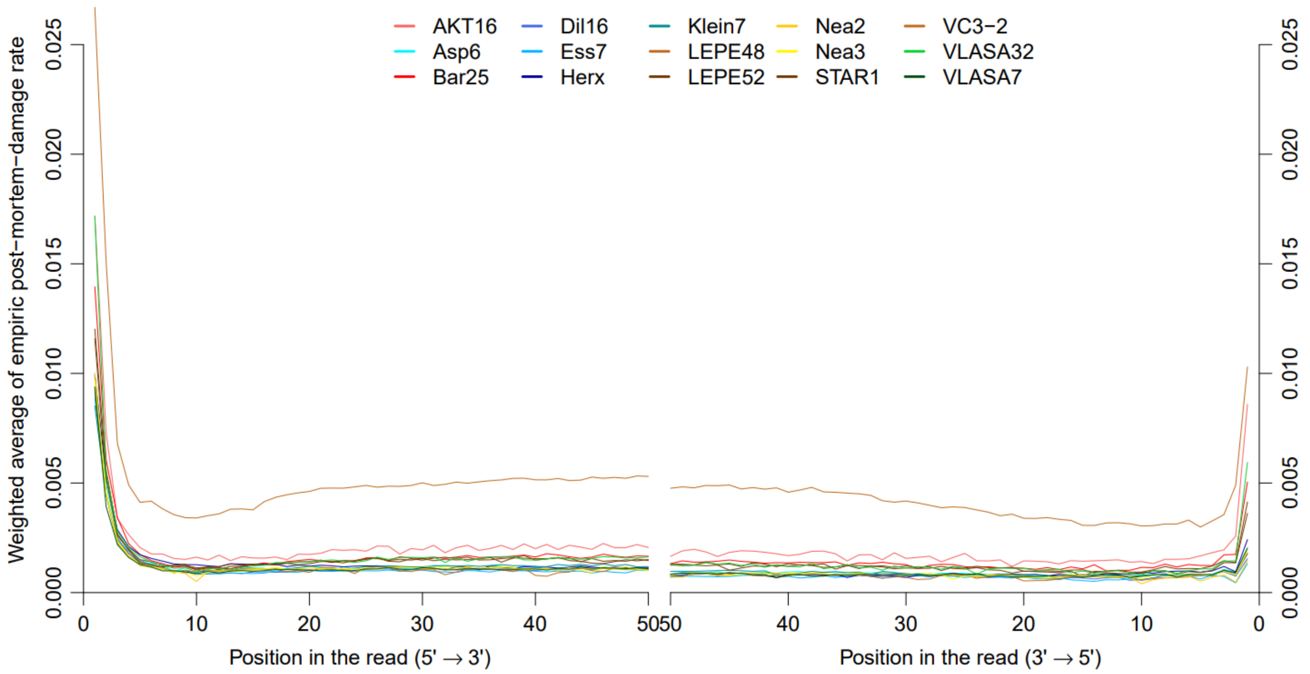


**Figure S3 - Estimated post mortem damage (PMD)** patterns for the first 50 bp (C→ T, left) and the last 50 bp (G → A, right). PMD was estimated per PCR-parallel. Shown here is the weighted average per sample. VC3-2 was not completely treated with UDG resulting in visibly higher PMD-patterns. All samples show a pattern of elevated PMD towards the end of the reads.

Step 5: Recalibration

This step was performed with the *snakemake* script *ATLAS-Pipeline*. We performed base-quality recalibration as described in (*23*) using *ATLAS task=recal* on known ultraconserved sites, obtained from https://ccg.epfl.ch/UCNEbase/data/download/ucnes/hg19_UCNE_coord.bed (*52*). We assumed equal base frequencies (*equalBaseFreq*) and provided the PMD estimates obtained above for each PCR-parallel (*pmdFile*). We inferred recalibration parameters for each PCR-parallel from all sites with a minimum depth of 2 (*minDepth=2*). Sequencing- and PCR-duplicates were not excluded from the estimation of recal parameters (*keepDuplicates*) since they add additional power. The quality transformation by base quality recalibration (created with *ATLAS task=qualityTransformation*) can be seen in Fig. S4.

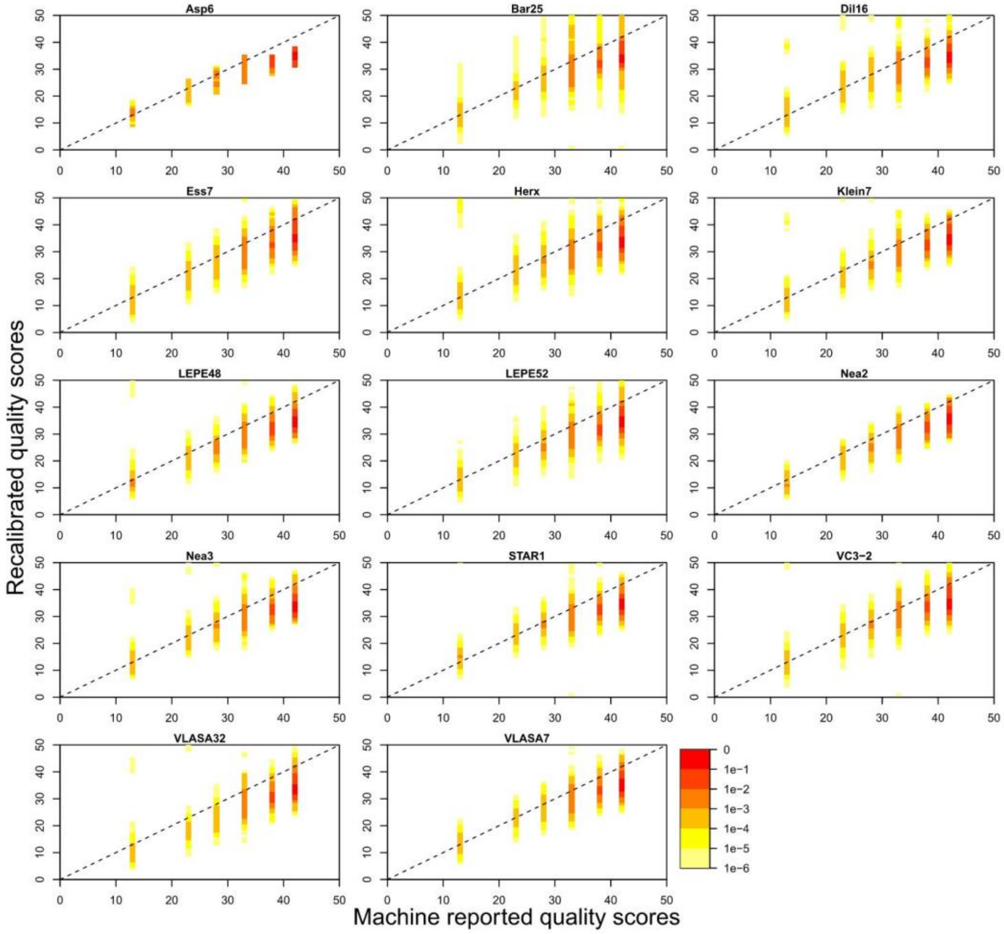


**Figure S4 - Quality Transformation for single-end genomes.** Base quality scores before and after recalibration. A general trend is visible, where the sequencing machine mostly overestimated the quality of the sequenced bases.

Step 6: Read-merging

This step was performed with the *snakemake* script *ATLAS-Pipeline*.

For paired-end samples and samples with partly paired-end sequenced libraries, the estimated recalibration parameters were directly corrected in the BAM file using *ATLAS recalBAM* providing the recal-parameters (*recal*) and the reference (*fasta=hs37d5.fasta*). Paired-end reads with corrected base qualities were then physically merged with *ATLAS task=mergeReads* to avoid the double-use of bases in the overlapping part. Reads that did not pass *picart-tools ValidateSamFile* were omitted. As a result of the physical merging, their read lengths changed, and we thus re-estimated PMD patterns as outlined above.

We also created recalibrated BAM files from single-end libraries, which we used in any subsequent analysis in which *ATLAS* was not involved (e.g. contamination estimation).

Step 7: Genotype calling

This step was performed with the *snakemake* script *ATLAS-Pipeline*.

We called Bayesian genotypes at all sites in the reference genome with *ATLAS task=callNew* and parameters *method=Bayesian*, *prior=theta*, *fixedTheta=0.001*, *infoFields=DP*, *equalBaseFreq*. The reference genome, PMD patterns and recal-patterns (for single-end read groups) were provided to the caller with *fasta*, *pmdFile and recal*, respectively. The first and last 2 bp of each read were ignored (*trim5=2*, *trim3=2*).

Step 8: Molecular sexing

The molecular sex (Table 1) was inferred using a script from (*53*) version 4.0, based on the ratio of reads aligning to the X and Y chromosomes (Fig. S5).

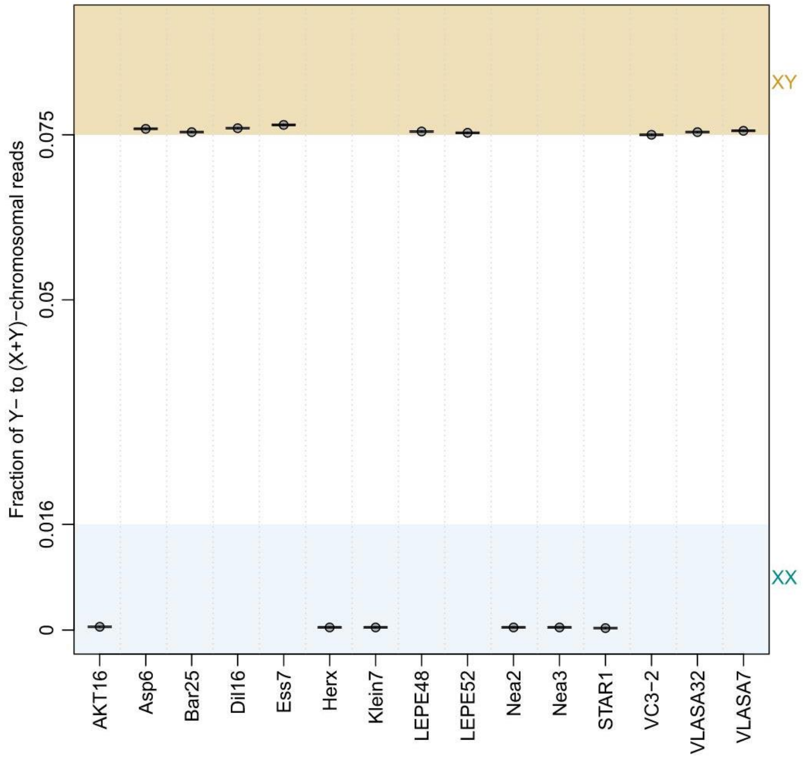


**Figure S5 - Inferred chromosomal sex for the 15 newly sequenced genomes,** based on the share of reads aligning to Y-chromosome from the reads aligning to X and Y chromosomes. A ratio below 0.016 indicates an XX genotype, a ratio above 0.075 indicates an XY genotype. Note that error bars showing 95% confidence intervals are too small to be visible in this plot.

Step 9: Contamination estimation

Blank controls of extraction, library-preparation and index-PCR were analyzed alongside the screening process. The concentration of potential contaminants was never higher than 0.81 ng/µl. A Bioanalyzer analysis (HS DNA, Agilent Technologies, Waldbronn, Germany) and the screening results for extraction- and library-controls confirm the detected DNA to consist of primer- and adapter-dimers with a maximum of 55 aligning reads per blank control (out of a potential share of 200,000 reads ; see Supp. Table 1).

To estimate modern human contamination in the sequencing results, two standard approaches were used:

**ContamMix:** The *ContamMix* R-script *estimate.R* estimates the amount of authentically mapping mitochondrial reads at 311 modern diagnostic marker positions. We run ContamMix with the option --*baseq20* on the following two input-files that we generated from all reads in the BAM file that mapped against the MT-Genome (*SAMtools*):

1. An alignment of mitochondrial reads against their own consensus (*-samFn*). This consensus was constructed by extracting iteratively mapping the reads with *MIA* (*mia*) and mapping the last iteration with *MIA* (*ma*) to obtain one FASTA-sequence. The MT-reads were mapped against their consensus using *bwa aln* and *samse (V.0.7.17)* and filtered for a minimum mapping quality of 30 with *SAMtools*.
2. A multiple alignment of the consensus genome and a fasta-file containing the diagnostic marker positions against each other (*--malnFn*) obtained with *mafft*.

**ANGSD:** For male individuals, contamination was additionally estimated using the *ANGSD contamination* script based on haploid X-chromosomal regions (*X:5000000-154900000*). We run *ANGSD* with the options *-doCounts 1* and *-iCounts 1*, followed by the executable *misc/contamination*, providing the publicly available *HapMap* file *HapMapChrx.gz*.

The contamination estimates obtained with both methods, the percentage of not authentically mapping reads by *ContamMix* and the estimated contamination rate by *ANGSD*, are shown in Fig. S6 and Supp. Table 1. While the two methods use complementary approaches (i.e. comparing mtGenomes at Neanderthal-specific sites vs heterozygosity on X-chromosomes in males), resulting contamination estimates are generally low (<2% in most cases, <4% in all cases).

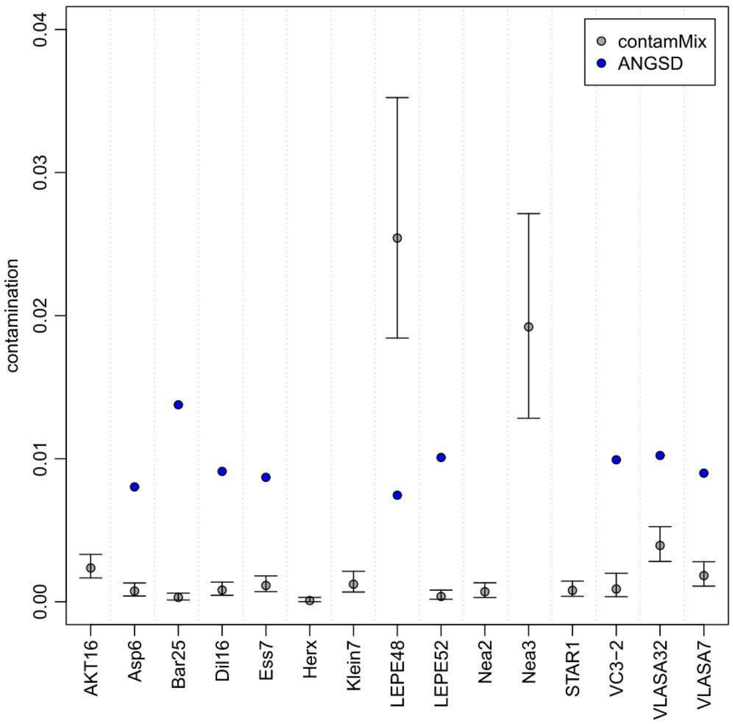


**Figure S6 - Percentage of contamination in the 15 newly sequenced samples.** All samples show a contamination rate below 4%. Grey: ContamMix, 1-mapAuthentic, error bars represent 95% CI. Blue: ANGSD (male individuals only), Method1, new_llh, ML, standard error for all ANGSD analysis is below 2e-12 and therefore not visible in this plot.

**Reference samples**

To minimize reference biases (*54*), we committed to analyze all previously published samples with the same pipeline outlined above. However, some adaptations were necessary for some samples:

- Blank characters in read names (e.g. from ENA accession numbers) were removed to ensure the proper detection of optical duplicates.
- The quality scores of the Bichon FASTQ files were converted from illumina 1.5 to illumina 1.9 with *setqk seq -V 64*.
- Stuttgart and Loschbour were received as unaligned and untrimmed demultiplexed raw data in BAM file format and transformed to unaligned FASTQ format using *picard-tools SamToFastq*.
- For the Loschbour files, individual library information was not recoverable. Therefore duplicates across all sequencing files were marked in this step of the pipeline.
- Raw sequencing data was not obtainable for Bon002 and SF12. For Bon002, we downloaded the published BAM files converted back into FASTQ files from ENA (run accession ERR1514027). For SF12, we used published BAM files with already physically merged reads and converted them back to FASTQ files using *picard-tools SamToFastq*. For both, we split the FASTQ files

by run identifiers to treat each run separately throughout the pipeline. Since those reads were already pre-processed, we omitted adapter trimming, removed orphaned reads and otherwise followed the pipeline for single-end samples without splitting read groups by length.

- For NE1, one of the FASTQ files was corrupted and only 25% of the contained reads could be recovered. As a result, we could only use 97% of the total number of reads generated for that sample (*38*). We then demultiplexed the fixed FASTQ-files with a custom bash script, allowing one mismatch at the first position and one mismatch at any other position of the index.
- Since the modern samples from the SGDP dataset were already aligned to the desired reference genome, we abstained from remapping them and analyzed them along the ancient samples, starting with marking optical and PCR duplicates (*picard-tools*) and local realignment.

For each ancient sample, all differences from the standard pipeline are detailed in Supp. Table 1.

### ***<u>Data sets</u>***

#### **Data filtering**

Individual VCFs obtained after Bayesian genotype calling (Step 7 of the Bioinformatics pipeline) were first filtered according to their read depth (DP): for each individual we excluded sites that had DP < 8 and those that had a DP bigger or smaller than 2.5 s.d. away from the mean DP for each individual. To minimize effects of outliers, the mean DP was calculated using the R function *optimize* for the sites comprised +/-10 of the mode DP and a tolerance of $10^{-6}$ (see Supp. Table 2). We also excluded sites that had poor genotype quality (GQ < 30), and we only kept autosomal sites. Furthermore, heterozygous sites were considered as homozygous if they had a significant allelic imbalance (*p-value* ≤ 0.1) tested by Fisher's exact test. After filtering, the modern individuals had on average 2,549,812,798 sites passing the filters against 1,980,736,974 for the ancients (ranging from 333,061,590 to 2,544,170,685). All VCF file processing was performed using *bcftools* v1.9.

Applying these filters led to a substantial increase in the quality of our heterozygous sites, reducing the amount of sites with high allelic imbalances and decreasing the asymmetry between singleton and non-singleton sites to a minimum (Fig. S7). To quantify this improvement explicitly, let us denote by $H_{is}$the set of loci at which individual *i* was called heterozygous either for a singleton (only non-reference allele across all samples, s = 1) or not (s = 0). We then define the imbalance statistics

$$r_{is} = \frac{\sum_{l \in H_{is}} \quad I(\frac{r_{il}}{d_{il}} > 0.5)}{\sum_{l \in H_{is}} \quad I(\frac{r_{il}}{d_{il}} < 0.5)},$$

where $r_{il}$ denotes the number of reference alleles observed in individual *i* at site *l* out of the total number of observed alleles $d_{il}$ at this site. To compare singleton (s = 1) against non-singleton (s = 0) positions, we further quantify $\rho = r_{i1} / r_{i0}$. As shown in Fig. S8, the chosen filters guarantee comparable imbalances at singleton and non-singleton sites, indicating similar quality of both categories. A striking outlier was sample CarsPas1, which was not included in any demographic analysis.

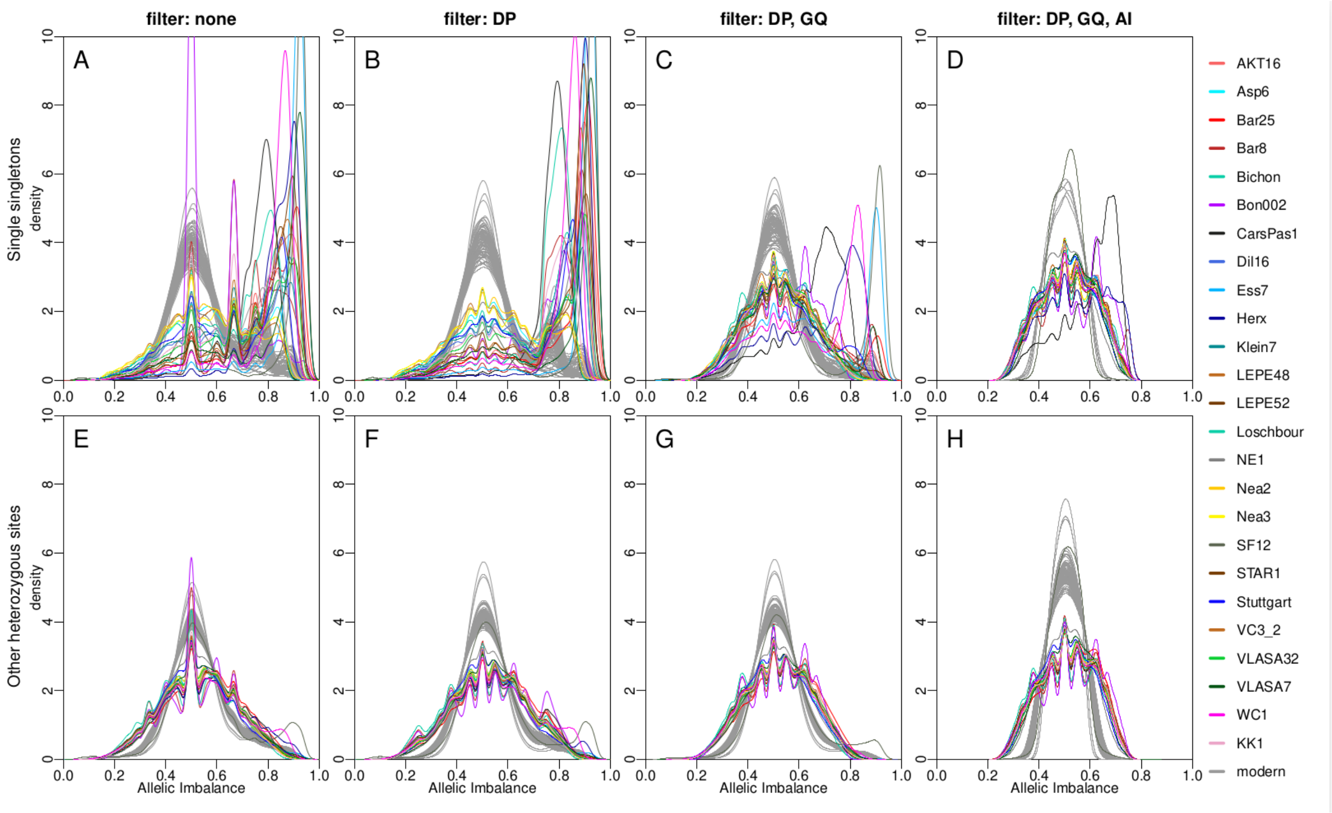


**Figure S7 - Successive effects of filtering on allelic imbalance.** Density of the allelic imbalance for singleton positions (top line panel) and other heterozygous positions (bottom line panel). (**A**,**E**) No filter; (**B**,**F**) Depth filter (DP) as described in the text; (**C**,**G**) DP filter and genotype filter (GQ ≤ 30); (**D**,**H**) DP, GQ and allelic imbalance filter (AI > 0.1).

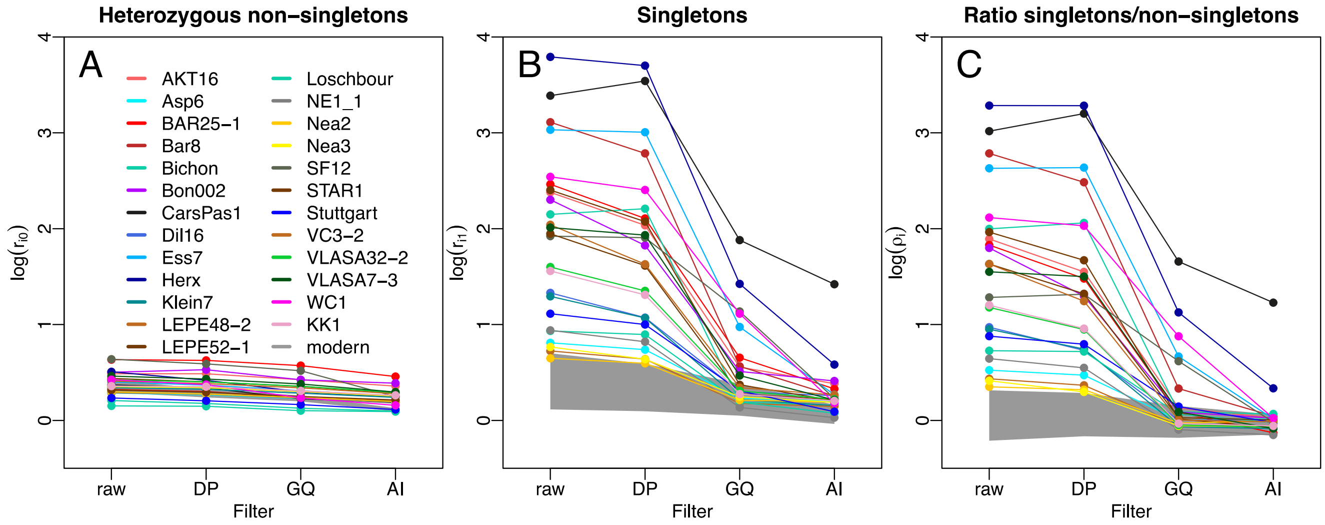


**Figure S8 - Imbalance statistics.** The imbalance statistics (**A**) $r_{i0}$ calculated for non-singleton sites and (**B**) $r_{i1}$ calculated for non-singleton sites. (**C**) The imbalance ratio $\rho_i = r_{i1}/r_{i0}$. The range of these statistics observed for modern samples is shown as grey shades in all panels.

**Dataset assembling**

The filtered individual VCFs were then merged into single VCFs, one only including the 25 ancient genomes ("*Ancient*" dataset) and one including all the 102 ancient and modern samples ("*102samples*" dataset). The data sets were then polarised with the Chimpanzee reference genome (*panTro4*), and we only kept the biallelic polymorphic sites (6,986,216 in the *Ancient* dataset; 14,896,103 in the *102samples* dataset). Note that individuals can have missing genotypes in these data sets (on average, the modern genomes show 4% of missing sites with a maximum of 6%, the ancients 29% ranging from 3 to 87% in the *102samples* dataset).

**Neutral portion of the genome**

We performed most of our genetic analyses on a "neutral" portion of the genome (*55*) in order to avoid biases due to background selection (BGS) and biased gene conversion (BGC) when estimating population diversity and relationships (*56*). We defined this portion as the a restricted set of sites that had the same reference allele in the chimpanzee and gorilla reference genomes, that were in regions with recombination rate >1 cM/Mb where BGS has little effect, outside of CpG islands, non CpG sites (i.e. when a C is followed by a G in the chimpanzee and gorilla genomes and has a T as alternative allele in our data; or a G preceded by a C in in the chimpanzee and gorilla genomes with an A as alternative allele) and with A↔T and G↔C mutations which are not affected by BGC or post-mortem damages. We obtained the Chimpanzee and the Gorilla hg19 nucleotide states from the chained and netted alignments http://hgdownload.cse.ucsc.edu/goldenpath/hg19/vsPanTro4/axtNet/ and http://hgdownload.cse.ucsc.edu/goldenpath/hg19/vsGorGor5/axtNet/ respectively; the recombination map for the YRI population from the 1000G project (ftp://ftp.1000genomes.ebi.ac.uk/vol1/ftp/technical/working/20130507_omni_recombination_rates/); the CpG islands listing from the UCSC genome browser (CpG Islands (cpgIslandExt) Track).

For the *fastsimcoal2 (57, 58)* demographic analyses, we prepared six panels from the *Ancient* dataset (Table S4). For each panel, we excluded sites with any missing data in any individual of the panel as well as sites with different reference allele for the chimp and gorilla reference genomes, CpG sites, sites found in CpG islands or in genomic regions with recombination rate less than 1 cM/Mb. We thus kept a total of $T_X$ sites for the panel$_X$, among which $S_X$ were polymorphic and $M_X$ monomorphic (the properties of each panel are found in Table S5). Second, in order to have neutral data sets best suited for demographic inference (*55*), we only kept $S_{neutral\_X}$ BGC-free A↔T and G↔C polymorphic sites, which represented $\alpha_X$~ 0.2 of the filtered polymorphic sites for any panel. Since we would have expected them to represent one third of all filtered sites if mutation rates had been equal for all mutation types, we computed the ratio $r_X = \alpha_X/(1/3) = 3\ \alpha_X$, which represents the reduction in mutation rates that has occurred at those sites in panel$_X$. We then estimated the number of neutrally evolving monomorphic sites in each dataset as $M_{neutral\_X} = M_X/3$, since we expect that one third of all mutations should be BGC-free.

**Phasing and imputation**

For the Runs of Homozygosity and MSMC analyses, we did genotype imputation and phasing for each chromosome on the *102samples* dataset that was polarised back with *hg37d5* human reference genome. We used *SHAPEIT4* v1.2 (*59*) with by-default parameters for sequencing data, the HapMap phase II b37 genetic map, and the Haplotype Reference Consortium (*60*) dataset (accession number EGAD00001002729 on the European Genome-phenome Archive) as reference panel. All the chromosomal phased-imputed VCFs were then concatenated into a single phased VCF with *bcftools* v1.9, which includes 2,795,127,079 sites in total.

**Table S4 - Population composition and ages used for demographic modeling.** Populations that are archeologically defined as Palaeolithic or Mesolithic are highlighted in light grey; the other populations belong to Neolithic cultures. The age of the population, given in generations with a generation time of 29 years (*61*), was taken within the range of the ages of its constituent samples.

| **Population** | **Age BP** | **Genomes** | **Panels** | | | | | |
|---|---|---|---|---|---|---|---|---|
| | | | *HGNeo* | *Aegeans* | *AKT* | *Bon* | *NeoEur* | *HG* |
| Konya Plain | 347 | Bon002 | | | | X | | |
| Zagros region | 317 | WC1 | X | X | X | X | | X |
| NW Anatolia (AKT) | 295 | AKT16 | | | X | | | |
| NW Anatolia (BAR) | 295 | Bar25 | | X | X | | | |
| Northern Greece | 280 | Nea2; Nea3 | X | X | X | X | X | |
| Central Serbia | 260 | STAR1; VC3-2 | | | | | X | |
| Lower Austria | 245 | Klein7; Asp6 | | | | | X | |
| Southern Germany | 240 | Dil16; Ess7; Herx | | | | | X | |
| Caucasus | 335 | KK1 | X | X | X | | X | X |
| Danube Gorges | 295 | VLASA7; VLASA32 | X | X | X | X | X | X |
| WHG (Bichon) | 472 | Bichon | | | | | | X |
| WHG (Loschbour) | 278 | Loschbour | | | | | | X |

**Table S5 - Properties of the different panels used for the demographic inferences.**

| **Label**[a] | **Description** | ***HGNeo*** | ***Aegean*** | ***AKT*** | ***Bon*** | ***NeoEur*** | ***HG*** |
|---|---|---|---|---|---|---|---|
| $T_X$ | Total number of sites passing filtering criteria[b] | 262,581,434 | 243,505,860 | 200,141,597 | 35,751,107 | 175,791,292 | 167,849,230 |
| $S_X$ | Polymorphic among $T_X$ | 436,966 | 410,130 | 331,510 | 56,823 | 309,817 | 256,668 |
| $S_{neutral\_X}$ | WWSS $S_X$ sites | 86,557 | 81,310 | 65,534 | 12,150 | 60,761 | 52,025 |
| $M_X$ | Monomorphic among $T_X$ | 262,144,468 | 243,095,730 | 199,810,087 | 35,694,284 | 175,481,475 | 167,592,562 |
| $M_{neutral\_X}$ | $M_X/3$ | 87,381,489 | 81,031,910 | 66,603,362 | 11,898,095 | 58,493,825 | 55,864,187 |
| $T_{neutral\_X}$ | $M_{neutral\_X}+S_{neutral\_X}$ | 87,468,046 | 81,113,220 | 66,668,896 | 11,910,245 | 58,554,586 | 55,916,212 |
| $\alpha_X$ | $S_{neutral\_X}/S_X = r_X/3$ | 0.1981 | 0.1982 | 0.1977 | 0.2138 | 0.1961 | 0.2027 |
| $\mu_{neutral\_X}$ | $r_X{*}\mu tot = r_X{*}1.25e{-}8$ | 7.43E-09 | 7.43E-09 | 7.41E-09 | 8.02E-09 | 7.35E-09 | 7.60E-09 |

[a] X indicates a given panel.

[b] All sites found without missing data for all individuals of a given dataset, that are non CpG sites nor in CpG islands, with similar reference allele in the chimpanzee and gorilla reference genomes and a recombination rate ≥ 1 cM/Mb

***Uniparental haplogroup determination***

We used phy-mer (*62*) to determine mitochondrial haplogroups from BAM files for the 15 newly sequenced genomes, with the minimal number of occurrences of a K-mer set to 10 (Supp. Table 3). For the samples genetically identified as men, Y-chromosomal haplogroups were determined from BAM files using Yleaf (*63*), using the recommended minimal base-quality of 20 (-q 20) and base-majority to determine an allele of 90% (-b 90).

***Phenotype predictions***

**Pigmentation**

Pigmentation phenotypes of hair, skin and eyes were predicted for each of the newly sequenced samples with the HIrisPlexS webtool (*64*, *65*) from the *Ancient* dataset. In case of missing data for any of the 41 SNPs used by HirisPlexS, *i.e.* sites with low or no depth, the positions were looked up in the BAM files directly (see Table S9). In BAMs, we only considered sites at least 3 bases away from either end of the read, with a base quality ≥ 25, and no C↔T and G↔A SNPs to avoid any effect of PMD on the prediction. In order to deal with the uncertainty associated to observing alleles in BAM files directly, two HirisPlex input files were created for each individual: one in which all sites missing from the VCF and meeting the criteria above were assumed to be homozygous for the allele found in the BAM, and another in which all positions were assumed to be heterozygous. Running HirisPlexS twice for each sample resulted in ranges of probabilities for each phenotype (Table S9). A prediction was accepted without further explanation if in both runs the same phenotype showed a probability ≥ 0.7 (*65*). If predictions differed between runs, the most parsimonious phenotype was chosen, following the approach of Walsh in (*42*).

**Standing height**

To predict standing height, a classical highly heritable polygenic trait (*66*), polygenic scores (PS) were computed based on a set of 670 SNPs (*67*) on the *Ancient* dataset. To account for missing data common even in high depth aDNA samples, we applied the generalized risk score approach described by (*68*) on samples where enough SNPs were present to account for at least 75% of the scores effect size. This led to the exclusion of five ancient individuals (Bar8, Bon002, Bichon, CarsPas1, AKT16) over the 25 included in this study.

**Other phenotypes**

Genotypes for SNPs associated with additional phenotypes of interest were inspected manually for each sample in the VCFs or BAM files if necessary: rs4988235 variant in *MCM6* gene and associated with lactase-persistence in Eurasia; rs3827760 in *EDAR* gene; rs17822931 in *ABCC11*; seven SNPs located in the *FADS1/2* gene complex.

### ***Population genetics***

**Genomic heterozygosity**

We estimated the level of genetic diversity of each individual as the proportion of heterozygous sites found in the neutrally evolving portion of its genome. Thus, for every genome, we divided the amount of neutral heterozygous sites observed in the *102samples* dataset by the expected number of neutral sites genotyped for this individual. This number is obtained by considering all the genotyped sites for the considered individual that i) have the same reference allele for both the chimpanzee and gorilla reference genomes, ii) are not CpG sites and out of CpG islands, iii) are in regions with recombination rate > 1 cM/Mb based on (*55*) (Details are given above in the "Data sets" section). Finally, in order to consider only sites unaffected by BGC, we divided the number of these previously defined sites by 3, as only one third of all mutations should be BGC free (i.e. A↔T and G↔C polymorphisms).

**Multi-Dimensional Scaling (MDS) on pairwise average nucleotide divergence**

Genetic relationships among individuals were estimated from pairwise average nucleotide divergence $\pi_{XY}$ *(69)*. For each pair of individuals X and Y, we identified the sites showing no missing data for the two individuals, and we computed their average nucleotide divergence $\pi_{XY}$ over these sites. We considered whole-genome sites or neutral sites only (in this case, the number of differences was divided by the expected number of neutral sites as defined above in the heterozygosity paragraph). We then represented the relationships between modern and ancient genomes from Europe and SW Asia (for sites from the *102samples* dataset; Fig.2A and Fig. S19A) or only the ancient genomes (for sites from the *Ancient* dataset; Fig. S19B), using a classical multidimensional scaling (MDS) approach implemented in R (*cmdscale* function).

**Runs of Homozygosity (ROHs)**

We inferred the segments of homozygosity-by-descent or Runs of Homozygosity (ROHs) from the imputed genomes of the 90 European and SW Asian modern and ancient individuals by using IBDSeq v. r1206 (*70*) with default parameters but *errormax = 0.005* and *ibdlod = 2*. We further processed the artificially long tracts spanning assembly gaps or centromeres (their genomic locations were obtained from UCSC genome browser: http://hgdownload.cse.ucsc.edu/goldenPath/hg19/database/gap.txt.gz) and split them into shorter tracks excluding the gap stretch, inspired by (*71*). As advised when genomes do not come from an homogeneous population (*70*), we focused on short (2-10 Mb) and long (>10 Mb) ROHs similarly to (*72*).

**Admixture clustering analyses**

Admixture coefficients of each ancient genome were estimated using the R package *LEA (73)* with parameters K = 2 or 3, alpha = 10, and number of repetitions = 5. We used the function *snmf* to calculate the fit (entropy) of each run and we plotted the admixture coefficients from the run with the smallest entropy value. The groups were defined in an unsupervised manner. In order to maximize the number of genomic sites to be used, we excluded for these analyses the three lowest quality Neolithic genomes (Bichon, Bon002, Bar8).

**Neanderthal introgression**

We quantified proportions of Vindija Neanderthal (*74*) introgression by computing $f_4$ ratio statistics of the form $\frac{f_4(Altai,Chimp; X,Dinka)}{f_4(Altai,Chimp; Vindija,Dinka)}$ as previously suggested (*75*) with *qpF4ratio* from the ADMIXTOOLS package (*18*) on diploid calls of the *HOIll* subset of the 1240k capture sites (*9*). Altai, chimp, Vindija and Dinka were retrieved from David Reich's lab webpage (https://reich.hms.harvard.edu/downloadable-genotypes-present-day-and-ancient-dna-data-compiled-published-papers, version 37.2).

**f-statistics**

Comparison to target-enriched data sets

We compared the samples obtained in this study with previously published, target-enriched datasets available through the reference 1240K dataset v42.4 at https://reich.hms.harvard.edu/downloadable-genotypes-present-day-and-ancient-dna-data-compiled-published-papers. We refer to this dataset as 1240K in the following. To ensure our data is comparable to the pseudo-haploid calls in 1240K, we generated majority calls for all our ancient individuals with *ATLAS* (task=majorityBase; commit 7cfc900) at the 1,233,013 SNP positions present in this reference dataset.

Confirming population assignments

We first used *f*-statistics to confirm that the assignment of samples to specific populations is not violated by the presence of shared drift with external samples. For this, we calculated all possible *f*-statistics of the form of *D(Individual 1 from the population tested, Individual 2 from the population tested; Other samples, Outgroup)* using ADMIXTOOLS (*18*) and *Mbuti* as the outgroup. We used *qpGraph* using 1,000 initial conditions to match specificities of the datasets.

Validating the *fastsimcoal2* model

To validate the model inferred by *fastsimcoal2,* we aimed at comparing *f*-statistics predicted under the *fastsimcoal2* model against those calculated from the data. For this, we first created a full admixture graph carefully matching the model in Fig. 3A. To validate this graph, we used data simulated with *fastsimcoal2* under the model shown in Fig. 3A (see section Final Model), which we fitted with *qpGraph* from ADMIXTOOLS v7300 (*18*) 100 times with 1,000 initial conditions each. However, *qpGraph* failed to identify a suitable set of branch lengths and admixture proportion to explain the simulated data with all runs resulting in predicted *f*-statistics very different from those calculated on the simulated data, suggesting that the SFS captures more information of the full data than *f*-statistics do.
Since the full graph could not be fitted, we next aimed at manually simplifying the graph. Specifically, we removed admixture edges, until we identified the simplest graph (in terms of admixture edges) for which no major differences between the *f*-statistics predicted under the model and those calculated from the simulated data were detected. This graph was then re-fitted with *qpGraph* on both the neutral set of sites used in our *fastsimcoal2* analysis as well as on the 1024K data set described above. In both cases, *qpGraph* was run 100 times with 1,000 initial conditions each. To present the fitted graphs and quantify the reliability of the fitting, we calculated the median and 90% quantiles for all branch lengths and admixture odds ratios across the 20 best graphs as judged by the final score.

**Joint Distribution of Fitness Effects (DFE) analysis**

To create a representative Neolithic model population, we aggregated all Neolithic samples that were newly sequenced, plus the WC1 genome (N = 14). To create a representative modern population, we used SGDP populations in rough proximity to the ancient populations (Polish, Bergamo, Czech, French,

Hungarian, Greek, Albanian, Bulgarian, and Turkish). We annotated synonymous and nonsynonymous SNPs from the *102samples* dataset using ANNOVAR (*76*) with the hg19/GRCh37 as the reference genome. Ancestral alleles were determined based on the Ensembl Compara 71 genome FASTA files. To account for missing data, we projected the joint allele frequency spectra downward to 16 Neolithic chromosomes and 28 modern chromosomes, to roughly maximize the number of segregating synonymous SNPs.

The joint DFE analysis requires a simpler demographic model than our main inference (Fig. 3A), and simulations suggest that joint DFE analysis is robust to demographic model details (*77*). Based on prior results (*78*), we fit a demographic model to the synonymous data in which the ancestors of the Neolithic population underwent a bottleneck to relative size $n_B$ followed by exponential growth (Fig. S24A) The ancestors of the modern samples diverged $T_B$ time units after the bottleneck, following the same growth rate to reach final size $n_F$. $T_S$ time units after this divergence, Neolithic chromosomes were sampled, $T_F$ time units before present. We also included a parameter $p_{mis}$ to account for potential ancestral state misidentification (*79*). To fit this model, we used grid points of [128, 138, 148], and the best-fit parameters were $n_B = 0.113$, $n_F = 1.42$, $T_B = 0.106$, $T_S = 0.0054$, $T_F = 0.0232$, $p_{mis} = 0.0634$.
We then fit a model including this demographic history plus a joint distribution of fitness effects (DFE) to the nonsynonymous data. We modeled the DFE as a bivariate lognormal distribution. We assumed that the population-scaled mutation rate $q$ for nonsynonymous mutations was 2.31 times that for synonymous mutations, and we again included a parameter for ancestral state misidentification. We assumed that selection coefficients were potentially different in the ancestors of the modern individuals (Fig. S24E). The best fit parameters for the joint DFE were $m = 3.29$, $s = 2.61$, and $r = 0.9968$, with 0 misidentification inferred.

### ***Demographic Analyses***

#### **MSMC2 analyses**

Data processing

We used *MSMC2 (80)* to infer past effective sizes of ancestral populations and their split times for all high-quality ancient and some representative modern individuals (Table S6) on the phased-imputed *102samples* dataset prepared with *SHAPEIT4*. To ensure high data quality especially high mappability and genotype quality, we followed (*80*) and used two masks: 1) per chromosome mappability masks (um75-hs37d5.bed.gz) for the human reference genome hs37d5 downloaded from *https://github.com/wangke16/MSMC-IM/tree/master/masks*; 2) sample-specific masks that we generated as suggested by (*80*). We then ran the "generate_multihetsep.py" script from MSMC-tools (*https://github.com/stschiff/msmc-tools, commit 07bc8a9*) to get single, multi-sample and pair population input files for *MSMC2*.

Example of a command line for two, four or eight haplotypes:

```
generate_multihetsep.py --chr 1 --mask ind1.chr1.mask.bed --mask mappabilityMaskperChr/chr1_m75-hs37d5.bed ind1.chr1.phased.vcf > ind1.chr1.multihetsep.txt
generate_multihetsep.py --chr 1 --mask ind1.chr1.mask.bed --mask ind2.chr1.mask.bed --mask mappabilityMaskperChr/chr1_m75-hs37d5.bed ind1.chr1.phased.vcf ind2.chr1.phased.vcf > pop1.chr1.multihetsep.txt
generate_multihetsep.py --chr 1 --mask ind1.chr1.mask.bed --mask ind2.chr1.mask.bed --mask ind3.chr1.mask.bed --mask ind4.chr1.mask.bed --mask mappabilityMaskperChr/chr1_m75-hs37d5.bed ind1.chr1.phased.vcf ind2.1.phased.vcf ind3.chr1.phased.vcf ind4.chr1.phased.vcf > pop1_pop2.chr1.multihetsep.txt
```

**Table S6 - Samples considered in the MSMC2 analyses** (n = 30).

| Population | Period | Samples |
|---|---|---|
| Africa | Modern | Mende-1, Mende-2 |
| Western Europe | Modern | French-1, French-2 |
| Eastern Asia | Modern | Han-1, Han-2 |
| Southern America | Modern | Karitiana-1, Karitiana-2 |
| Central Serbia | Neolithic | STAR1, VC3-2 |
| NW Anatolia | Neolithic | AKT16, Bar25 |
| Hungary-Neo | Neolithic | NE1 |
| Lower Austria | Neolithic | Asp6, Klein7 |
| Northern Greece | Neolithic | Nea2, Nea3 |
| Southern Germany1 | Neolithic | Dil16, Ess7 |
| Southern Germany2 | Neolithic | Herx, Stuttgart |
| Zagros Region | Neolithic | WC1 |
| Danube Gorges-Meso | Mesolithic | VLASA7, VLASA32 |
| Northern Europe-Meso | Mesolithic | SF12 |
| Caucasus | Late Mesolithic | KK1 |
| Western Europe-Meso | Upper Palaeolithic - Mesolithic | Bichon, Loschbour |
| Lepenski Vir | Transformational - Neolithic | LEPE48, LEPE52 |

Inference of human population size

We inferred past effective population size for each diploid individual separately (Fig. S25A) as well as using two samples per population (Fig. S25B), if available (see Table S6), using command lines such as:

```
msmc2 -t11 -s -o ind1.2haps.msmc2 ind1.chr*.multihetsep.txt
msmc2 -t11 -I 0,1,2,3 -s -o pop1.4haps.msmc2 pop1.chr*.multihetsep.txt
```

All *MSMC2* results were scaled using a mutation rate of $1.25 \times 10^{-8}$ per base pair per generation (*80*, *81*), and a generation time of 29 years (*61*).

Divergence time between populations

To estimate split times between population pairs, we used *MSMC2* 1) to estimate coalescent rates among the samples of the first population, 2) to estimate coalescent rates among the samples of the second population, and 3) to estimate coalescent rates across the two populations. Example command lines used for two populations pop1 and pop2 were:

*msmc2 -t11 -I 0,1,2,3 -s -o pop1.4haps.msmc2 pops.chr*.multihetsep.txt,*
*msmc2 -t11 -I 4,5,6,7 -s -o pop2.4haps.msmc2 pops.chr*.multihetsep.txt,*
*msmc2 -t11 -I 0-4,0-5,0-6,0-7,1-4,1-5,1-6,1-7,2-4,2-5,2-6,2-7,3-4,3-5,3-6,3-7 -s -o pop1-pop2.8haps.cross.msmc2 pops.chr*.multihetsep.txt.*

We then used the *MSMC2* script *combineCrossCoal.py* to create a single output file with all three rates:

*combineCrossCoal.py pop1-pop2.*.cross.msmc2.final.txt pop1.*.msmc2.final.txt pop2.*.msmc2.final.txt > pop1-pop2.combined.msmc2.final.txt*

For each population-pair, we then plotted the relative cross-coalescence rate (CCR), which is estimated by taking the ratio of the across-rate and the mean within-rate (Fig. S26) (*80*). The relative CCR indicates when two populations were a single population (values around 1) and when they were well separated into two isolated populations (values close to zero) (*80*). However, translating the relative CCR into estimates of split times is difficult for two reasons: First, if a population split was followed by migration, the relative CCR will remain high even after the split. Second, the uncertainty associated with the different colaescent rates translates into a gradual change in the relative CCR, even under a hard split. As a rough estimate, (*80*) recommends estimating split times as the time when the relative CCR hits 0.5, which we show in Fig. S27A-B for all pairwise comparisons.

Bootstrap analysis for the $N_e$ and CCR estimates

To obtain confidence intervals around coalescence rate estimates, we generated 20 artificial genomes by block-bootstrapping *MSMC2* input files in 5Mb blocks as suggested in (*80*).

### ***fastsimcoal2*** **analyses**

The analyses were carried out on six different panels of newly sequenced individuals (Table S4), on the neutral SFS and with neutral mutation rate adjusted for each panel from the basal mutation rate to take into account the potentially lower rate of A↔T and G↔C mutations (see Table S5).

Parameter inference via maximum likelihood

Parameter estimates were obtained by maximizing the model likelihood over 50 independent runs of *fastsimcoal2 (58)*, 100 expectation conditional maximization (ECM) cycles per run and 500,000 coalescent simulations per estimation of the expected SFS (except for the *HGNeo* models for which 200,000 simulations were run). The command line used for the estimation was of the type:

*fsc -t xxx.tpl -n500000 -d -e xxx.est -M -l100 -L100 -q -C5 --multiSFS --logprecision 18 -c1 -B1* where fsc is the *fastsimcoal2.7* program (available on http://cmpg.unibe.ch/software/fastsimcoal2/) and *xxx* the generic name of the input files. The .est and .tpl input files used for inference under the best model will be made available upon request.

Likelihoods comparison and model choice

When several models were tested for the same dataset, we retained the model with the highest estimated likelihood over 50 runs, and recorded the maximum likelihood (ML) parameters of this model. In order to take into account the variance in the estimation of the likelihoods due the limited number of performed coalescent simulations (here 200,000 or 500,000) to estimate the expected SFS, we also compared the likelihoods of the models estimated on the basis of 10 million coalescent simulations done under the ML parameters. We repeated this procedure 100 times per model to check if the distributions of these likelihoods were overlapping and thus not distinguishable. We used the following command line to get these 100 likelihoods:

*fsc -i xxx_maxL.par -R100 -n10000000 -d -u -C5 --logprecision 18 -q*

Finally, we also computed the Akaike criterion (AIC) (*82*) and the model relative likelihoods assuming site independence, even though our "neutral" sites may be linked on some chromosomes.

Confidence intervals

Confidence intervals around ML parameter point estimates were obtained via a parametric bootstrap approach, for which we first generated 100 SFS covering $T_{neutral\_X}$ nucleotides using estimated ML parameters, with the command line:

*fsc -i xxx.par -n100 -j -d -s0 -x –I -q -u*

Then, for each of these bootstrapped SFS, we re-estimated the parameters of the model using 20 independent runs starting at the ML parameters value (option *--initvalues* in *fastsimcoal2*). We used 60 ECM cycles for each run and performed 500,000 simulations for estimating the expected SFS under a given set of parameter values and to estimate the model likelihood. The *fastsimcoal2* command line used for the bootstrap was of the type:

*fsc -t xxx.tpl -n500000 -d -e xxx.est --initvalues xxx.pv -M -l60 -L60 -q -C5 --multiSFS --logprecision 18*

The limits of 95% confidence intervals were finally estimated by computing the 2.5% and 97.5% quantiles of the distribution of the 100 newly estimated ML parameter values.

### Supplementary Text

#### ***Archeological context of the samples***

In this study, we present new whole-genome sequences for 15 ancient human individuals (see Table S7 for an overview). Our sample set consists of 2 Mesolithic individuals from Vlasac (Serbia), 2 individuals from Transitional and Neolithic layers at Lepenski Vir (Serbia) and 11 early Neolithic individuals. The Neolithic samples originate from Aktopraklık and Barcın in Turkey (1 individual each), from Nea Nikomedeia in Greece (2 individuals), from Vinča-Belo Brdo and Grad-Starčevo in Serbia (1 each), from Kleinhadersdorf and Asparn-Schletz in Austria (1 each) and Essenbach-Ammerbreite, Dillingen-Steinheim and Herxheim in Germany (1 each).

#### Archaeological background

Anatolia and the Aegean appear to have been sparsely populated at the height of the Last Glacial Maximum, ca. 26-20 kya (*83–89*). The few cave sites that are securely dated to this period, including Asprochaliko and Kastritsa in Epirus, Theopetra in Thessaly and Franchthi in Argolis, are thought to have been mainly used seasonally as hunting stations and do not show much stratigraphic time-depth (*87*). Aurignacian industries are succeeded by Gravettian industries ca. 26 kya in mainland Greece, which however lack the typical Gravette points observed elsewhere in the Balkans (*86, 87*).

The onset of the northern Hemisphere deglaciation, ca. 20-19 kya (*90*), is associated with major changes in the Aegean region. Eustatic rise in sea-level, in the order of 120 m by the mid-Holocene (*91, 92*), is thought to have caused the breakup of the Cycladic mega-island (*93, 94*) and, from ca. 9.5 kya, the flooding of the Black Sea (*95*). Epigravettian industries, characterized by small backed blades with abrupt retouch, perhaps used as inserts for composite projectiles, are found on both sides of the Aegean Basin and in the Antalya Bay (*28, 96*). The emergence of new seafaring networks in the Aegean Basin is indicated by the long-distance procurement of obsidian, a volcanic glass found on the island of Melos, from the Final Palaeolithic onwards (*97*).

**Table S7 - Archeological information on the new palaeogenomes included in this study.**
The $^{14}$C dates were calibrated in OxCal 4.4.2 (*98*) using the IntCal20 calibration curve (*99*). Dates from Danube Gorges individuals with associated $\delta^{15}$N>8.3‰ were corrected for freshwater reservoir effect (FRE) following the method described by (*100*), later amended by (*101*). Both new (marked with an asterisk) and previously published dates are reported here.

| Sample ID | Site | Phase | Archaeological/ Burial ID | 14C Lab Number, uncal. age BP | cal. BP 2σ |
|---|---|---|---|---|---|
| VLASA7 | Vlasac | Late Mesolithic | Burial 31 | AA-57777: 8196 ± 69<br>((*102*): Tab.1)<br>FRE corrected: 7707 ± 93 | 8764-8340 |
| VLASA32 | Vlasac | Late Mesolithic | Burial 16 | *MAMS-46044: 9064 ± 27<br>FRE corrected: 8596 ± 66 | 9741-9468 |
| AKT16 | Aktopraklık | Anatolian Late Neolithic (Aegean Early Neolithic) | 89 D 14.1 | MAMS-25475: 7792 ± 28<br>((*105*): Tab.7) | 8635-8460 |
| Bar25 | Barcın | Anatolian Late Neolithic (Aegean Early Neolithic) | M10-455<br>(BH 43347 (lot. 1856)) | *MAMS-46043: 7506 ± 27 | 8384-8205 |
| Nea2 | Nea Nikomedeia | Early Neolithic | #7 | MAMS-24004: 7290 ± 31<br>((*106*): Tab. 2.4) | 8173-8023 |
| Nea3 | Nea Nikomedeia | Early Neolithic | T XII | *MAMS-46042: 7388 ± 27 | 8327-8040 |
| LEPE48 | Lepenski Vir | Transformational/ Early Neolithic | Burial 122 | OxA-16005: 7190 ± 45 (Borić 2011: appendix 1),<br>FRE corrected: 7115 ± 46 | 8017-7844 |
| | | | | OxA-16006: 7190 ± 40 (Borić 2011: appendix 1),<br>FRE corrected: 7127 ± 41 | 8020-7860 |
| | | | | OxA-16005 + OxA-16006: 7122 ± 31<br>X2-Test: df=1 T=0.0(5% 3.8) | 8012-7867 |
| LEPE52 | Lepenski Vir | Early-Middle Neolithic | Burial 73 | BA-10652: 7265 ± 30 (Borić and Price 2013: ESM),<br>FRE corrected: 6983 ± 47 | 7931-7693 |
| STAR1 | Grad-Starčevo | Early Neolithic (Starčevo) | Grave 1 | BRAMS-2407: 6671 ± 27<br>(*107*) | 7589-7476 |
| VC3-2 | Vinča-Belo Brdo | Early Neolithic (Starčevo) | Grave V (group burial) | OxA-28634/UBA-22463: 6581 ± 34<br>((*109*): Tab. 2) | 7565-7426 |
| Asp6 | Asparn-Schletz | Early Neolithic (LBK) | Ind 44 (646 Part 152, Schnitt 10 LM70) | *MAMS-46038: 6657 ± 26 | 7580-7473 |
| | | | | *MAMS-48728: 6627 ± 35 | 7573-7430 |
| | | | | MAMS-46038 + MAMS-48728: 6646 ± 21<br>X2-Test: df=1 T=0.5(5% 3.8) | 7575-7474 |
| Klein7 | Kleinhadersdorf | Early Neolithic (LBK) | Grave 56 (25,937) | *MAMS-46040: 6208 ± 27 | 7244-7000 |
| | | | | VERA-2167: 6090 ± 50<br>((*108*): tab.36) | 7158-6796 |
| | | | | MAMS-46040 + VERA-2167:<br>X2-Test: df=1 T=4.277(5% 3.8) - combination fails at 5% | – |
| Ess7 | Essenbach-Ammerbreite | Early Neolithic (LBK) | Grave 2 | – | – |
| Dil16 | Dillingen-Steinheim | Early Neolithic (LBK) | Grave 24, Befund 24 (1997) | *MAMS-46039: 6200 ± 25 | 7235-6998 |
| Herx | Herxheim | Early Neolithic (LBK) | 281-19-6 | *MAMS-46041: 6189 ± 25 | 7164-6993 |

First settled communities appear on the Central Anatolian Plateau ca. 15.5-15 kya, shortly before or at the time of the Bølling-Allerød warming phase (*110*). The Epipalaeolithic site of Pınarbaşı in the Konya Plain, has produced geometric microliths, including lunates, showing influence from contemporary Levantine pre-Natufian and Natufian communities (*110*). Similar tools appear later on in the Antalya region at Öküzini (*111*). Large numbers of marine shells at the site of Pınarbaşı 1,000 m above sea level suggest strong links with the Mediterranean area during the Late Pleistocene, despite the natural boundary formed by the Taurus mountains (*112*). It is not clear how the cold spell of the Younger Dryas affected interactions between the Anatolian Plateau and the Aegean Basin. There is limited evidence for occupation during the interval 12.9-11.7 kya at Pınarbaşı (*110*, *112*).

Recent surveys in Western Anatolia, in the Karaburun and Bozborun peninsulas (*113*, *114*), fill an important gap between Epipalaeolithic traditions on the Mediterranean sea shore, e.g. Öküzini Cave in the Antalya Bay (*111*), and contemporary traditions in the North Aegean, e.g. Ouriakos, on the island of Lemnos (*115*). Characteristic lithic scatters found in the course of these surveys indicate the presence of foragers in Western Anatolia at the end of the Pleistocene, who shared similar tool sets (e.g. geometric microliths, backed blades and bladelets) and lithic technology, and were presumably integrated in wider Levantine-East Mediterranean-Aegean maritime interactions during the Younger Dryas (*115*). The picture changes at the beginning of the Holocene (coinciding with the start of the Aegean Mesolithic) when Western Anatolia shows stronger connections with the Aegean islands and the Greek mainland. There was limited interaction at that time with aceramic Neolithic sites of Southwest Asia (*113*).

In the Fertile Crescent, the Neolithic or agricultural 'revolution' is thought to have reached a tipping point by about 11.7 kya, at a time of dramatic post-Pleistocene climate change and expanding carrying capacities (*116–118*). Sites in what is today Southeast Turkey, Northern Syria and Western Iran, are among the first to show unambiguous signs of ungulate domestication (*1*, *119*). From about 10.6 kya, a range of founder crops and animals from the Fertile Crescent are introduced piecemeal among sedentarizing communities of Central Anatolia and Cyprus (*110*, *120*, *121*). The small 'aceramic' site of Boncuklu, is thought to be one of the oldest communities on the Anatolian plateau to practice small-scale plant cultivation, starting ca. 10.3 kya (*122*).

The Western half of the Anatolian Peninsula does not transition to farming before about 8.7/8.6 kya ((*123*, *124*) and references therein). **Barcın** and **Aktopraklık** are among the first Neolithic communities in NW Anatolia (*20*, *125*, *126*). The new subsistence economy, characterized by a near-complete 'package' of domestic crops and animals, is abruptly introduced by farmers, who live on settlement mounds, build elaborate rectilinear houses, and use pressure technology to create sets of tools for domestic activities, such as plant harvesting (*121*, *127–129*). Broadly comparable early Neolithic communities appear all over mainland Greece and Crete approximately within a century (*130–135*). **Nea Nikomedeia** belongs to an advanced phase of that expansion.

From about 8.2 kya, at a time of rapid climate change, farming spreads northwards to the Central Balkans, along the Struma, Mesta and Vardar-Morava-Danube corridors (*136*, *137*). The distribution of $^{14}$C-dated Neolithic sites shows a gradient away from the main river axes, with Alpine regions left mostly off-course during the initial phase of Danube expansion(*138*). Neolithic groups settled in enclaves along the main river system all the way to the Carpathian Basin, for instance at **Vinča-Belo Brdo** and **Grad-**

**Starčevo** near today's Belgrade. The narrow stretch of the Danube river known as the Danube Gorges or Iron Gates bears witness to important cultural and genetic interactions between indigenous sedentary fishers and incoming farmers at **Vlasac** and **Lepenski Vir** (*103*, *139–142*).

The emergence of the Linearbandkeramik (LBK) in Central Europe ca. 7.5 kya involves a major redefinition of the Neolithic pattern of existence, with settlement mounds now abandoned in favour of flat extended sites. A steep decline in crop diversity is observed north of the Alps, with a number of southerly-oriented species like lentils and chickpeas dropped from the Neolithic 'package' (*143*). Typical long houses appear in this period. The dead are increasingly buried in formal cemeteries outside settlement areas, such as **Kleinhadersdorf** in Lower Austria (*144*). Most LBK burials date to an advanced phase of the early Neolithic after ca. 7.2 kya.

Increased competition between different LBK communities of Central Europe is thought to have resulted in the first large-scale 'massacres' (*118*). **Asparn-Schletz** and **Herxheim** show signs of interpersonal violence on a scale that has never been observed before in the Neolithic (*145–148*). At Asparn-Schletz, one LBK community is thought to have attacked another, resulting in tens of dead lying unburied in the latest ditch system (*149*). In Herxheim, several LBK groups are thought to have come together to participate in a large-scale ritual event, involving the systematic killing and dismembering of hundreds of people from outside the community, whose bones were then meticulously smashed into small fragments whereas the skulls were manipulated leaving only the calottes intact. Besides the human victims, which are thought to represent human sacrifices, precious pottery and stone tools were also destroyed, underlining the ritualistic character of the event. Body fragments, calottes and artefact remains were finally deposited in large finds concentrations in two ditches surrounding the former settlement (*150*, *151*).

**Aktopraklık**

Excavated since 2004 by a team from the Prehistory Department of the University of Istanbul, Aktopraklık is one of the oldest Neolithic sites in Northwest Anatolia (*125*). The site is located on one of the terraces overlooking the eastern edge of Lake Uluabat, ca. 25 km west of the city of Bursa. Unusually for the Anatolian Neolithic, Aktopraklık is a flat extended settlement and cemetery, consisting of three distinct areas (Aktopraklık A, B and C). Previously-published AMS dates on human bones indicate that occupation spanned from the early 9$^{th}$ to mid-8$^{th}$ millennium BP (*105*, *125*). The oldest levels (Area C) are characterized by circular semi-subterranean structures, ca. 3-6 m in diameter, which were perhaps used as houses or ‘huts’ and fit the coastal Fikirtepe tradition defined by Mehmet Özdoğan in the NW Anatolia region ((*20*): S422-S423). Previous studies have indicated that the first inhabitants at Aktopraklık, including the individual sampled here, consumed terrestrial $C_3$ food and practiced animal husbandry (*128*, *152*, *153*).

In the Neolithic phase, the dead were buried in close proximity to houses or immediately beneath them, sometimes with monochrome pottery, bone tools and stone beads (*154*). Individual 89D 14.1. (AKT16) stems from a double burial and was identified genetically as female (Fig. S9). This middle-aged to old adult was buried in a contracted position on her left side with the head oriented to the east – facing south ((*155*): 25). The head of individual 14.1 was placed on the feet of adult female 17.1, who shared the same grave. There were no grave goods as such except faunal remains in the pit fill and on the skeletons. Direct dating of this skeleton (MAMS-25475: 7792 ± 28 BP) indicates that she died ca. 8635-8460 cal. BP at 2σ - in other words at the very beginning of farming expansion in the region ((*105*): tab 7; tab 21).

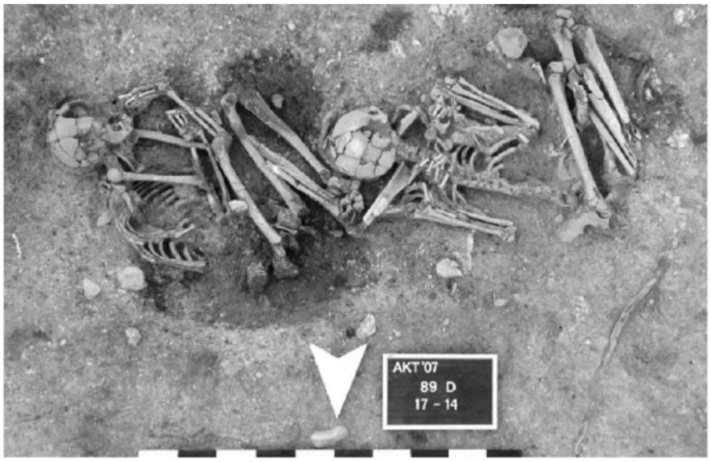


**Figure S9 - Aktopraklık**: double burial 89D 17.1 and 14.1 ((*155*): fig.9)

**Barcın**

Barcın Höyük is a multi-layered mound in Northwest Anatolia, excavated from 2005 to 2015 by an international team under the auspices of the Netherlands Institute in Turkey (*126*). The site, which has already featured in several ancient DNA studies (*5*, *7*, *156*), is located in the Yenişehir Basin, between the Iznik Lake and the Uludağ Mountains, providing convenient access to the heart of the Anatolian Peninsula. A series of over 30 radiocarbon dates place the beginning of the Barcın occupation (levels VIe and VId1) 8,550 cal. BP (*123*). Despite its early date, Barcın is characterized from the outset by a fully agrarian economy, based on cultivated cereals and pulses, while domestic cattle and sheep dominate the faunal assemblage (*121*).

The settlement structure in the early phases consists of a row of rectilinear mud and timber houses facing a central courtyard. The dead are generally buried in the courtyard, except neonates and infants who are also buried within or directly outside houses, next to walls. The individual Bar25 sampled here (M10-455) was found with several other infant burials in a narrow annex of structure 19, a structure which had an unusual red-coloured floor – a feature often associated with "special purpose" buildings in Southeast and Central Anatolian Neolithic sites (*157*). This and neighbouring buildings of level VId1 were all destroyed by a fire, which happened no later than 8350-8325 cal. BP according to the latest radiocarbon determinations. Culturally speaking, Barcın demonstrates early and ongoing connections with Central Anatolia, visible in such elements as the production and uses of ceramics, the incorporation of cattle and goat horns in the architecture and the predominance of Central Anatolian sources in the obsidian assemblage (*126*).

Burial M10-455 is a single primary inhumation in a small pit. The young infant, genetically identified as male, was placed in an east-west orientation with cranium to the east. There were no grave goods associated with the burial. The skeleton has been directly dated to 8384-8205 cal. BP at 2σ (MAMS 46043: 7506 ± 27 BP).

**Nea Nikomedeia**

The early Neolithic mound of Nea Nikomedeia in Northern Greece was excavated from 1961 to 1964 by Robert J. Rodden under the auspices of the British School at Athens. The site, which is located ca. 10.5 km north-east of the city of Veroia in the plain of Macedonia, has produced three building phases consisting of broadly overlapping rectilinear timber structures with foundation trenches (*158–161*). The excavation project marked an important turning point for research on Macedonian prehistory, due to the involvement of multiple teams of international scientists and the recognition of Nea Nikomedeia as one of "the oldest Neolithic sites found in Europe" ((*160*): 83). An unusually large building at the site, described by the excavators as a "shrine", was destroyed in a violent fire with all its contents, including elaborate female figurines ((*162*): 564). Based on a series of radiocarbon dates obtained in the 1990s on archival samples of domestic cereals and bones, the early Neolithic site of Nea Nikomedeia has been dated to the second half of the $9^{th}$ millennium BP, perhaps starting ca. 8400 cal. BP.

The funerary record of the site has never been published in detail, although preliminary reports suggest that at least 35 individuals came from regular burials inside the early Neolithic settlement, including 13 adults, 13 children and nine infants; in addition, 31 adults and 21 children were represented by single bones or partial skeletons ((*163*): 103; (*164*): tab. 4.5; tab. 4.7). Most burials were single primary inhumations placed in earth pits, in contracted position under house floors or in-between houses ((*159*): 605).

The two individuals sampled here, both genetically females, have been radiocarbon dated to 8327-8040 cal. BP (Nea3, MAMS-46042: 7388 ± 27 BP) and 8173-8023 cal. BP (Nea2, MAMS-24004: 7290 ± 31 BP). Based on the macroscopic examination of the skeletal remains, which are in extremely poor condition, both individuals appear to be early infants, respectively ~6-9 (T XII, Nea3) and ~18 months old (#7, Nea2). The latter shows enlarged nutrient foramina on the diaphyses and flaring of the epiphyseal ends of the long bones. These skeletal changes have been linked to hereditary anemias such as thalassemia and sickle cell anemia (*165–167*). Differential diagnosis for flaring and frayed metaphyseal cortex also includes rickets/osteomalacia and genetic syndromes (*166*).

**Vlasac**

Set like its neighbour Lepenski Vir in the spectacular landscape of the Danube Gorges, Vlasac has been first salvage-excavated by D. Srejović and Z. Letica in 1970-1971; a team led out by D. Borić returned to the site in 2006-2009 to conduct new excavations on the unsubmerged section of the settlement. The individuals analysed in this study both originate from the old Srejović excavation area. Remarkably, some of the architectural practices observed at Lepenski Vir, such as trapezoidal floor plans, rectangular hearths and red limestone floors, are already present (if only in basic form) in Late Mesolithic contexts at Vlasac.

Burial 31 (VLASA7) is a typical Mesolithic burial in extended supine position (Fig. S10). The burial, which belongs to Phase I, truncated the eastern side of Dwelling 2, which is dated to the first half of the $9^{th}$ millennium BP according to Borić *et al.* ((*102*): 273-274). A single radiocarbon date (AA-57777: 8196 ± 69 corrected 7707 ± 93 BP) from the skull provided a range of 8764-8340 cal. BP at 2σ after correction for freshwater reservoir effect ((*103*): appendix 1). This adult individual, identified as genetically male ((*105*): tab S3), has produced high stable isotope values ($\delta^{15}N$: 16.1‰ and $\delta^{13}C$: -20.7‰) consistent with substantial intake of aquatic proteins ((*103*): appendix 1). Previously published $^{87}Sr/^{86}Sr$ ratios indicate a local range for this individual ((*104*): SI tab. 1).

Mesolithic Burial 16 (VLASA32) consists of the disarticulated remains of an old adult male (Fig. S10), radiocarbon dated to 9741-9468 cal. BP at 2σ after correction for freshwater reservoir effect (MAMS-46044: 9064 ± 27 corrected 8596 ± 66 BP). The previously published $^{87}Sr/^{86}Sr$ ratio for Burial 16 is consistent with a local range ((*104*): SI tab. 1).

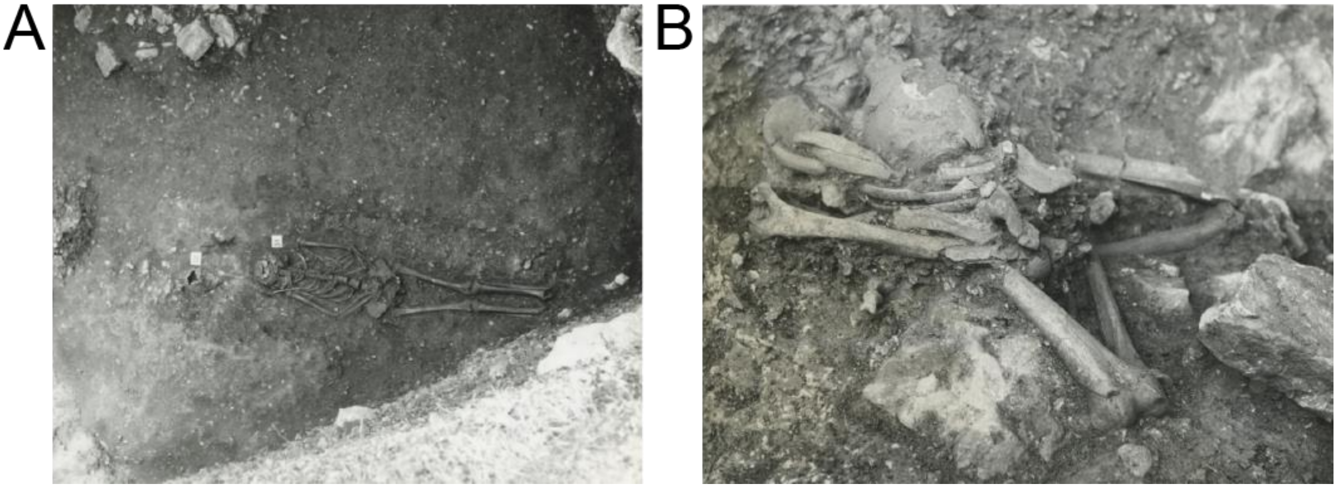

**Figure S10 - Vlasac.** (**A**) Burial 31 in extended supine position. (**B**) Remains of Burial 16 (image credit: Documentation of the Archaeological collection, Faculty of Philosophy, University of Belgrade).

**Lepenski Vir**

Lepenski Vir is one of the most emblematic Mesolithic and early Neolithic sites in Europe. Completely excavated from 1965 to 1970 by Dragoslav Srejović, in advance of the construction of the hydroelectric Đerdap I dam in the Danube Iron Gates, Lepenski Vir has since been extensively re-studied by teams from the Universities of Belgrade and Cambridge. The osteological and mortuary evidence have been the focus of a number of recent high-impact publications, making the site an ideal case-study to address the transition from Mesolithic to Neolithic (*105*, *139*, *168*). Occupied from the 12$^{th}$ millennium BP by foragers and fishers, Lepenski Vir bears witness to important changes after 8150 cal. BP, when pottery appears in the Central Balkans. The more iconic trapezoidal houses with limestone floors, sculpted boulders and subfloor burials belong to this period. Despite the arrival of early Starčevo farming communities in the region, there are no domesticates beside dogs at Lepenski Vir before Phase III, dated to the early 8$^{th}$ millennium BP ((*169*): 52).

The two individuals reported in this study, which are very close chronologically, belong to the Transformational/early Neolithic phase I-II (Burial 122) and the early-Middle Neolithic Phase III (Burial 73). Burial 122 (LEPE48) is a stray sub-adult skull without a mandible (15-18 year old, genetically male) discovered in the packing layer between the superimposed floors of trapezoidal houses 47 and 47’, at the centre of the settlement (Fig. S11). Two radiocarbon dates (OxA-16005: 7190 ± 45 corrected 7115 ± 46 BP and OxA-16006: 7190 ± 40 corrected 7127 ± 41 BP) directly date this skull to 8012-7867 cal. BP at 2σ after correction for reservoir effect ((*139*): 211). Strontium isotope analysis has previously indicated that skull burial 122, which bears intriguing cut marks, falls just outside the local range ((*104*): fig. 3).

Burial 73 (LEPE52) is a single primary inhumation of a genetically adult male ((*105*): tab 22), ca. 30-40 years old, crouched on his right side in the northern zone of Lepenski Vir, outside the habitation space (Fig. S12). A single radiocarbon date (BA-10652: 7265 ± 30 corrected 6983 ± 47 BP) places Burial 73 to 7931-7693 cal. BP at 2σ after correction for freshwater reservoir effect (see (*104*): ESM; (*139*): tab 1.1; 293). Isotopically, Burial 73 is in the local range ((*104*): fig.3). Starčevo pottery and a green stone pendant were recovered in the fill of this burial.

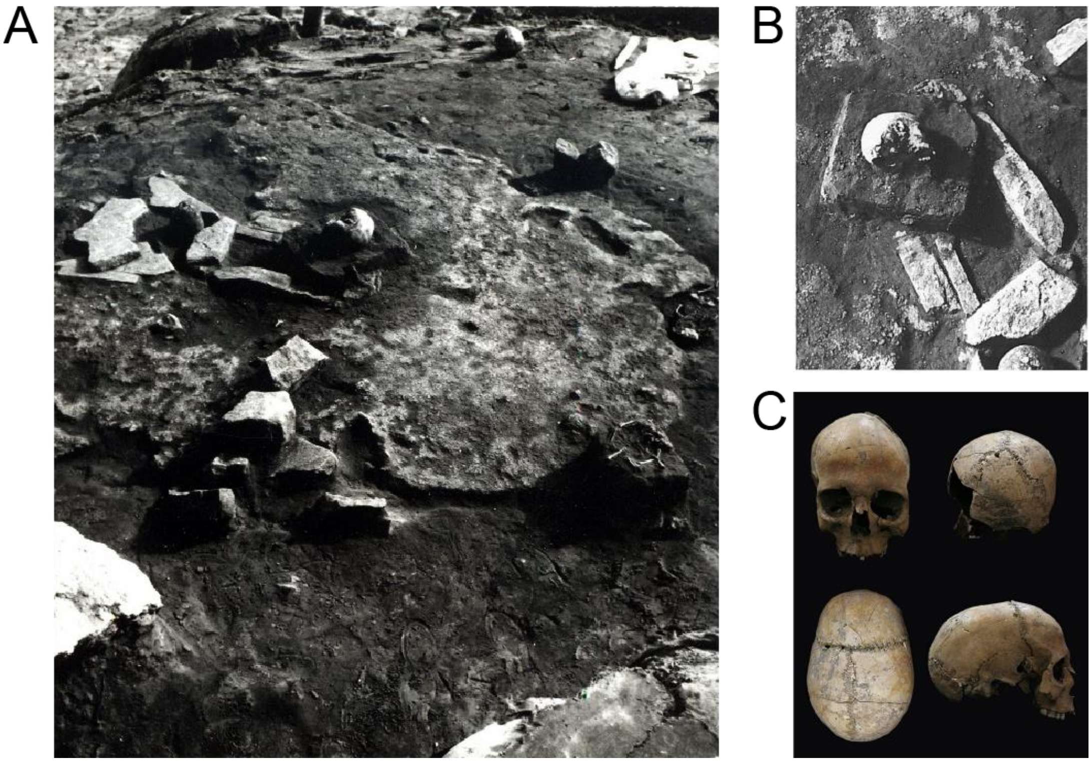

**Figure S11 - Lepenski Vir**. (**A**) Burial 122; the skull with house 47'; (**B**) detail; (**C**) frontal, occipital, vertical and lateral projection (image credit: reproduced after (*140*): fig. 68-69).

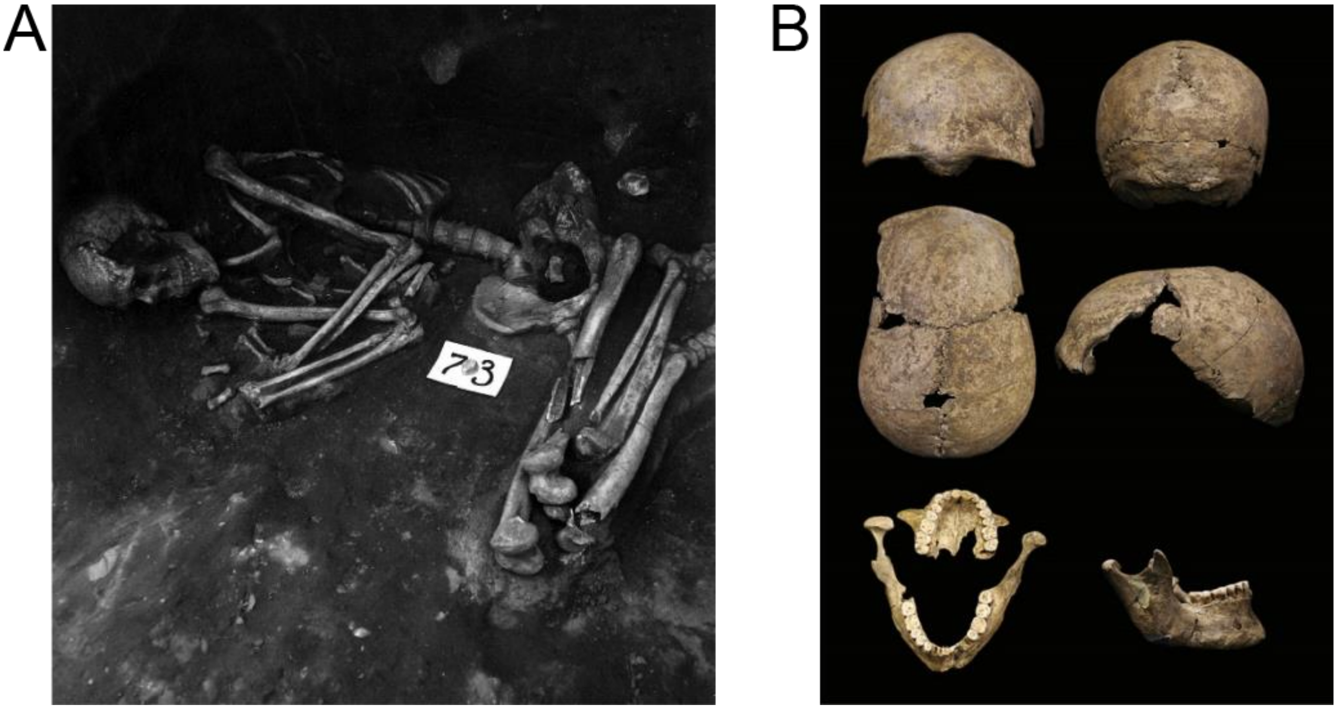


**Figure S12 - Lepenski Vir.** (**A**) Burial 73; (**B**) skull of Burial 73, frontal, occipital, vertical and lateral projection and maxilla and mandible (image credit: reproduced after (*140*): fig. 52-53).

**Grad-Starčevo**

Situated just across Vinča on the other side of the Danube, along an old bank of the river that was liable to flooding until the turn of the $20^{th}$ century, the type-site for the Starčevo culture in the Central Balkans has been repeatedly excavated since 1928 by teams from the National Museum in Belgrade ((*170*): 59). The first systematic excavations were carried out in 1932 by Vladimir Fewkes with funding from the Peabody and Fogg Museums at Harvard and from the American School of Prehistoric Research ((*170*): 60). New excavations took place in 1969-1970 under the direction of Draga Garašanin and Robert Ehrich. In 2003-2004, the Institute for the Heritage Protection in Pančevo conducted a series of small trench excavations totaling about 70 $m^2$ ((*171*): 54).

The site, which has been traditionally ascribed to the later Starčevo period, is radiocarbon-dated to ca. 7900-7350 cal. BP (*172*). Grad-Starčevo has documented a range of domestic activities typically associated with Neolithic societies, including food-production, some degree of sedentism with 'pit-houses' (though no postholes can convincingly be associated with them) and a rich ceramic assemblage including monochrome and painted vases ((*170*): 64-65). Domestic animals include cattle, sheep/goat and pigs ((*173*): tab 5).

The individual sampled for this project, Skeleton 1, Grave 1 (STAR1), which is now radiocarbon dated to 7589-7476 cal. BP at 2σ (BRAMS-2407: 6671 ± 27 BP), was discovered during the 2004 excavation season at the bottom of the pit dwelling in trench 5/2004 ((*171*): appendix 1). Identified as an older female, ca. 55-60 years old, Skeleton 1 was buried in a contracted position on the right side, without any grave goods (Fig. S13). New isotopic values for this individual indicate that she had a mostly terrestrial diet (*107*). A 5-7 year old child was found nearby, at a distance of 0.6 m on the same level ((*171*): appendix 1).

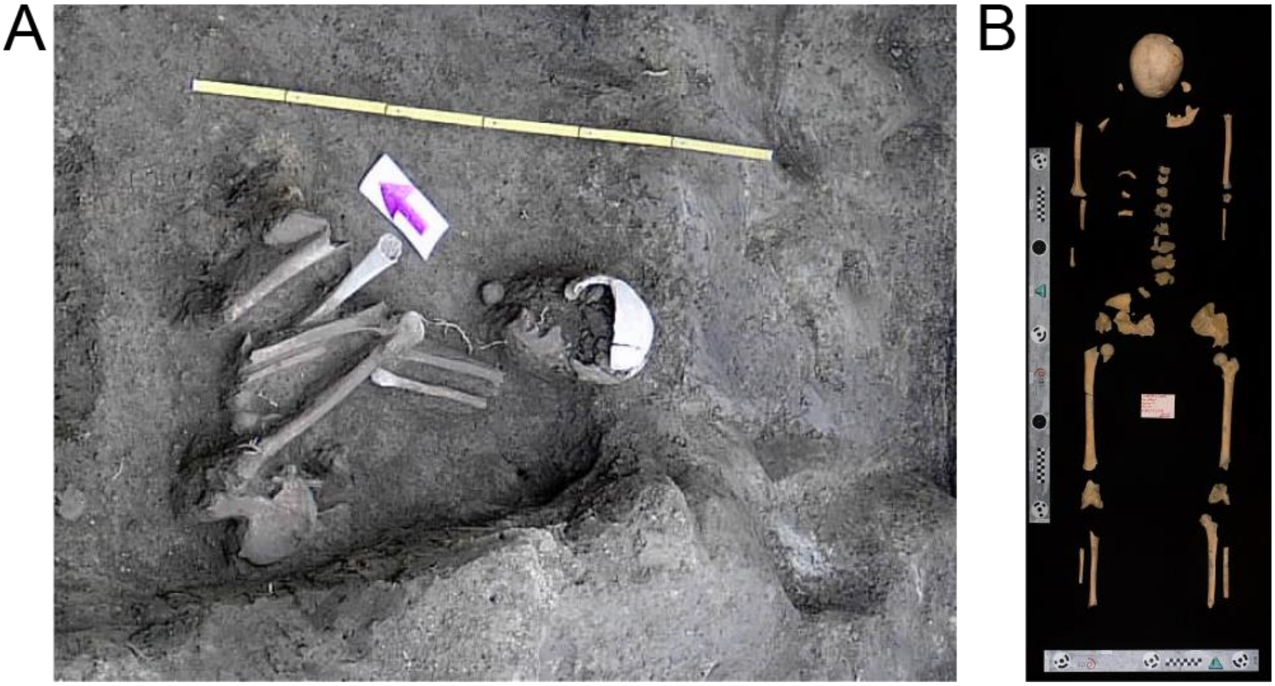


**Figure S13 - Grad-Starčevo.** (**A**) Grave 1; (**B**) detail (image credit: Documentation of the Monument Protection Institute - Pančevo/Jugoslav Pendić).

**Vinča-Belo Brdo (Starčevo phase)**

Located on the right bank of the Danube, ca. 14 km east of Belgrade, the site of Belo Brdo in Vinča has been excavated from 1908 onward by Miloje Vasić, rapidly becoming the type-site for the eponymous Vinča culture in the Central Balkans. The site is unusual in being the only known tell or settlement mound in a radius of 100 km, at 8 m above the Danube river valley ((*174*): 205). New excavations began in 1978 under the auspices of the Serbian Academy of Sciences and Arts (*175*). The site was first occupied in an advanced phase of the Starčevo period before the development of the Vinča culture.

The individual sampled for this project, individual V (VC3-2), stems from a collective tomb (pit Z) in the lowermost level of the mound (Fig. S14). The partial commingled remains of at least 12 individuals, including two females and nine indeterminate adults and sub-adults, were associated with this feature

((*171*): 58-59), which has been variously interpreted as an “ossuary”, a “pit-dwelling” and a “tomb with entrance hall – dromos” ((*176*): 230-232). Individual V forms part of the collection studied by I. Schwidetzki in 1937, which was damaged during the bombing of Belgrade in WWII ((*171*): Appendix 1). In her PhD thesis, Jelena Jovanović indicates that individual V consists exclusively of skull fragments; age could be inferred to between 25-35 years based on tooth abrasion; isotopic values of $\delta^{13}C$, $\delta^{15}N$ and $\delta^{34}S$ provided by Nehlich *et al.(177)* are consistent with a terrestrial diet with a small intake of aquatic proteins ((*171*): Appendix 1). Individual V has a genetic signature consistent with male ((*105*): tab S3). The Starčevo burials have been radiocarbon dated in the 2000s to between 7669-7324 cal. BP at 2σ after correction for reservoir effect (*109*, *178*). Given the $\delta^{15}N$ values of between +10.3-13.6‰ (Δ = 3.3‰; n = 6) for some of these individuals, possibly indicating some intake of freshwater fish ((*176*): 230-231; (*177*): tab.3; (*141*): ESM3), a small reservoir correction factor equivalent to that used in the Danube Gorges was applied by Tasić *et al. (109)*.

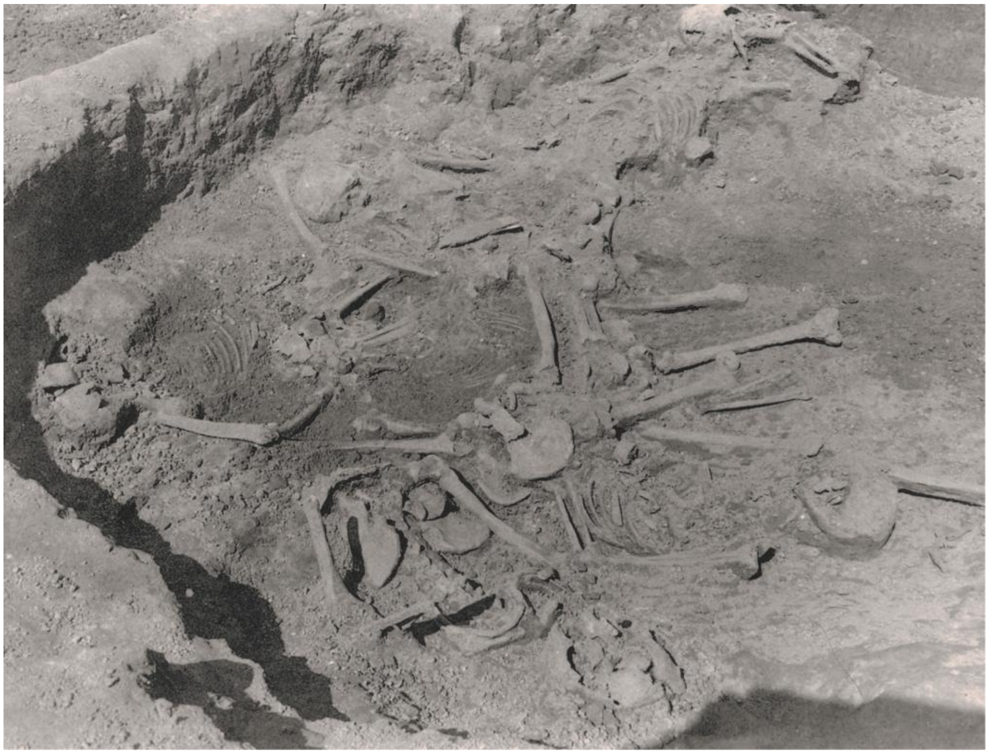

**Figure S14 - Vinča:** collective tomb of the Starčevo period (image credit: Documentation of the Archaeological collection, Faculty of Philosophy, University of Belgrade)

**Asparn-Schletz**

Excavated from 1983 to 2005 by Helmut Windl, Niederösterreichisches Landesmuseum, the site of Asparn-Schletz, in the Lower Austrian Weinviertel, is one of the most enigmatic Linearbandkeramik culture (LBK) sites. Schletz is a large settlement enclosed by two ditch systems, a trapezoidal one, ca.

400 m in diameter, and an oval one, ca. 330 m in diameter ((*149*): 191). The latter was renewed several times, broken by several entranceways or earthen bridges and probably reinforced by a palisade. There is evidence for habitation, in the form of long houses, and food production at the site ((*179*): 56). In its latest phase, the oval ditch system provided a final resting place for approximately 130 individuals, who were given no formal inhumation and often lay in unnatural contortions.

The skeletons, which have been studied by anthropologist Maria Teschler-Nicola and colleagues, bear multiple traumas of perimortem origin, mostly caused by blunt force to the skull and arrow piercing (*148*). Some perimortem fractures on postcranial remains were also identified. It is likely that the corpses remained unburied for a period of time as limb extremities are often missing, and bones bear gnaw marks. Presumably the 'massacre' was a single event which happened some time towards the end of the LBK, ca. 7150-6900 BP, based on a sequence of 15 internally consistent $^{14}$C dates on human bones ((*180*): table 1). Young females are underrepresented in the mortality profile (only five females to 17 males in the 20-40 years age bracket). The abandonment of the settlement appears to have coincided with the 'massacre' itself; the final ditch system was partially infilled with settlement debris ((*149*): 192). Up to now ca. 20 regular graves have been recorded in the ditch and the settlement beside the individuals of the massacre. Most individuals buried in graves were subadults. Regular graves came from an earlier LBK level.

The individual sampled for this project, ind. 44 (Asp6) from the 1987 excavation season, comes from the juncture of the two ditches, which precisely overlap at this point, and possibly belongs to the 'massacre' context. The individual has been identified, both anthropologically and genetically, as male. Age at death has been recorded as between 35-45 years. The cranial remains exhibit not only an old healed trauma, but also perimortal fractures. In the course of the new Forschung-Technologie-Innovation (FTI) Strategy project, supported by the Lower Austrian government, the excavation records and other documents are being systematically re-investigated. For this burial, earlier excavation records indicate that the individual was "buried in the ditch, E-half".

The new $^{14}$C date reported here, which stems directly from the skeleton sampled (MAMS-46038: 6657 ± 26 BP, 7580-7473 cal. BP at 2σ) is significantly older than anticipated, based on its stratigraphic location on top of other slain individuals (F. Pieler, comm.). The $^{14}$C date was confirmed by repeating the graphitization and measurement, using collagen prepared for MAMS-46038 (MAMS-48728: 6627 ± 35 BP, 7573-7430 cal. BP at 2σ). The ongoing FTI project, led by Franz Pieler and Maria Teschler-Nicola, will help to clarify the dating of this individual.

**Kleinhadersdorf**

Discovered in 1911, the Linearbandkeramik (LBK) cemetery of Kleinhadersdorf near Poysdorf, in the Lower Austrian Weinviertel, has been first excavated in 1931; more systematic excavations to rescue the site from erosion and agricultural activities took place in 1987-1991, under the directorship of Johannes Wolfgang Neugebauer from the Austrian Bundesdenkmalamt. The findings, including the results of anthropological analyses, have been recently published in a monograph ((*144*), and references therein). The cemetery was located some 150-200 m north of the nearest LBK settlement, on a slope at the edge of a very fertile loess basin. Based on ceramic typology and a sequence of 19 radiocarbon dates, the site has been dated to ca. 7250-6750 BP, spanning from the phase LBK I/II to phase LBK III; at least four burial phases could be identified ((*144*): 110; 151-152).

Of the more than 60 graves excavated at the site, 41 have provided detailed anthropological information (*181*). Most were single primary inhumations in a contracted position on the left side ((*144*): 57-59). There was some variation in the layout and orientation of the bodies, and at least 12 were buried on their back. Beside regular graves, there were 26 ‘empty’ ones, which contained no or very few human remains, bringing the total number of individuals recorded at the site to 62. Intriguingly, several features point to a local continuation of Mesolithic traditions, including microliths, arrowheads, and deposits of red ochre in some of the grave pits ((*182*): 115-118).

The individual sampled, Klein7, comes from grave 56 (Fig. S15) and was buried in a contracted position on the left side, with the head to the northeast, the mouth wide open ((*144*): plate 36). The arms were folded together, with the left hand in front of the face covered in red ochre powder. The individual was identified as a mature female, ca. 40-50 years old, and shows signs of degeneration on the spine ((*181*): 341). There were perimortem fractures on the right humerus and right fibula. Ceramic sherds from the grave fill show an early Želiezovce ornament and should be dated to shortly after phase IIb of the site ((*144*): 98). The new radiocarbon date (MAMS-46040: 6208 ± 27 BP) is slightly older than the one published before for grave 56 ((*108*): tab.36; VERA-2167: 6090 ± 50 BP) and is consistent with the stratigraphy of the site, indicating that Klein7 should be dated to 7244-7000 cal. BP at 2σ. Previously published $\delta^{13}C$ (-20.0‰), $\delta^{15}N$ (9.8‰) $^{87}Sr/^{86}Sr$ (0.709612) values for this individual suggest that she sourced her food nearby and probably had access to meat ((*183*): tab.38).

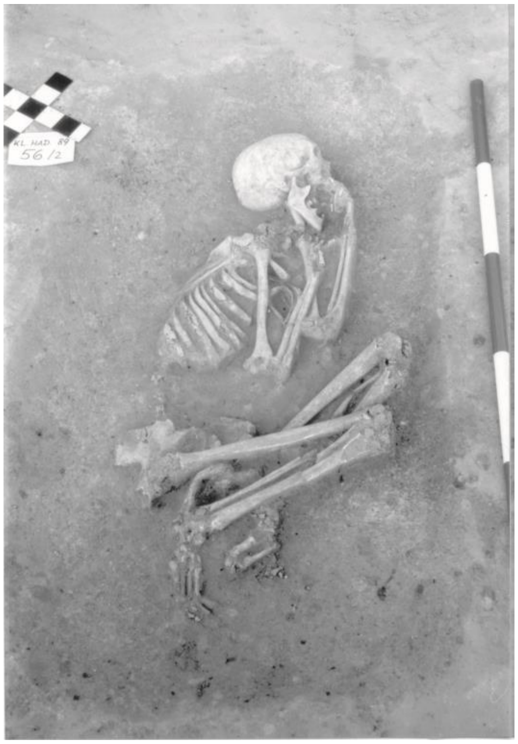

**Figure S15 - Kleinhadersdorf:** grave 56 (image credit: J.-W. Neugebauer).

**Essenbach-Ammerbreite**

The LBK cemetery and settlement of 'Ammerbreite' in Essenbach, Landshut county, Lower Bavaria, was discovered in 1981 during construction work on the north-western edge of the modern village. Subsequent excavations until 1986 by Henriette Brink-Kloke uncovered remains of LBK habitation and a concentration of 29 graves, ca. 40 m away, attributed to an advanced phase of the LBK. One additional burial (grave 7) was found inside a reused settlement pit in the direction of the cemetery ((*184*): 428). Only a section of the cemetery could be excavated. Graves were damaged by ploughing activities. Anthropological study of the skeletons determined that 10 were children or infants, and 15 adults or sub-adults, of which seven were female and six were male ((*184*): 433).

The individual sampled for this project (Ess7) was the child in grave 2, ca. 9-10 years old, buried in contracted position on the left side, with the head oriented to the northeast (Fig. S16). Grave 2 was one of the more richly furnished graves, including objects normally associated with adult males (graves 16 and 24), such as a stone axe and a complete vessel. The vessel contained the fragment of a perforated bone comb and the left, upper incisor tooth of the child ((*184*): 458). The dating of the skeleton has not been established through $^{14}$C dating. The objects deposited in the grave are typical of the LBK, with the comb suggesting a 'young' or 'late LBK' date, possibly in the range 7050-6900 BP (J. Pechtl, comm.).

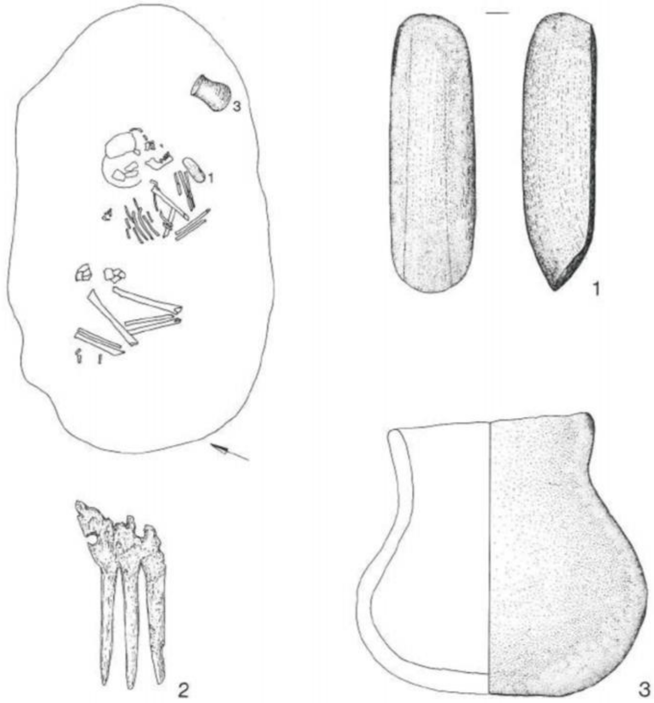


**Figure S16 - Essenbach-Ammerbreite:** content of grave 2 (image credit: Zeichnungen Bayerisches Landesamt für Denkmalpflege, Außenstelle Landshut, reproduced with permission of Germania(*184*)).

**Dillingen-Steinheim**

Located on a high terrace with thick loess soils, between the Danube and the Egau rivers, the LBK site of 'Steinheim' near Dillingen, Swabia, was excavated during a seven-month long season in 1987 by the Bavarian State Office for the Preservation of Monuments – Augsburg ((*185*): 60; (*186*)). The site consists of a short-lived LBK settlement possibly enclosed by a ditch. Two groups of burials were reported for this site – a small burial cluster, comprising up to 27 early Neolithic graves, some distance away from the settlement (*185*) and a group of regular graves within the ditch (J. Pechtl, comm.). There is no evidence of massacre at Steinheim; formal inhumations in LBK ditches are common in Southern Germany. The individual sampled (Dil16) comes from the ditch, belonging to an 11-13 year old, genetically male. The new radiocarbon date obtained from this skeleton (MAMS-46039: 6200 ± 25 BP, 7235-6998 cal. BP at 2σ) confirms that the ditch was infilled in an 'older LBK' phase, ca. 7150-7050 BP ((*187*): 16).

**Herxheim**

Located in the south of the Rhineland-Palatinate State in Western Germany, Herxheim is one of the most frequently discussed LBK sites in relation to Neolithic interpersonal violence alongside Schletz and Talheim. Initially a rescue excavation under Annemarie Häußer, in 1996-1999, the scientific responsibility of the project has since been entrusted to Andrea Zeeb-Lanz, from the General Directorate for Cultural Heritage Rhineland-Palatinate, who oversaw from 2005 to 2008 a new excavation and led a research project financed by the German Research Foundation (DFG) for eight years. Herxheim may have started off as a fairly conventional LBK site at around 5300 BC, enclosed during the so-called "ritual phase" by a series of elongated pits, which ended up forming two broadly parallel trapezoidal ditches or ditch segments, ca. 250 x 230 m in size; in places these run up to 4 m beneath the settlement level. Twenty-nine radiocarbon dates place the ritual phase of the settlement toward the end of the LBK, ca. 7160-7000 cal. BP at two standard deviations. Bone and pottery refits from the finds concentrations in the ditches suggest a much shorter time for the rituals and the infilling of the ditch segments ((*188*): 34).

The highly fragmented remains of over 500 individuals were found deposited in the two 'ditches' (Fig. S17), usually in heaps or scatters that also include pottery, faunal remains, stone and bone tools (*150*, *189*). The human bone assemblage is dominated by skulls, especially skull caps, which were in some cases deposited separately in little clusters. These were shaped in a regular manner, probably with the help of an adze, and often bear cut marks that indicate removal of the scalp. Individuals of all age groups and sexes are represented among the deceased. Patterns of systematic and targeted bone breakage and defleshing have led some to infer mass cannibalism at the site (*146*), though the same evidence can be used to suggest ritual killing or 'sacrifice' (*151*).

Much of the pottery deposited in the pits appears to be exotic, with stylistic elements indicating connections with communities as far east as the Elbe Valley in the North of today's Czech Republic (Bohemia). Hence a question arises as to whether the dead might have come from as far as the pottery itself. Strontium isotope analysis on tooth enamel of nearly eighty individuals, including two regular LBK 'hocker' graves from the inner pit ring, shows an intriguing pattern in which those individuals that have received formal inhumation fall within the local range of variation, while disarticulated or dismembered human remains belong to 'non-locals' (*190*). The majority of the Sr-values hint at an origin of the butchered individuals from low mountain ranges, totally untypical for LBK settlement patterns.

The *Pars petrosa* sampled for this project (Herx), which belongs to a genetically female individual, comes from a stray skull fragment recovered in the northern end of the outer trench ring, section 281-19, excavated in 1998. The 'ditch' is about 2.5 m deep and V-shaped in this section, yielding only isolated human bones, faunal remains, loam and pottery fragments from a depth of 40 cm down. The infilling is likely to have taken place in the late/final LBK, ca. 7000 cal. BP, based on interpretation of the $^{14}C$ dates with Bayesian methods (*191*) and ceramic finds including decorated sherds. The individual reported here is now $^{14}C$ dated to 7164-6993 cal. BP at 2σ (MAMS 46041: 6189 ± 25 BP).

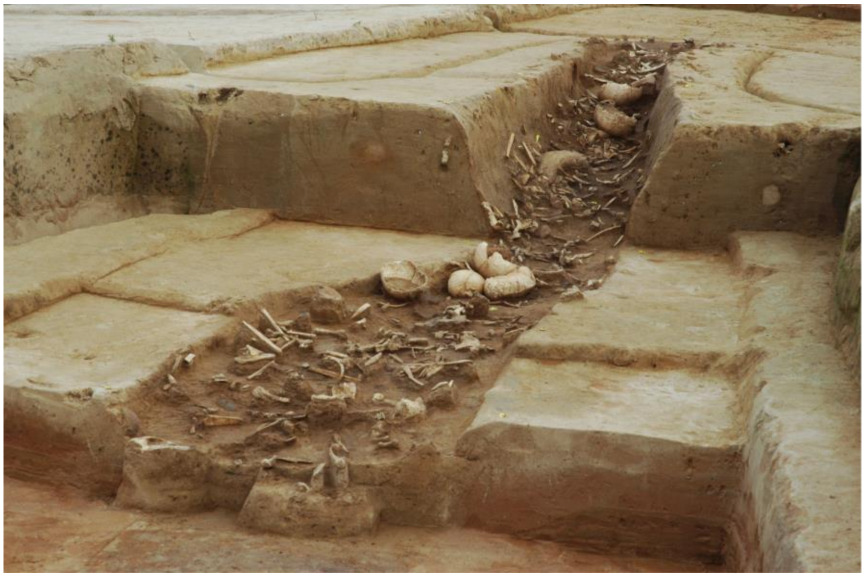

**Figure S17 - Herxheim:** one of the bigger find concentrations (image credit: GDKE - Landesarchäologie Speyer).

**Radiocarbon dating and stable isotope analysis**

We report new radiocarbon dates (see Table S7) and stable isotope values (carbon and nitrogen; see Table S8) for seven individuals. The analyses were performed at the Curt-Engelhorn-Zentrum Archäometrie gGmbH (Mannheim, Germany). The collagen used for the analyses was extracted from fragments of the petrous bones used for palaeogenomic analysis. The isotope results were used as a basis for palaeodietary inference (see Fig. S18) and for freshwater reservoir effect correction in the case of the Vlasac sample (Table S7). Radiocarbon dates on human bones from archaeological sites in the Danube Gorges are known to be influenced by a large freshwater $^{14}C$ reservoir effect, due to high intake of freshwater food, and need to be corrected before calibration (*100*). This was done here following the method described by Cook *et al. (100),* later amended by Bonsall *et al. (101)*: any Danube Gorges date with associated $\delta^{15}N>8.3‰$ is corrected using a weighted mean age offset of 545 ± 70 radiocarbon years for 100% fish-based diet (corresponding to $\delta^{15}N = 17‰$).

For the palaeodietary reconstruction, we analyzed the delta values of carbon ($\delta^{13}C$) and nitrogen ($\delta^{15}N$)(*192*). Carbon isotopes are generally used to determine the amount of marine and terrestrial proteins in ancient diets, as well as $C_3$ and $C_4$ photosynthetic pathways. Marine endpoints of $\delta^{13}C$ are -12 ± 1‰ while values from terrestrial proteins centre around -21 ± 1‰ (*193*, *194*). Nitrogen isotopes are often used to determine the trophic level of the organism within a specific ecosystem and for distinguishing between aquatic and terrestrial food sources. Terrestrial herbivores $\delta^{15}N$ values typically range between 4 and 7‰, whereas carnivores - including humans - show values about 3-4‰ higher on average than the organisms they consume (*193*, *195*).

The $\delta^{13}C$ values of the seven individuals reported here range from -19.13‰ to -20.54‰, while the $\delta^{15}N$ values range from 8.73‰ to 15.77‰ (Table S8). In order to place the sampled individuals within a broader dietary context, we plotted their values together with published isotopic data from Mesolithic and Neolithic individuals of Central and Southeast Europe, Northwestern (NW) Anatolia, together with faunal remains from the same sites (Fig. S18).

The Neolithic individuals from Herxheim (Herx), Asparn-Schletz (Asp6) and Kleinhadersdorf (Klein7) cluster with other Neolithic individuals from Central Europe (Fig. S18). Without detailed examination of local food web variations, inferences made in this section remain tentative. The $\delta^{13}C$ values observed indicate a diet based on terrestrial animal protein with less plant contribution. Herx shows values suggestive of higher $C_3$ plant (e.g. wheat, barley, leafy vegetables) consumption, when compared with other sampled individuals. Asp6 presumably consumed more animal than plant proteins. The individual from Dillingen-Steinheim (Dil16) has lower $\delta^{15}N$ values compared to Central European Neolithic individuals, but overall falls within the range of distribution of Neolithic farmers in Central Europe. The lower $\delta^{15}N$ may be influenced by the quality of the terrestrial proteins, meaning that the proxy includes animals lower in the food chain, i.e. herbivores ($\delta^{15}N$ values typically +5.3 ±1.9). It is worth bearing in mind that nitrogen isotopic variations differ greatly within and between ecosystems and trophic levels; herbivore bone collagen values are affected by many factors including climate, organism's physiology *etc* (*196*). The Neolithic individual from Barcın (Bar25) has the lowest $\delta^{13}C$ value in the dataset and plots outside the Neolithic group. The sample's $\delta^{13}C$ intermediate value may be influenced by a mixed $C_3$ and $C_4$ (e.g. millet) plant consumption, as well as animal proteins. Nea3, the ~6-9 month old infant from Nea Nikomedeia has higher $\delta^{15}N$ value compared to other Neolithic individuals of this study, related to breastmilk consumption (breastfed infants typically have 2-3‰ higher $\delta^{15}N$ values than their mother (*197*, *198*)). The Mesolithic individual from the Danube Gorges (VLASA32) has the highest $\delta^{15}N$ value, and plots outside the range of published Mesolithic individuals from Central Europe (*199*). Its high $\delta^{15}N$ value suggests a large intake of freshwater food, primarily large fish (e.g. anadromous salmonids), and is consistent with recent published values from the Danube Gorges Mesolithic sites (*141*).

**Table S8 - Summary of the $\delta^{13}C$ and $\delta^{15}N$ values obtained from the 7 individuals analyzed in this study.** The values given are the nitrogen content (N [%], average of triple determination) with standard deviation (1σ; N [%] SD), the carbon content (C [%], average of triple determination) with standard deviation (1σ; C [%] SD), the atomic C:N ratio, the $^{15}N/^{14}N$ isotopic ratio (D/C $\delta^{15}N$ [‰ AIR]) relative to the standard AIR with drift correction (D/C) (average of triple determination) with standard deviation (1σ; D/C $\delta^{15}N$ [‰ AIR] SD) and the $^{13}C/^{12}C$ isotopic ratio (D/C $\delta^{13}C$ [‰ VPDB]) relative to the standard VPDB with drift correction (D/C) (average of triple determination) with standard deviation (1σ; D/C $\delta^{13}C$ [‰ VPDB] SD).

| Sample ID | Collagen yield [%] | N [%] | N [%] SD | C [%] | C [%] SD | C:N | D/C $\delta^{15}N$ [‰ AIR] | D/C $\delta^{15}N$ [‰ AIR] SD | D/C $\delta^{13}C$ [‰ VPDB] | D/C $\delta^{13}C$ [‰ VPDB] SD |
|---|---|---|---|---|---|---|---|---|---|---|
| VLASA32 | 2.76 | 10.71 | 0.69 | 29.71 | 1.94 | 3.24 | 15.77 | 0.12 | -20.13 | 0.09 |
| Bar25 | 7.07 | 15.97 | 0.08 | 43.79 | 0.09 | 3.20 | 10.20 | 0.01 | -19.13 | 0.04 |
| Nea3 | 5.18 | 12.87 | 0.79 | 35.37 | 1.88 | 3.21 | 11.88 | 0.03 | -19.91 | 0.07 |
| Klein7 | 1.00 | 10.07 | 0.29 | 28.38 | 0.80 | 3.29 | 9.65 | 0.02 | -20.31 | 0.06 |
| Dil16 | 2.76 | 13.57 | 0.23 | 36.95 | 0.72 | 3.18 | 8.73 | 0.08 | -20.52 | 0.02 |
| Asp6 | 1.16 | 7.17 | 0.97 | 21.54 | 2.81 | 3.51 | 10.01 | 0.14 | -19.97 | 0.07 |
| Herx | 6.75 | 14.39 | 0.13 | 39.32 | 0.16 | 3.19 | 10.28 | 0.02 | -20.54 | 0.01 |

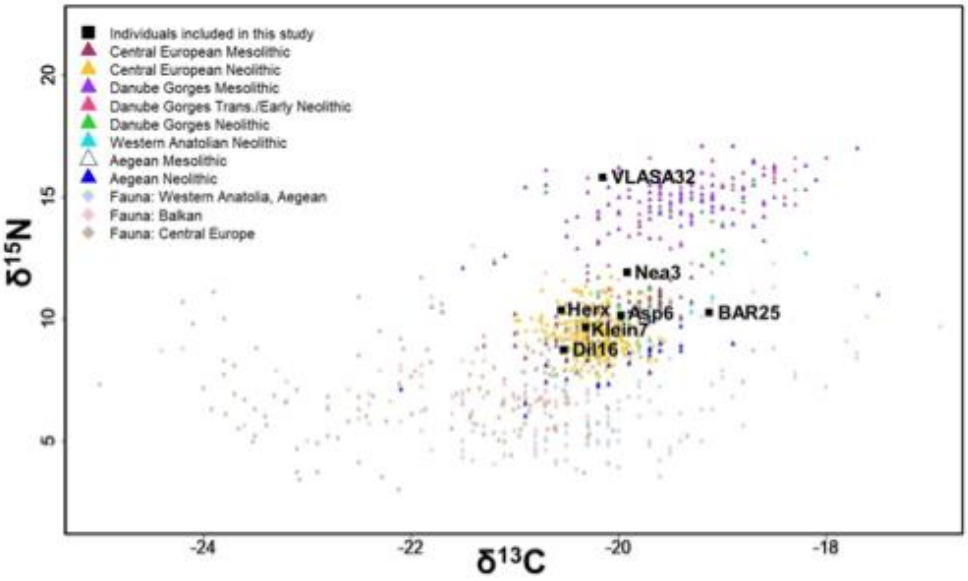


**Figure S18 - Scatterplot of $\delta^{13}C$ and $\delta^{15}N$ values obtained from the individuals analyzed in this study and published data** from 613 Neolithic and Mesolithic humans from Central Europe (*199*), the Danube Gorges (*16*, *100*, *101*, *103*, *104*, *141*, *177*, *200–203*), the Aegean Basin (*16*, *204*, *205*) and NW Anatolia (*16*, *152*, *153*). Published faunal values from the same period/regions are included for comparison (*103*, *141*, *152*, *153*, *177*, *206*, *207*).

### *Genetic structure analysis*

#### Multi-Dimensional Scaling (MDS) on pairwise average nucleotide divergence

Contrastingly to the analyses performed only on neutrally evolving sites (Fig. 2A), a MDS analysis done on the whole genome including sites potentially affected by selection (Fig. S19A; *102samples* dataset) reveals a slightly different picture as it suggests strongest affinities of European Neolithic farmers with modern individuals from Southern Europe other than Sardinians (Crete, Greece, Italy, Albania, Spain), which is at odds previous analyses based on the projections of ancient individuals on Principal Components computed from modern individuals only (*4*). It also suggests some genetic continuity since at least Neolithic times as the early Neolithic individual from Iran (WC1) and the Caucasus HG (KK1) show strongest genetic affinities with modern Iranians and individuals from the Northern Caucasus. Note that the MDS performed on ancient genomes only (Fig. S19B; *Ancient* dataset) shows more differences between WC1 and KK1 than when moderns are also included.

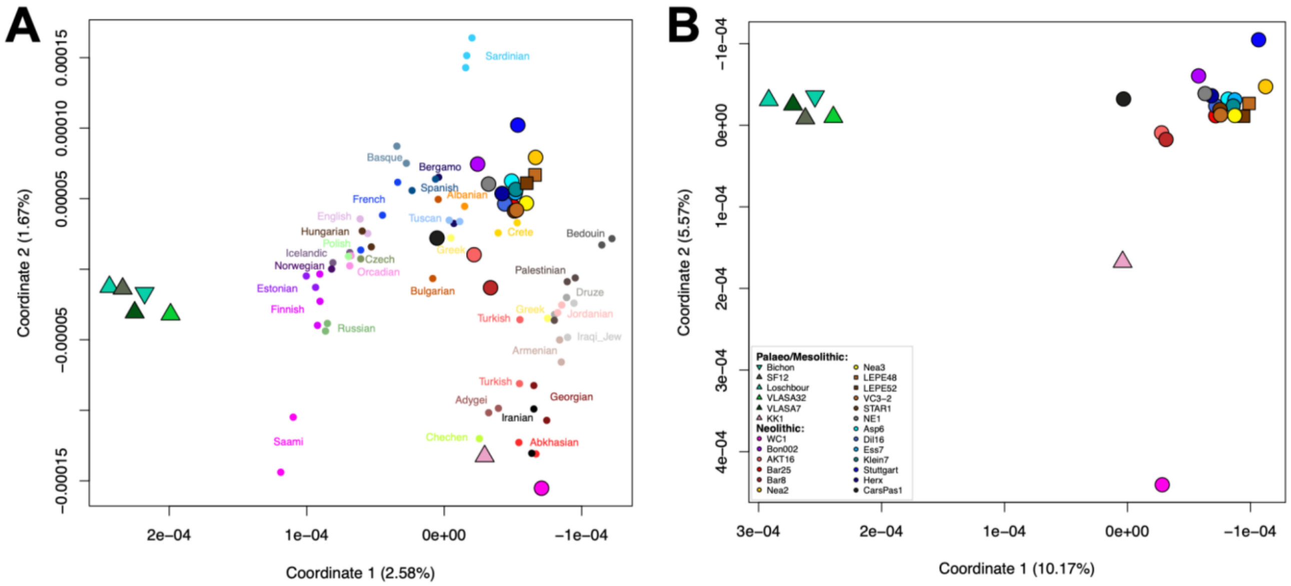


**Figure S19 - 2D-MDS performed on the average nucleotide divergence ($\pi_{XY}$) matrix computed over the whole genome** of (**A**) all Europeans and SW Asians (n = 90; *102samples* dataset), ancient and modern (shown with small circles); (**B**) ancient individuals only (n = 25; *Ancient* dataset).

### Runs of Homozygosity (ROHs)

We detected short (2-10 Mb) and long (>10 Mb) segments of homozygosity-by-descent or Runs of Homozygosity (ROHs) from the imputed genomes of the European and SW Asian modern and ancient individuals (Fig. 2D, Fig. S20). The number and total genome size made up of the short ROHs (Fig. S20A) are high in ancient genomes, especially those of HG individuals in keeping with previous results (*10*, *208*, *209*) and of a few early Neolithic individuals (Bon002, WC1 and LEPE52; STAR1 and AKT16 to a lesser extent), as well as of some modern individuals from Europe and the Near East. These ROH are expected to be negatively correlated with population size, so that a larger number of short ROHs indicate remote inbreeding and potentially smaller population sizes. We indeed found a negative correlation between the cumulative length of these short ROHs and the expected neutral heterozygosity observed in ancient genomes (Fig. S20C; Spearman' rho test = -0.5, p-value < 0.01).

For the long ROHs indicative of recent inbreeding between close relatives (potentially second cousins or closer (*210*), some modern individuals present the larger amount and some ancient genomes to a lesser degree (WC1 and LEPE52 are inferred to be the most consanguineous ancient individuals, followed by Stuttgart, Bichon and Loschbour). The three other HGs were not found to have long ROHs nor Bon002 and AKT16. For modern individuals, most of the consanguineous individuals have both short and long ROHs but not necessary (e.g. Turkish-2 and Russian-1 exhibit long ROHs but only a few short ROHs, respectively 3 and 0 Mb in short ROHs).

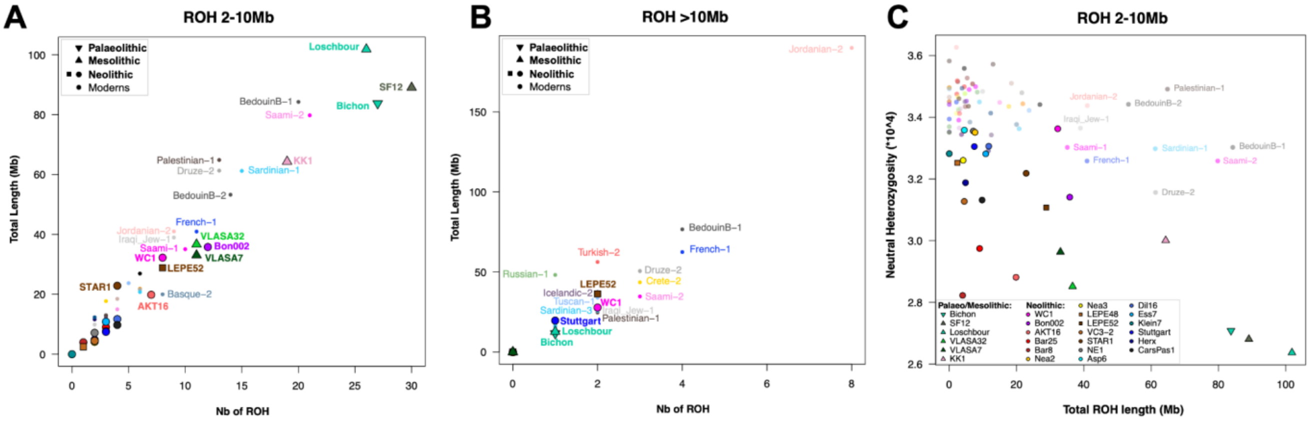


**Figure S20 - ROHs found in the imputed European and SW Asian genomes** (n = 90). Number and cumulated length of (**A**) short ROHs (size between 2 and 10Mb) and (**B**) long ROHs (>10Mb); (**C**) Neutral expected heterozygosity as a function of the cumulative length of short ROHs.

### Sex-specific markers and haplogroups

Consistently with the other individuals from the Danube Gorges published to date (*16*, *211*), the two newly sequenced HGs from Vlasac site were found to carry mitochondrial haplogroup U5 (subclade U5a2a), the most common haplogroup in the Danube Gorges during Mesolithic times. Their Y-chromosomal haplogroups (VLASA32: R1b1, VLASA7: I2) also fall within the known diversity as, to date, all Mesolithic individuals from this area exclusively show I or R (R1b) haplogroups (*16*, *211*).

In contrast, Y-chromosomal haplogroups of the newly sequenced Neolithic males either belong to haplogroup G (Asp6: G2a2b2a3, Bar25: G2a2b2a1, Ess7: G2a2b2a1a1, LEPE52: G2a2b2a1a1c, VC3-2: G2a2a1a3~) or haplogroup C (LEPE48: C1a2b, Dil16: C1a2b). While haplogroup G was common among early Neolithic farmers (*16*), haplogroup C was less frequent in comparison (G: 56.52%, C: 17.39%, estimated based on version v42.4 of https://reichdata.hms.harvard.edu/pub/datasets/amh_repo/curated_releases/V42/V42.4/SHARE/public.dir/v42.4.1240K.anno).

Mitochondrial haplogroup frequencies observed in the 13 newly sequenced Neolithic individuals were consistent with those previously reported from a Neolithic context: we found five individuals with haplogroup K (Nea2: K1a, Nea3: K1a2c, LEPE48: K1a1, Herx: K1a4a1i, AKT16: K1a3), one individual each with haplogroups N1a1a1 (Bar25), J1c6 (Dil16), W1-119 (Klein7), T2e2 (STAR1) and HV-16311 (VC3-2). Furthermore, U5 haplogroups, common among HGs (*16*) and increasing in frequencies during the middle and late Neolithic periods, were found in early Neolithic individuals from Southern Germany and Lower Austria (Ess7: U5b2c1, Asp6: U5a1c1). Additionally, haplogroup H3 was inferred for an individual from the early-Middle Neolithic in the Danube Gorges (LEPE52); this haplogroup is rare or even absent in early Neolithic individuals but is found more frequently in the Middle Neolithic in Germany (*21*, *212*) as well as the Iberian Peninsula (*5*, *213*, *214*). While it was previously suggested that H3 haplogroup was associated with a glacial Iberian refugium and has spread throughout Europe from there (*215*), our data indicates that H3 was also already present in Neolithic Serbia.

### Neanderthal introgression

As a result of interbreeding between Neanderthals and modern humans some time after the out-of-Africa migration, all non-African individuals derive a small portion of their genome from Neanderthals. We estimated the Neanderthal ancestry proportions in the ancient individuals used in *fastsimcoal2* demographic modelling (Fig. S21) as described in the Methods section. Besides elevated values in Bon002, Bichon, and Bar8, proportions are approximately 4%. This may be expected as all analysed individuals share a common ancestor ~25kya and are linked by a number of admixture events. The high ancestry proportions inferred in Bon002, Bichon, and Bar8 are likely artifacts of lower quality genomes. In line with previous results by (*75*, *216*), we find no significant difference in Neanderthal ancestry levels between ancient hunter-gatherers and Neolithic individuals (permutation test *p*-value = 0.711). (Fig. S22).

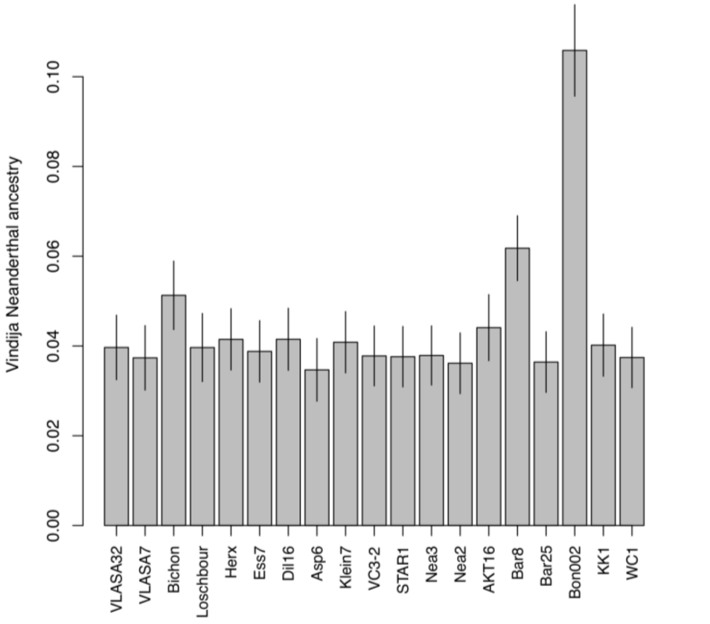


**Figure S21 - Inferred Vindija Neanderthal ancestry proportions for ancient individuals** (n = 19; individuals used to select and fit the *fastsimcoal2* demographic modelling).

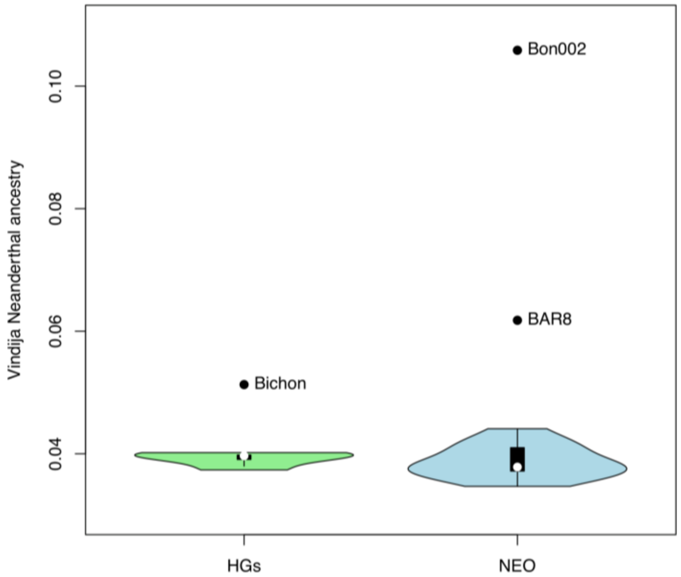


**Figure S22 - Comparison of Vindija Neanderthal ancestry proportions** between ancient hunter-gatherers and Neolithic individuals (see Fig. S21 for individual estimates).

### ***Phenotypic analysis***

#### Pigmentation

With the HirisPlexS webtool (*64*, *65*), we managed to predict pigmentation phenotypes for the 15 newly sequenced ancient individuals, except for eye-color in VLASA32 for which no allele at the SNP rs12913832 could be retrieved.

For some individuals, one or several SNPs in the *MC1R* and TUBB3 genes associated with hair color and skin pigmentation (*217*, *218*) were missing in the VCF. As described in the Material & Methods, these SNPs had to be looked up in the BAM files and we considered they could be in homozygous or heterozygous state. This resulted in some predictions of red hair (Dil16, Ess7, Herx, Klein7). Given the overall low frequency of the derived alleles associated with light pigmentation phenotypes in European populations (*219*, *220*) as well as in ancient data (*5*), a red haired phenotype is highly unlikely and we considered as an artifact.

We found that the vast majority of the newly sequenced early Neolithic individuals most likely had an intermediate to light skin complexion, while the two Mesolithic individuals were inferred to have darker skin tone in comparison. A dark (brown to black) hair color was inferred for all but two individuals: LEPE52 and VC3-2 had more likely a light brown phenotype. Eye color variation was similarly low, with the majority of individuals showing highest probabilities for brown eyes, except for two farmers of the Neolithic Starčevo culture (STAR1 and VC3-2) who were likely blue-eyed. Thus, the highest phenotypic variation in our dataset seems to originate from Serbian individuals.

**Table S9 - Phenotypes for eye and hair color as well as skin tone, inferred from HirisPlexS and their probabilities for the 15 newly sequenced individuals.** The resulting HirisPlexS input and results file can be found in Supp. Table 3.

| Individual | BAM SNPs[a] | Eye Color | p(Eye Color)[b] | Hair Color | p(Hair Color)[b] | p(Shade)[b] | Skin Tone | p(Skin Tone)[b,c] |
|---|---|---|---|---|---|---|---|---|
| AKT16 | rs885479, rs12821256, rs4959270, rs3114908, rs2238289, rs1126809, rs1545397, rs8051733 | Brown | pBrown(0.993); pBrown(0.993) | Dark Brown | pBrown(0.387), pBlack(0.586); pBrown(0.420), pBlack(0.543) | pD(0.910); pD(0.800) | Intermediate/ Pale | pInt(0.776); pP(0.563), pInt(0.410) |
| Asp6 | rs885479, rs1805008, rs2228479, | Brown | pBrown(0.996); pBrown(0.996) | Dark Brown to Black | pBrown(0.387), pBlack(0.586); pBrown(0.420), | pD(0.910); pD(0.800) | Intermediate/ Pale | pInt(0.776); pP(0.563), pInt(0.410) |

| | | | | | | | | |
|---|---|---|---|---|---|---|---|---|
| | rs1110400, rs2378249, rs1426654 | | | | pBlack(0.543) | | | |
| Bar25 | rs683, rs3114908, rs2238289 | Brown | pBrown(0.954); pBrown(0.954) | Dark Brown to Black | pBrown(0.426), pBlack(0.550); pBrown(0.408), pBlack(0.573) | pD(0.916); pD(0.936) | Intermediate to Dark | pInt(0.799); pInt(0.560), pD(0.395) |
| Dil16 | rs11547464, rs885479, rs1805008, rs1805007, rs1110400, rs4959270, rs1393350, rs683, rs1126809, rs1470608, rs1545397, rs8051733 | Brown | pBrown(0.938); pBrown(0.921) | Dark Brown to Auburn | pBrown(0.600), pBlack(0.311); pRed(0.995) | pD(0.809); pL(1.000) | Pale/ Intermediate to Dark | pInt(0.455), pD(0.512); pVP(0.444), pP(0.385) |
| Ess7 | rs11547464, rs885479, rs1805008, rs1805007, rs1110400, rs2238289, rs1126809 | Brown | pBrown(0.997); pBrown(0.997) | Dark Brown to Auburn | pBrown(0.332), pBlack(0.663); pRed(0.966) | pD(0.994); pL(1.000) | Pale/ Dark | pD(0.972); pVP(0.658), pP(0.129) |
| Herx | rs11547464, rs885479, rs1805008, rs1805007, rs1805009, rs2228479, rs1110400, rs683, rs1800414, rs2238289, rs1126809, rs1545397 | Brown | pBrown(0.986); pBrown(0.986) | Dark Brown to Auburn | pBrown(0.604), pBlack(0.261); pRed(1.000) | pD(0.817); pL(1.000) | Pale/ Intermediate to Dark | pInt(0.507), pD(0.436); pVP(0.671), pP(0.270) |
| Klein7 | rs885479, rs1805008, rs1805007, rs1110400, rs4959270, rs10756819, rs2238289, rs1126809, rs6059655 | Brown | pBrown(0.938); pBrown(0.938) | Dark Brown to Auburn | pBrown(0.538), pBlack(0.370); pRed(0.865) | pD(0.658); pL(0.938) | Pale/ Intermediate | pInt(0.760); pP(0.792) |
| LEPE48 | rs885479, rs1805008, rs10756819, rs2238289 | Brown | pBrown(0.997); pBrown(0.997) | Black | pBlack(0.729); pBlack(0.778) | pD(0.984); pD(0.971) | Intermediate | pInt(0.784); pInt(0.913) |

| | | | | | | | | |
|---|---|---|---|---|---|---|---|---|
| LEPE52 | rs885479, rs1805008, rs2238289, rs17128291, rs1126809, rs1470608 | Brown | pBrown(0.767); pBrown(0.767) | Light Brown | pBlond(0.298), pBrown(0.514), pBlond(0.357), pBrown(0.427); | pL(0.593), pD(0.407); pL(0.727) | Intermediate | pInt(0.651), pD(0.146); pP(0.530), pInt(0.258) |
| Nea2 | rs1805009, rs2228479, rs12203592, rs2238289, rs1126809 | Brown | pBrown(0.989); pBrown(0.979) | Dark Brown/ Black | pBrown(0.484), pBlack(0.497); pBrown(0.513), pBlack(0.468) | pD(0.980); pD(0.996) | Intermediate | pP(0.198), pInt(0.508); pInt(0.289), pD(0.683) |
| Nea3 | rs885479, rs12203592, rs2238289, rs17128291, rs1126809, rs1545397 | Brown | pBrown(0.968); pBrown(0.941) | Dark Brown/ Black | pBrown(0.386), pBlack(0.598); pBlack(0.787) | pD(0.952); pD(0.994) | Intermediate | pInt(0.554), pD(0.386); pP(0.190), pInt(0.657) |
| STAR1 | rs2378249, rs2238289, rs1126809 | Dark Blue | pBlue(0.605), pBrown(0.296); pBlue(0.605), pBrown(0.296) | Brown/ Dark Brown | pBrown(0.480), pBlack(0.430); pBrown(0.540), pBlack(0.367) | pL(0.523), pD(0.477); pL(0.475), pD(0.525) | Pale/ Intermediate | pP(0.389), pInt(0.535); pInt(0.709) |
| VC3-2 | rs1393350, rs3114908, rs2238289, rs1126809, rs1470608, rs1545397, rs8051733 | Blue | pBlue(0.784); pBlue(0.836) | Light Brown | pBlond(0.303), pBrown(0.523); pBlond(0.303), pBrown(0.523) | pL(0.790); pL(0.790) | Pale/ Intermediate | pP(0.294), pInt(0.683); pP(0.642), pInt(0.279) |
| VLASA32 | rs885479, rs1805008, rs1110400, rs1800407, rs1126809, rs1545397 | NA | NA | Dark Brown/ Black | pBrown(0.406), pBlack(0.588); pBlack(0.768) | pD(0.989); pD(0.983) | Intermediate/ Very Dark | pVD(0.983); pInt(1.000) |
| VLASA7 | rs1805008, rs2238289, rs6497292 | Brown | pBrown(0.998); pBrown(0.998) | Dark Brown/ Black | pBrown(0.319), pBlack(0.678); pBrown(0.391), pBlack(0.602) | pD(0.997); pD(0.992) | Intermediate/ Dark | pInt(1.000); pD(0.821) |

[a] SNPs for which the allele was taken from the BAM file, because they were missing in the VCF.

[b] For each phenotype two independent sets of probabilities are given, separated by a semicolon: the first were obtained assuming that all the BAM SNPs were homozygous for the found allele and the second assuming a heterozygous state. If per set the highest probability was not larger than 0.7, the two highest values are reported separated by a comma.

[c] (D = Dark, VD = Very Dark , Int = Intermediate, L = Light, P = Pale, VP = Very Pale).

### Additional phenotypic variation

Lactase persistence

All individuals investigated in this study had the ancestral G allele at rs4988235, the variant in the *MCM6* gene of which the derived allele is highly associated with lactase persistence into adulthood in Eurasia (*221*). Since no derived alleles were found, it is unlikely that any of the individuals was able to digest lactose, consistently with an increase in frequency of the lactase-persistence allele at a later stage (*222*).

EDAR/ABCC11

We analyzed two SNPs associated with incisor shape/hair thickness (rs3827760, *EDAR*; derived alleles are associated with thick straight hair and shovel shaped incisors (*223*)) and earwax type/body odor (rs17822931, *ABCC11*; derived alleles at rs17822931 are associated with dry earwax and reduced body odor (*224*)), both having high derived alleles frequencies in modern East-Asian populations (*225–227*). All ancient individuals presented here only had ancestral alleles for both SNPs.

Fatty acids synthesis

We further investigated seven SNPs located in the *FADS1/2* gene complex, reported to have been selected for in various populations (*228*, *229*). Variation in *FADS1/2* is known to influence the ability to synthesize omega-3 and omega-6 long-chain polyunsaturated fatty acids (LC-PUFAs) from precursor molecules (*229*, *230*). Since LC-PUFAs are also found in food, especially from animal sources, the dietary shift during the Neolithic transition is hypothesised to have created a selective pressure that lowered diversity at *FADS* associated loci.

Genotypes for each SNP were looked up in the filtered VCF. Individuals were grouped by subsistence into hunter-gatherers and Neolithic farmers (excluding WC1, KK1 and the Lepenski Vir individuals, see Table S10). A two-sided binomial test was used to compare counts of derived alleles in our ancient individuals with frequencies in a modern population. We used the Central Europeans from Utah (CEU, N = 99) from the 1000Genomes project as a proxy for a modern Central European population.

For rs74771917, while the derived T allele is almost lost in CEU (0.030), it is found in significantly larger numbers in farmers (3/26 = 0.115, $p<0.043$). No differences could be found between CEU and farmers for any of the other SNPs. For HGs, the allele counts suggest lower frequencies for the SNPs rs174546_C ($f_{CEU} = 0.641$; HG: 1/10 = 0.100, $p<0.001$), while higher frequencies in comparison to CEU are inferred for rs174570_T ($f_{CEU} = 0.162$; HG: 3/4 = 0.750, $p<0.015$) and rs174594_A ($f_{CEU} = 0.616$, HG: 10/10 = 1.000, $p<0.009$). For rs174594_A, counts suggest a higher frequency in HGs compared to farmers (Farmer: 17/28 = 0.607, HG: 10/10 = 1.000, p<0.008). A similar result was found for rs97384_C (Farmer: 12/26 = 0.462, HG: 7/8 = 0.875, p<0.028).

These results are consistent with selection affecting variation in *FADS1/2* already during the early Neolithic (*5*), rather than starting later in the Bronze Age as proposed more recently (*229*). This is because, with the exception of one SNP, frequencies in farmers are close to those in modern populations, but multiple SNPs differ in frequencies between HGs and modern populations. While it is possible that migration and admixture during and after the Late Neolithic led to a shift in frequencies at *FADS* related loci, our observations are also consistent with selective pressure associated with the Neolithic diet having shaped allele frequencies in early Neolithic populations to approach the levels observed in modern populations today.

**Table S10 - Derived allele frequencies for seven SNPs located in the FADS1/2 gene complex** for modern Europeans (CEU) and counts for the derived (p) and total number of alleles for the ancient samples, grouped by subsistence ($n_{HG}$ = 5, $n_{Farmers}$ = 16).

| SNP | Derived Allele | CEU derived frequency | No. of derived alleles | |
|---|---|---|---|---|
| | | | Farmers | HGs |
| rs174546 | C | 0.641 | 12/24 | 1/10 |
| rs174570 | T | 0.162 | 8/28 | 3/4 |
| rs174594 | A | 0.616 | 17/28 | 10/10 |
| rs97384 | C | 0.606 | 12/26 | 7/8 |
| rs74771917 | T | 0.030 | 3/26 | 1/8 |
| rs174455 | A | 0.647 | 13/20 | 5/10 |
| rs174465 | T | 0.702 | 7/12 | 4/8 |

Polygenic Score for Height

Height is a classical polygenic trait, which is highly heritable, but also strongly influenced by environmental variables (*66*). Hundreds of variants are known to influence height in humans, with the majority alleles having only a tiny effect (*231*). To measure the distribution of height-related alleles, we computed generalized polygenic scores (PS) for standing height using a set of 670 SNPs (*67*) for 20 ancient individuals (Fig. S23).

The highest scores were found for the two individuals from Vlasac, Serbia (VLASA32: 20.033, VLASA7: 20.147), while the lowest score was obtained for the individual from Herxheim, Germany (Herx: 18.086). We found a strong positive correlation of individual height PS with the average age of the samples (Pearson's r = 0.77577, p<6e-05). The correlation remained strong when considering only Neolithic individuals between 8,300 and 7,000 BP, both with (Pearson's r = 0.6537, p<0.0082) and without the Lepenski Vir individuals (Pearson's r: 0.6565, p<0.0148), suggesting that decreasing standing height was under selection during the Neolithic expansion along the Danubian corridor.

These findings are consistent with previous studies reporting selection for decreasing predicted height in early Neolithic populations from Southern and Central Europe (*5*) as well as modern populations from Sardinia (*232*), but differ from a morphometric analysis of prehistoric skeletons (*233*). Considering the high amount of ancestry attributed to early Neolithic farmers in modern Sardinians (*6*), these signals may be related. Although we find significant differences between mean scores for Mesolithic and early Neolithic individuals including Lepenski Vir (Student t-test, t = 2.8257, p<0.0112), these differences could be explained by the older date (except for Loschbour and WC1) of the Mesolithic individuals in comparison. The decrease in height PS values could be a continuation of a trend of decreasing stature between the Upper Palaeolithic and the Mesolithic (*234*), a result supported both by genetics and physical measurements; however, (*234*) did not find any differences between Mesolithic and Neolithic individuals.

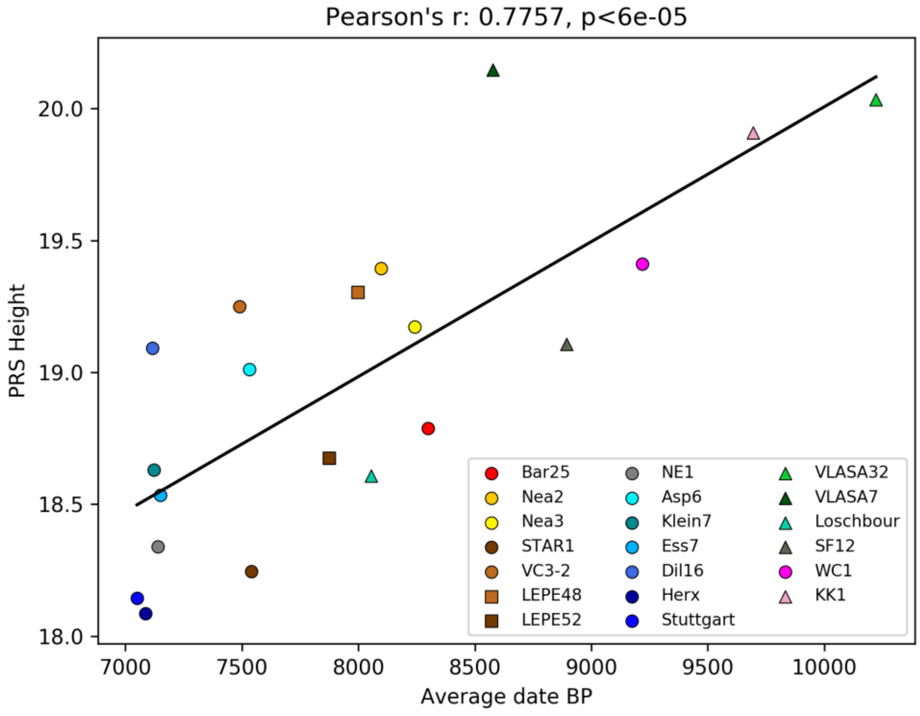


**Figure S23 - Scatter plot of the average age in BP vs. height PS values for each individual** (n = 20; Bichon, Bon002, Bar8, CarsPas1, AKT16 being excluded based on genotype-filtering thresholds). The best-fit straight line illustrates the Pearson correlation ($r^2 = 0.6018$) between height PS and sample age, indicating a decline in PS values over time.

***Joint Distribution of Fitness Effects (DFE) analysis***

Selective constraints may differ between Neolithic and modern populations, due to changes in cultural and environmental contexts over time. To test for such differences in purifying selection, we fit a model of the joint distribution of fitness effects (DFE) of deleterious mutations to our data (*77*). To facilitate modeling selection, we first fit a simple demographic model to our synonymous mutation data in which the ancestors of modern Europeans diverged from the ancestors of our Neolithic samples (Fig. S24A). In our joint DFE model, the fitness effects of all mutations were allowed to change after this divergence, under a joint bivariate lognormal distribution (Fig. S24E). When we fit this model to our nonsynonymous mutation data, our best-fit estimate of the correlation between selection coefficients in ancient versus modern populations was 0.997. We thus conclude that selective constraint has changed little in European populations between the Neolithic era and today.

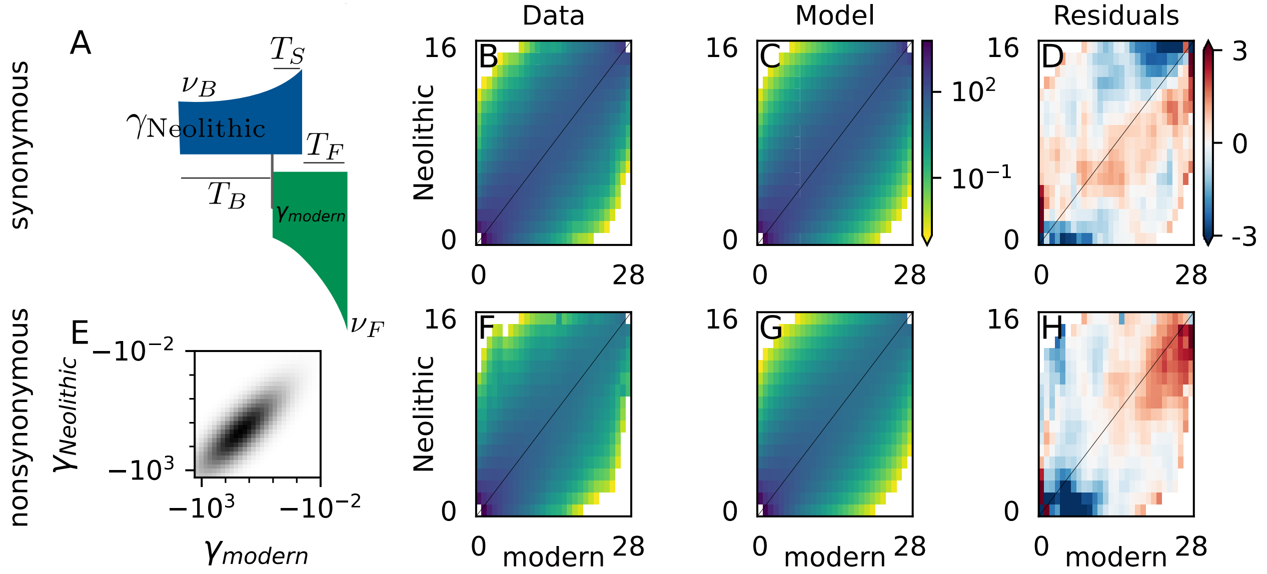


**Figure S24 - Joint Distribution of Fitness Effects (DFE) analysis.** (**A**) The demographic model that was fit to the synonymous data. In this model, the population from which the Neolithic genomes were sampled went through a bottleneck to relative size $n_B$ and then grew exponentially. The population from which the modern genomes were sampled diverged from that Neolithic population after $T_B$ time units and grew at the same rate to reach its final size $n_F$. The population from which Neolithic genomes were sampled diverged for $T_S$ time units until genomes were sampled $T_F$ time units ago. (**B-D**) The synonymous data and best-fit model allele frequency spectra, and the residuals between them. (**E**) Illustration of the bivariate lognormal DFE that was fit to the nonsynonymous data (shown with correlation 0.8). (**F-H**) The nonsynonymous data and best-fit model spectra, and the residuals between them.

***MSMC2 analysis***

**Inference of human population size**

The population effective size was estimated per sample (Fig. S25A) and per population (Fig. S25B) if more than one sample from the same region and period was available. The *MSMC2* results were similar for all non-African populations before 35 kya and included a deep bottleneck at roughly 50-60 kya, a pattern that has been reported previously (*235*). As expected, we were not able to get good recent estimates from a single diploid individual, although a clear difference in the population size between farmers and hunter-gatherers is observable. The resolution at recent times was much improved when using two individuals per population, in which case we found that all the Upper Paleo-Mesolithic HG populations and the population from the Zagros Region (WC1) exhibit a small population size of roughly $N_e$ = 3,000 up to 35 kya. In contrast, we estimated a constant increase of population size after 35 kya in all European and Anatolian Neolithic populations. This increase resulted in population sizes around $N_e$ = 10,000 - 15,000 about 10 kya, with NW Anatolia showing the smallest population size, North Greece, Lepenski Vir and Central Serbia showing similar sizes also around $N_e$ = 10,000 and smaller than the Austrian-German Neolithic populations (Lower Austria, Southern Germany1, Southern Germany2).

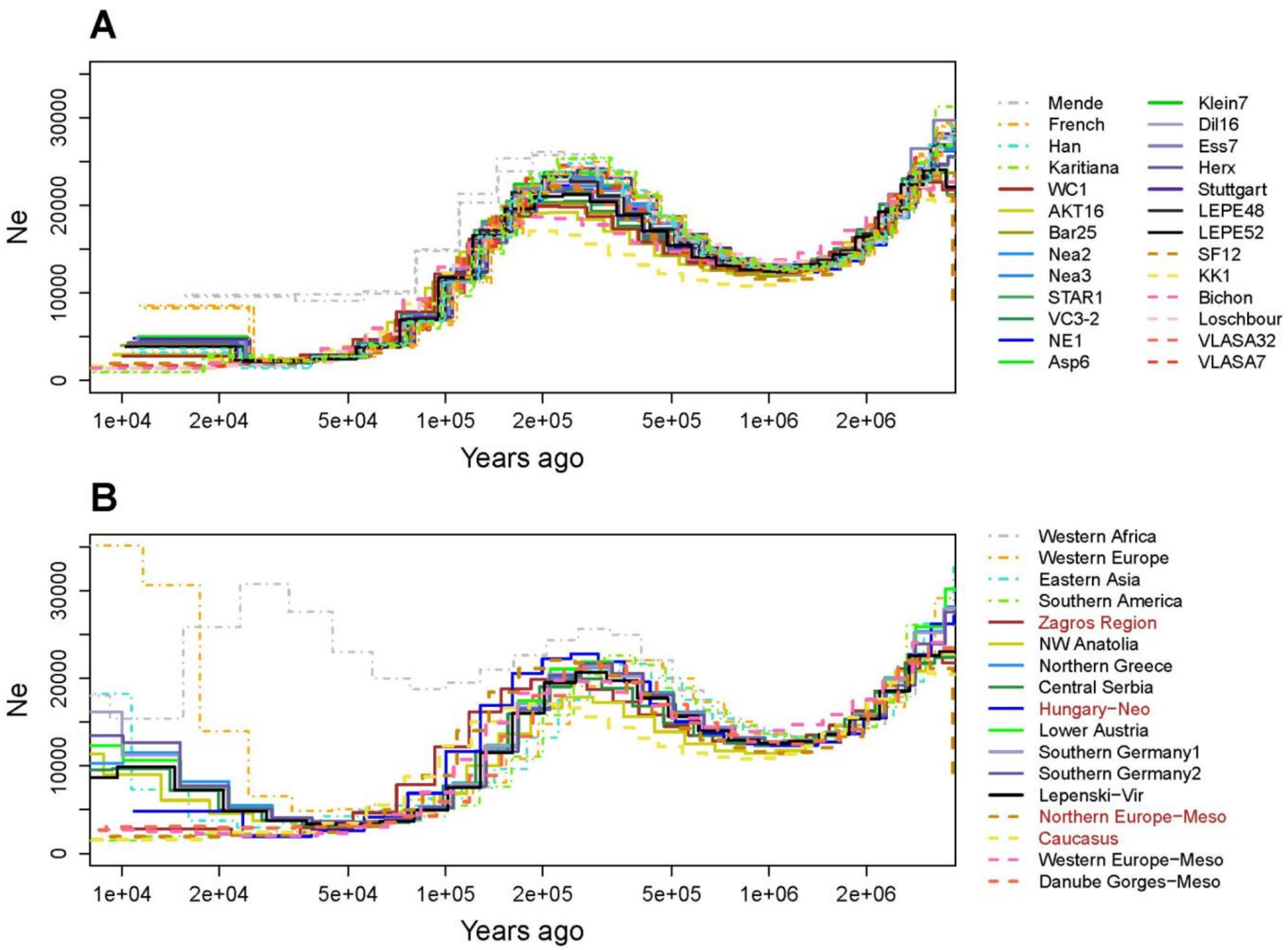


**Figure S25 - MSMC2 population size estimates scaled using a mutation rate of $1.25\times10^{-8}$ per generation per site and a generation time of 29 years.** (**A**) Past demography obtained for each individual (two haplotypes; $n_{ind}$ = 30). (**B**) Past demography obtained for populations for which two individuals (four haplotypes) were available (i.e. not for WC1, NE1, SF12 and KK1, who represent a unique group each; $n_{pop}$ = 17). Both analyses suggest smaller population sizes in the most recent times for HG populations compared to farmers.

**Divergence time between populations**

The relative cross coalescent rate (rCRR) was also calculated (Fig. S26) to estimate roughly the split times between populations (Fig. S27A-B). As a first observation, we note that split time estimates involving populations with a single sample (Northern Europe-Meso, Caucasus, Zagros Region and Hungary-Neo) tend to be much older. Indeed, CCR estimates from one-sample populations drop to zero more quickly than corresponding estimates from two-sample populations among Neolithic and Mesolithic populations. We therefore caution against interpreting the older split time estimates of Northern Europe-Meso from all other populations as strong evidence for the influence of Eastern HGs, which diverged from Western HGs at an earlier period.

Focusing on two-sample Neolithic populations, the estimated split times capture the old split time between Upper Palaeolithic-Mesolithic and Neolithic populations, and also show that all Neolithic populations split at roughly the same time. However, no clear order of split times emerges among the Neolithic populations, with large variations depending on the exact comparison. Among the most consistent results across several comparisons is a relatively old split of NW Anatolia, followed by populations from what is today Greece, the Balkans and finally Austria-Germany, in accordance with a stepwise migration between neighbouring regions. Lepenski Vir for instance, split first from NW Anatolia and Northern Greece (~24 kya), then from Central Serbia (~21 kya) and from the Austro-German Neolithic populations (~19-22 kya). But we note that several individual estimates are at odds with this sequence of events: NW Anatolia, for instance, was inferred to have split from Lower Austria and Lepenski Vir (~24 kya) earlier than from Northern Greece and Central Serbia (~22 kya). Similarly, Northern Greece split from NW Anatolia earlier than from Central Serbia and Southern Germany1 (~21 kya). Our CCR results may thus not provide sufficient resolution at this scale. In addition, they are likely affected by recent admixture. The relatively old split time we estimated between NW Anatolia and the other Neolithic populations, along with the more recent split time we estimated between NW Anatolia and the two Upper Palaeolithic-Mesolithic populations (Danube Gorges-Meso and Western Europe-Meso), might be well explained by recent HG admixture into NW Anatolia, in particular AKT16 (Fig. 2B).

Interestingly, the split time between the Danube Gorges-Meso and Western Europe-Meso is estimated to be the most recent (~15 kya) among all populations, suggesting they share more recent common ancestors. While this observation is in line with their clustering in our MDS analysis (Fig. S19), it is somewhat younger than estimates obtained with *fastsimcoal2* (see below), and potentially a result of the very estimates of their recent population sizes (and hence low intra-coalescent rates) and the uncertainty associated with cross-coalescent rates.

In line with the final demographic scenario inferred from SFS analyses and shown in Fig. 3, MSMC2 infers split times of ~22 kya among Neolithic populations and ~35 kya between Neolithic and Mesolithic samples (Fig. S27). We note, however, that Bichon and Loschbour were estimated to have split from the Danube Gorges Mesolithic samples more recently (~15 kya) than what is inferred from the site frequency spectrum (23.3 kya). These estimates obtained from the analysis of whole genomes are older than what is inferred with *fastsimcoal2* on the neutrally evolving sites only, but are not incompatible given that they assume no gene flow or admixture between samples. Previous analyses have also shown that while MSMC and SFS-based inference programs were giving very congruent results on simulated data, they could lead to inconsistent result when applied to real genomes (*236*), implying that the two approaches were differentially affected by genomic factors not included in their assumptions.

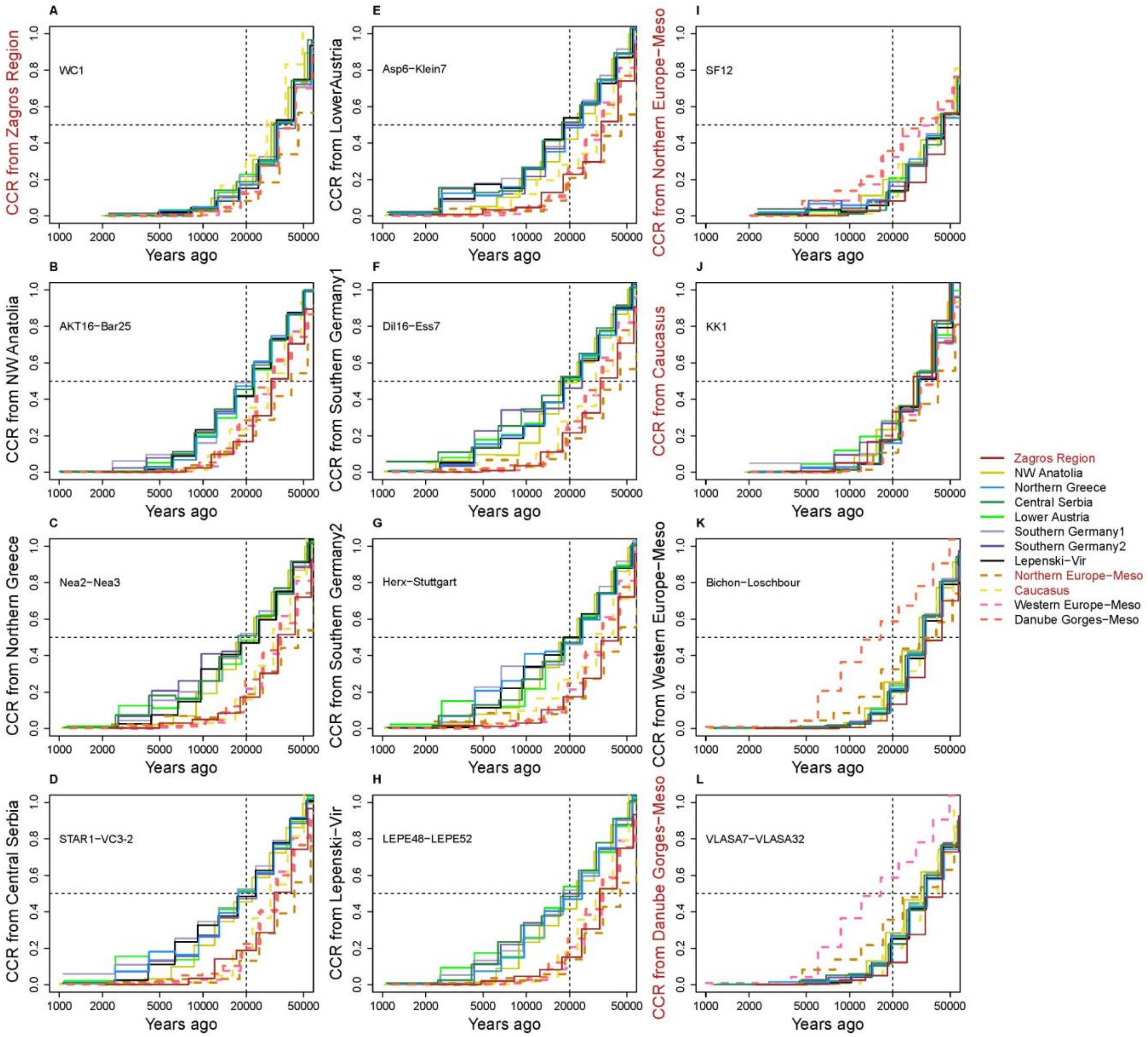


**Figure S26 - Relative cross coalescence rate (CCR) estimated with MSMC2 using pairs of individuals per population where available** (WC1, SF12 and KK1 are the unique representative of one population each). We estimated the relative CCR for all possible pairwise group combinations. The results were scaled using a mutation rate of $1.25{\times}10^{-8}$ per generation per site and a generation time of 29 years. Each subplot shows the pairwise CCR estimates of one population against all others. **First two columns**: Populations from the Neolithic period (A - G) and transitional and Neolithic layers at Lepenski Vir in the Danube Gorges (H). **Last column:** Populations from the Upper Paleo-Mesolithic period (I - L).

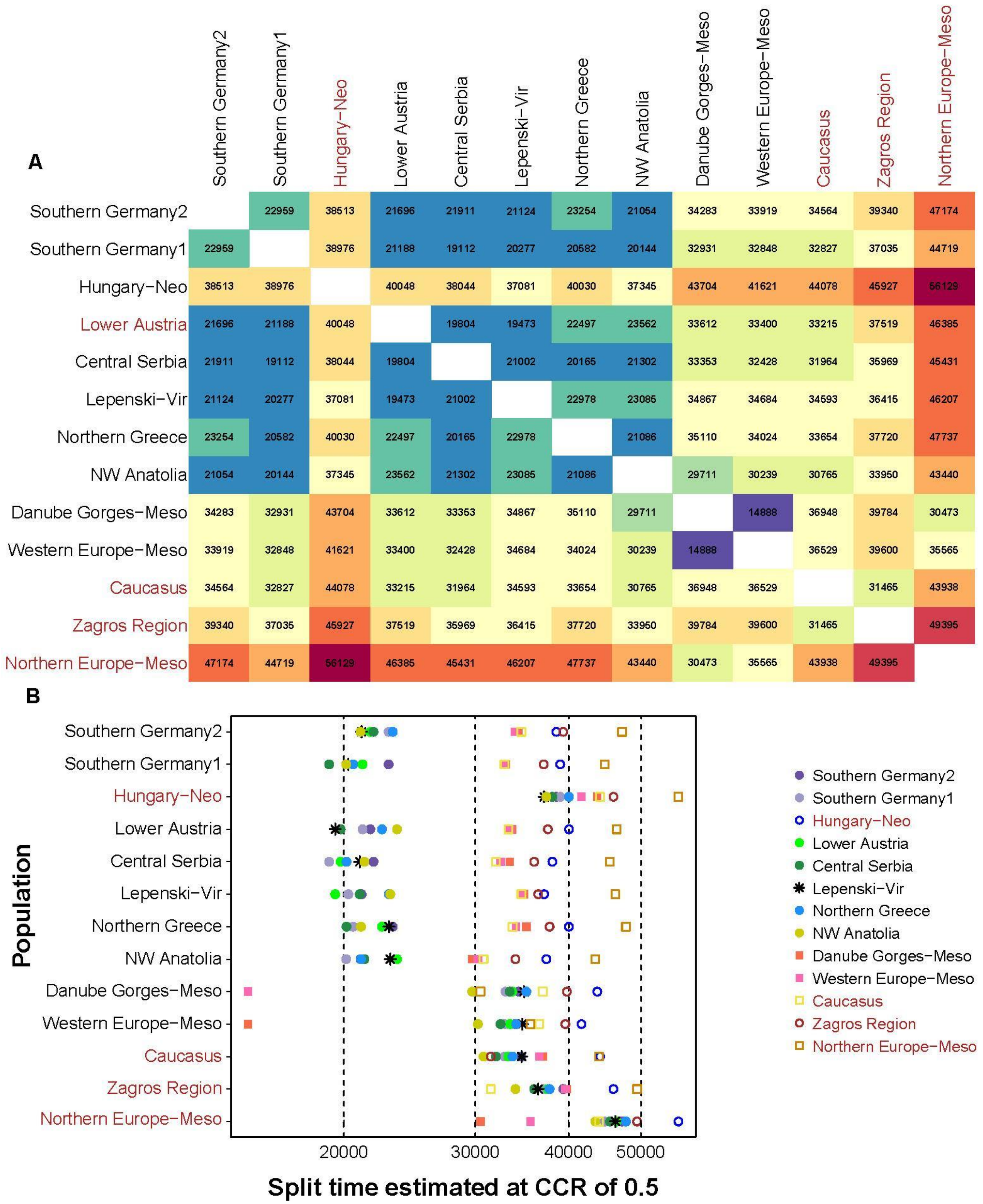


**Figure S27 - Divergence time estimates obtained with MSMC2 at the relative CCR of 0.5 for all pairwise comparisons**. (**A**) Square matrix showing the corresponding split time values between all pairs. (**B**) Point split time estimates for all pairwise comparisons. Populations with only one individual (Hungary-Neo, Caucasus, Zagros Region and Northern Europe-Meso) are marked in red and those comparisons are plotted using symbols with outlines only (not filled).

### Bootstrap analysis for the $N_e$ and CCR estimates

We show the bootstrap estimates together with the original estimates of the effective population size for all ancient populations (Fig. S28) as well as the bootstrap estimates and original values of the relative CCR between NW Anatolia and all other ancient populations (Fig. S29). We observed that there is little uncertainty around the estimates of the original dataset based on the bootstrap estimates, although the uncertainty increases for the most recent times (after 10 kya), for the cases where there are fewer haplotypes per run, or samples of lower depth (Fig. S29), as expected (*80*).

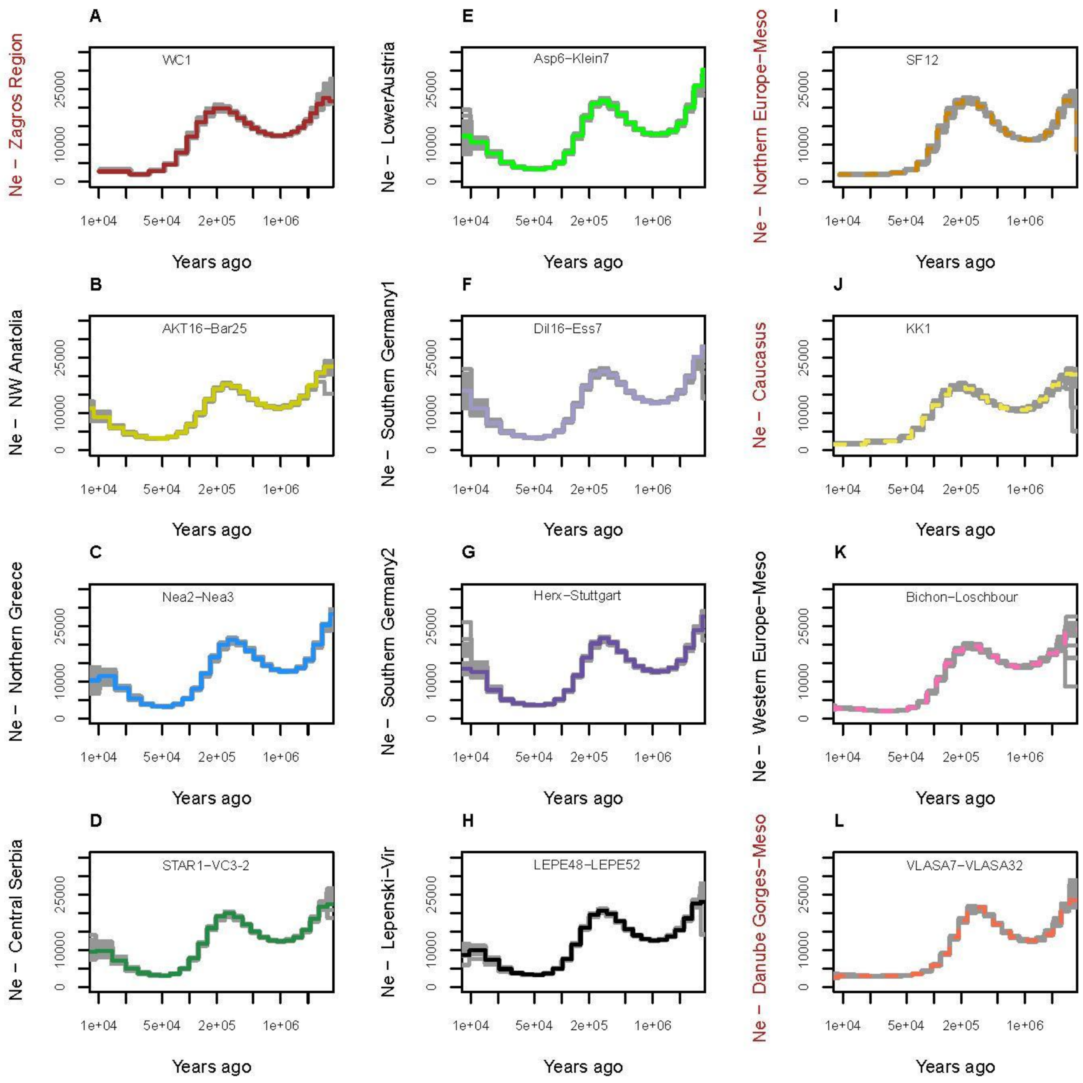


**Figure S28 - Variation in estimates of past effective sizes for all ancient populations obtained from 20 artificial data sets per population generated by bootstrapping 5Mb blocks. First two columns**: Populations from the Neolithic period (A - G) and transitional and Neolithic layers at Lepenski Vir in the Danube Gorges (H). **Last column:** Populations from the Upper Paleo-Mesolithic period (I - L).

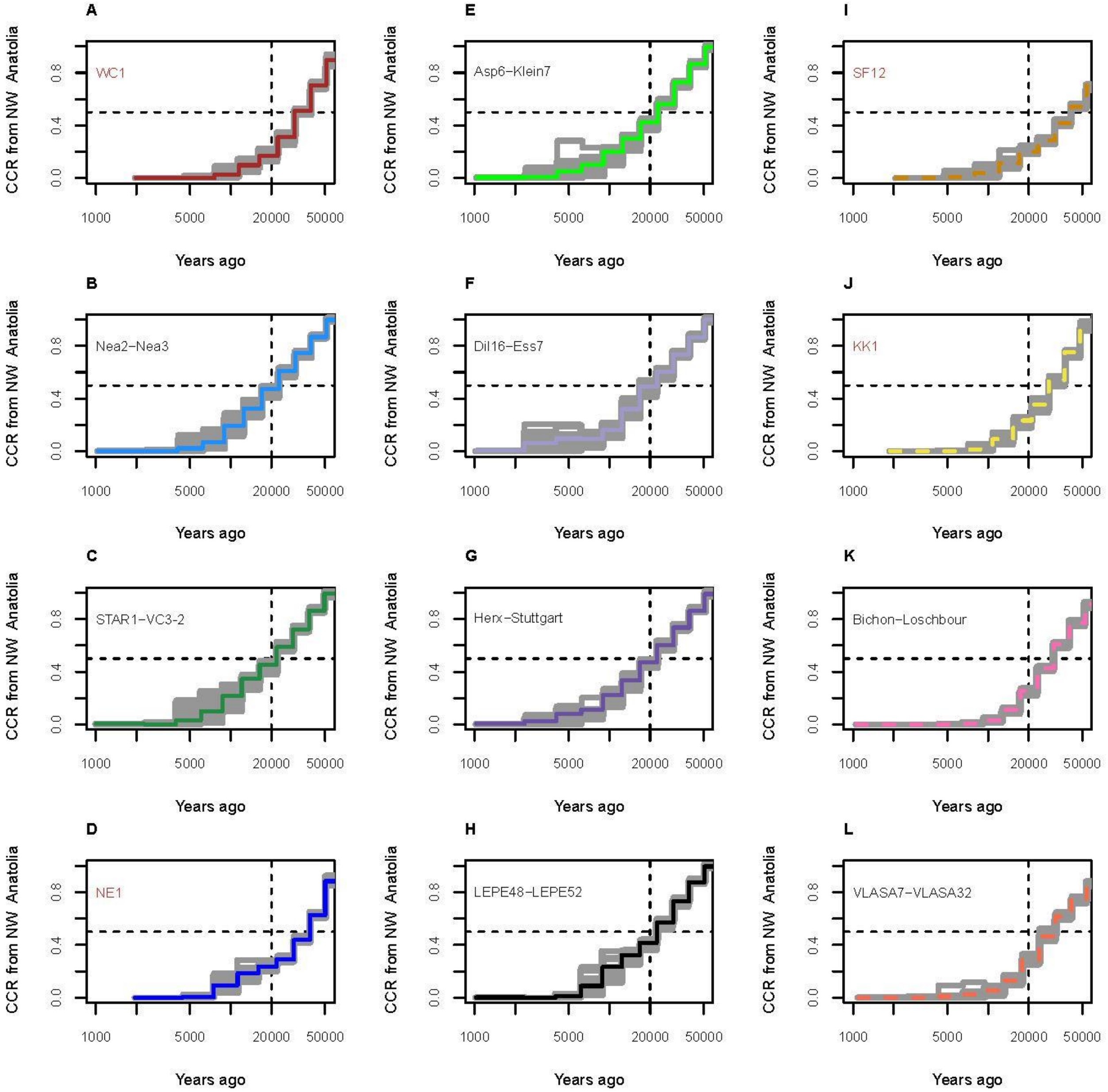


**Figure S29 - Variation in estimates of coalescence rates (CCR) for all comparisons involving NW Anatolia and obtained from 20 artificial data sets per population generated by bootstrapping 5Mb blocks. First two columns**: Populations from the Neolithic period (A - G) and transitional and Neolithic layers at Lepenski Vir in the Danube Gorges (H). **Last column:** Populations from the Upper Paleo-Mesolithic period (I - L).

***Demogenomic inference with fastsimcoal2***

**Framework**

Given the relatively high depth (>10X) of our newly sequenced individuals, it was possible to perform demogenomic inference on ancient genomes in a way similar to what is done on modern individuals with *fastsimcoal2 (58)*. By computing the site frequency spectrum (SFS) on unascertained polymorphic "neutral" sites, we could evaluate the likelihood of various historical scenarios and infer parameter values under these models. We nevertheless had to make several adjustments to take into account the temporal and spatial heterogeneity of the collected samples. First, in order to perform population level analyses, we had to pool geographically and culturally similar individuals into the same population, even though they would have been living centuries apart (Table S4). Since this temporal heterogeneity can lead to a Wahlund effect (*237*) that translates into a faster rate of coalescence between the two homologous gene copies of an individual relative to two gene copies from different individuals, we introduced population inbreeding coefficients in our modeling as nuisance parameters to specifically account for this population subdivision effect (*238*). Second, we introduced unsampled populations from which sampled populations could receive migrants, to reflect the fact that human populations are seldom isolated when not living on islands, which has a direct effect on the shape of the SFS within populations (*239*) and could thus introduce biases in demographic inferences if not properly taken into account. These unsampled populations can also be considered as a set of populations (a metapopulation) surrounding sampled populations (i.e. other unsampled farmer populations still exchanging genes with sampled farmer populations, or surrounding hunter-gatherer (HG) groups contributing to the early farmer gene pool by admixture) (*240–242*). Note also that we have modeled gene flow between metapopulations and sampled populations as a single pulse for modeling convenience, even though gene flow might have been continuous over several generations.

### Demographic models and parameter estimation

Relationship between major groups

*Tested models*

In order to study the genetic relationships between the populations that occupied SW Asia and Southern Europe before the Neolithic period, we have investigated the fit of data for several models (Fig. S30). The data consisted of the multidimensional unfolded SFS computed on the *HGNeo* panel with samples representative of the four groups that are clearly distinguishable on the two first axes of the MDS analysis (Fig. 2A, Fig. S19): two genomes from the Danube Gorges Mesolithic population to represent the HG cluster, the early Neolithic Iranian genome, two genomes from the Northern Greece Neolithic population from the Neolithic cluster and one from Mesolithic Caucasus that was found to be close but divergent from the Neolithic cluster.

In all tested models, we assumed that the sampled populations belonged to larger pools of unsampled populations from which they diverged and could receive some migrants (we modeled such migrations as single pulses of gene flow occurring 10 generations before the populations were sampled). Here we model these sets of large and unsampled populations as metapopulations (called *MetaWest*, *Central* and *East*), sometimes described as ghost populations (*240*, *241*) or continents (*57*, *242*). We assumed that the *Eastern* and *Western* metapopulations diverged first, and then the *Central* metapopulation split from the *Eastern*. We explored alternative scenarios in which the *Central* metapopulation could receive some gene flow from the *Western* metapopulation, modeled as a single pulse after the divergence events.

Based on the MDS analyses (Fig2A, Fig. S19) and geographic information, we considered the Danube Gorges Mesolithic population to be related to the *Western* metapopulation, the Neolithic population from the Zagros region closer to the *Eastern* metapopulation, and the Neolithics from Northern Greece related to the *Central* metapopulation. One class of models (A) assumed that the Caucasus Mesolithic population related to Kotias KK1 genome was closer to the *Central* metapopulation (i.e. as a sister population of the Greek Neolithic population), while in model B), the Caucasus Mesolithic population was closer to the *Eastern* metapopulation and to the early Neolithic Iranians from the Zagros Region as it is classically described (*9*).

Furthermore, while A) and B) models include some population size changes in the three metapopulations, we explored a model C) without any bottleneck as done in former GPhocs analyses (*11*, *14*). Finally, we explored a climatic change inspired model D) in which the *Eastern* and *Western* metapopulations were constrained to split before the Last Glacial Maximum (LGM) and to undergo a bottleneck 23 kya during the LGM.

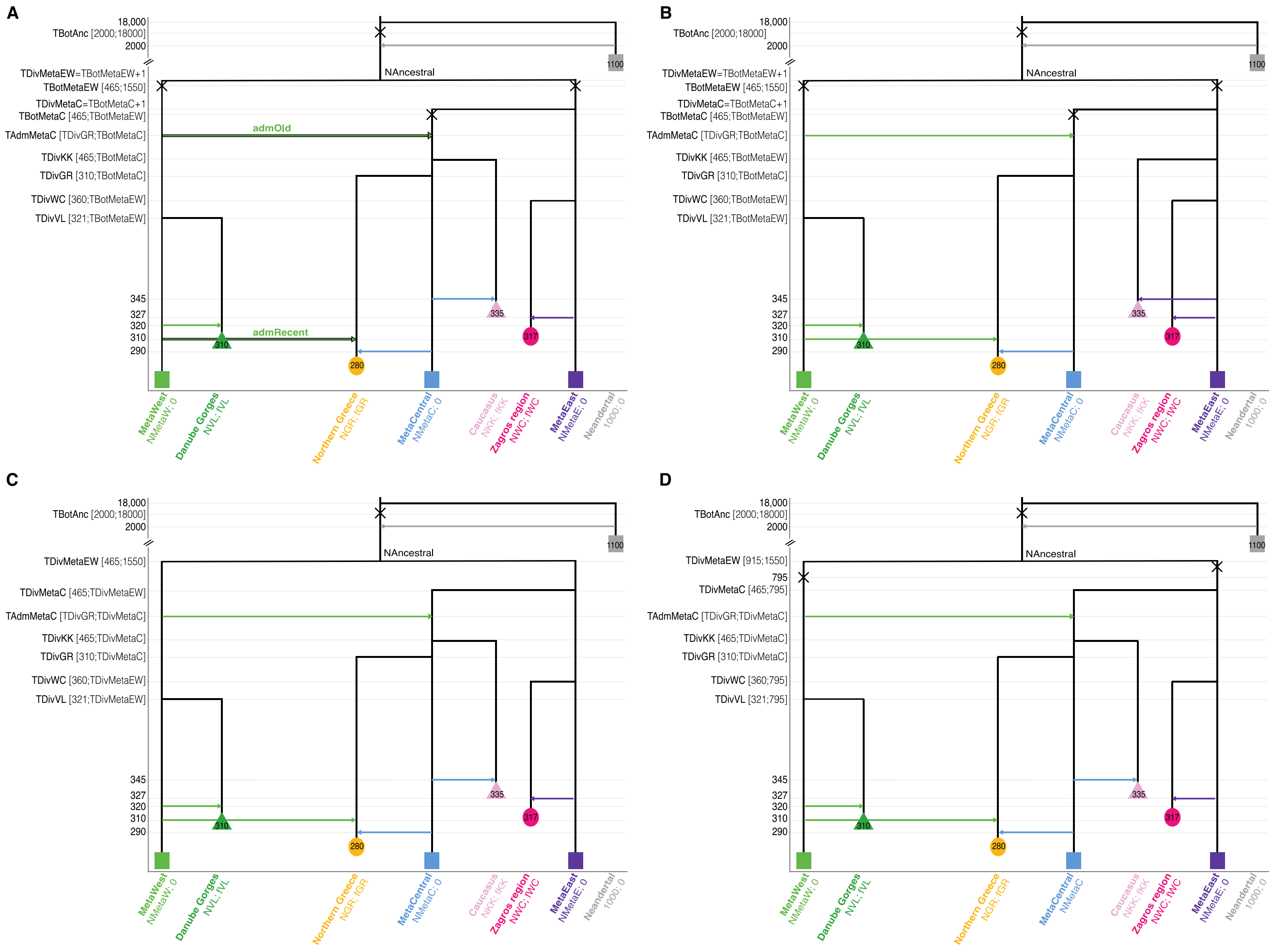


**Figure S30 - Schematic description of models tested for the *HGNeo* panel.**

(**A**) **Mesolithic Caucasus population branching on the *Central* metapopulation.** The *Eastern* and *Western* metapopulations diverged TDivMetaEW generations ago; the *Central* metapopulation split TDivMetaC generation ago; the divergences are followed by bottlenecks (X) one generation later to model founder effects. The Mesolithic population from the Danube Gorges then diverged from the *Western* metapopulation TDivVL generations ago; the Neolithic population from the Zagros region diverged from the *Eastern* metapopulation TDivWC generations ago; the Caucasus Mesolithic and the Northern Greek Neolithic population diverged from the *Central* metapopulation respectively TDivKK and TDivGR generations ago. Each sampled population received some admixture from its source metapopulation 10 generations before its sampling time (indicated in the geometric shape representing the population). Three admixture scenarios were tested: i) no admixture from the *Western* metapopulation toward the *Central* metapopulation and to the Northern Greece Neolithic population, ii) an admixture “admOld” from the *Western* to the *Central* metapopulation TAdmMetaC generations ago, iii) both the “admOld” admixture between the *Western* and *Central* metapopulations and a more recent admixture from the *Western* metapopulation directly into the Northern Greek population 310 generations ago which corresponds to the onset of farming in this area (*123*, *134*). All migration events are modeled as single pulses of gene flow.

(**B**) **Mesolithic Caucasus population branching on the *Eastern* metapopulation.** Same as (A) with

both “admOld” and “admRecent” migrations authorized but the Caucasus Mesolithic population branched off directly from the *Eastern* metapopulation.
(**C**) **No bottlenecks.** Same as (A) with both “admOld” and “admRecent” migrations allowed, but without bottleneck in the metapopulations.
(**D**) **Resize during the LGM.** Same as (A) with both “admOld” and “admRecent” migrations allowed, but the divergence between the *Eastern* and *Western* metapopulations had to occur before the LGM and these two metapopulations undergo a bottleneck during the LGM 23 kya (= 795 generations).
On each figure, we show the parameters to estimate with their search ranges within brackets. Further information about the parameters can be found in Supp. Table 4 and the properties of the sampled populations in Table S4.

*Best model parameters estimates and SFS fit*

Within the tested models, we find that the model of class A) with both an old admixture from the *Western* metapopulation toward the *Central* metapopulation and a recent admixture toward the Northern Greece Neolithic population is best supported (Fig. S31A, Supp. Table 4), with excellent fits between the observed and estimated SFS (Fig. S31C).

In this model, the population sizes of the Mesolithic populations from the Danube Gorges and Caucasus are in the range of the Neolithic population sizes. However, there is a significant difference in the bottlenecks modeled as lasting a single generation occurring on the metapopulations branches: the bottlenecks on the *Eastern* and *Central* metapopulation branches are relatively weak (~50 individuals) while the ancestral bottleneck and the bottleneck occurring on the *Western* metapopulation branch are stronger (~6 individuals [4-10]).

For the split times inferred in this model, we find a deep divergence between the *Eastern* and *Western* metapopulations (~25.6 kya [17.3-31.3]) and a more recent divergence between the *Central* and *Eastern* metapopulations (~15.8 kya [14.3-25.6 kya]). However, these divergences are much younger than the previously inferred divergence time between Iranian and European early farmers (46-77 kya) (*11*) or that between European early farmers and Western European HGs (27-76 kya) (*14*), both estimations obtained under a simple model with constant population sizes and no admixture. For our model *C*, which does not include any bottleneck in the metapopulations, we obtain older divergence time (32 kya [26.6-37.4]) between the ancestors of *Western* and *Eastern* metapopulations, but the fit is much worse (Fig. S31A, Supp. Table 4). The model *D* in which *Western* and *Eastern* metapopulations are constrained to split before the Last Glacial Maximum (LGM) also leads to an older divergence time between these metapopulations (32.7 kya [28.7-41.8]), but it is less well supported than the best model *A_admOld&Recent* (Fig. S31A, Supp. Table 4). However, despite some imposed old divergence, its fit is better than *C* as the *Western* and *Eastern* metapopulations undergo a bottleneck 23 kya during the LGM. Therefore, it suggests that population size changes are important in this model.

The divergence of the sampled populations from their source metapopulation are relatively old but have quite broad confidence intervals: Danube Gorges Mesolithic (16.0 kya [9.7-24.5]); Iranian Neolithic (13.6 kya [11.0-24.6]); Northern Greece Neolithic population (13.5 kya [9.6-21.7]). The Caucasus Mesolithic population is found to have diverged from the *Central* metapopulation about 14.2 kya [13.7-19.0]: the scenario where Caucasus HGs are connected to the *Central* metapopulation (model *A*) is better supported than the one where they are directly connected to the *Eastern* metapopulation (model *B*). This split from the *Central* metapopulation happens just after the ancestral population received some admixture from the *Western* metapopulation (14% [8-26]). Therefore, both the Greek Neolithic and the Caucasus Mesolithic populations descend from the ancestral admixture between the *Western* and *Central* metapopulations. Indeed, models *A_admOld&Recent* and *A_admOld* in which the Caucasus Mesolithic receives some *Western* metapopulation admixture are more likely than the model without migration *A_noadm* or *B* (Fig. S31A, Supp. Table 4). In addition to this ancestral admixture, the Greek population seems to have received an extra admixture from the *Central* metapopulation (about 15% [11-24]).

Interestingly, our best model requires a significant amount of gene flow (5-11%) from metapopulations to sampled populations, showing that human populations are seldom isolated (*239*), also validating the need to model surrounding unsampled populations.

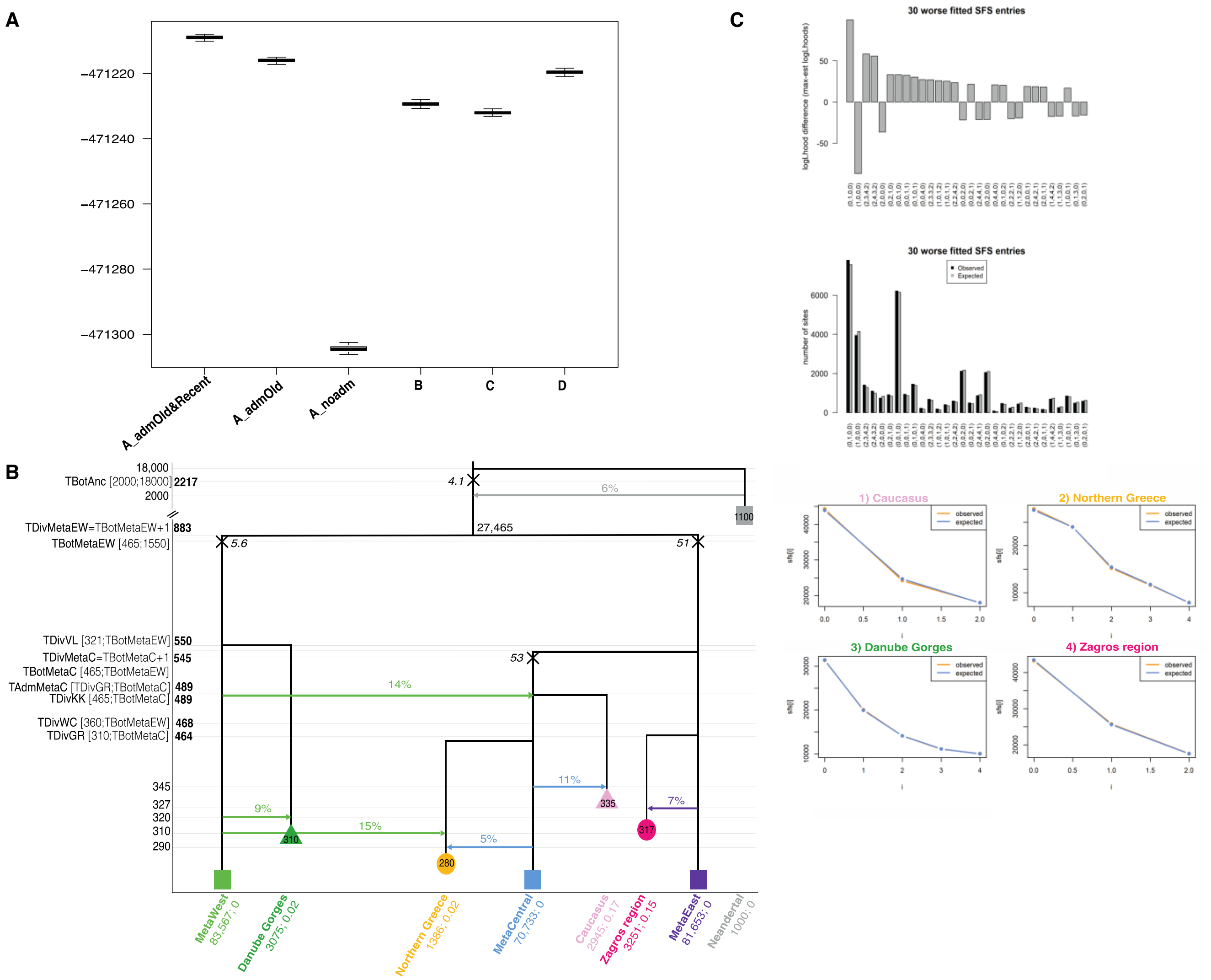


**Figure S31 - Exploration of models best explaining the observed multi-dimensional *HGNeo* SFS**

(**A**) **Distribution of the log10-likelihoods** obtained by simulation from the maximum likelihood point estimates for the six tested models described in Fig. S30. Higher likelihoods (top of the plot) indicate better fit.

(**B**) **Schematic representation of the ML demographic model *A_admOld&Recent*.** Values of various parameters are reported with their 95% CI estimated from 100 parametric bootstraps. Close to the bottleneck symbol (X), we report the size of a bottleneck, conveniently modeled as a single generation bottleneck (*instbot fastsimcoal2* option) .

(**C**) **Fit between the observed and estimated SFS under the ML model** (multidimensional SFS on the upper panel; marginal SFS for each population on the bottom).

### Neolithic Aegean structure

*Tested models*

In a second step, we extended the best model inferred from the *HGNeo* panel by including an individual from NW Anatolia (*Aegean* panel) in order to estimate the divergence times between the sampled Aegean populations, i.e. from Greece and NW Anatolia. We tested two classes of models: A) models in which an ancestral Aegean population would have diverged from the *Central* metapopulation and then split into NW Anatolian and Northern Greece Neolithic populations; B) a model in which NW Anatolian and Northern Greece Neolithic populations directly and separately diverged from the *Central* metapopulation, without the existence of an Aegean shared genetic structure (Fig. S32). Furthermore, in the class A), we authorized different admixture events from the *Western* metapopulation: one occurring directly in the Aegean ancestral population one generation after its founding and/or separate admixture events into the Northern Greek and NW Anatolian populations 310 generations ago (i.e. approximately when Neolithic was introduced in these areas and, therefore, when the Neolithic populations were founded (*123*)).

The divergence times, population sizes, inbreeding coefficients and admixture rates related to the Aegean populations were left to be freely estimated while the other parameters were inherited from the best model inferred for *HGNeo.*

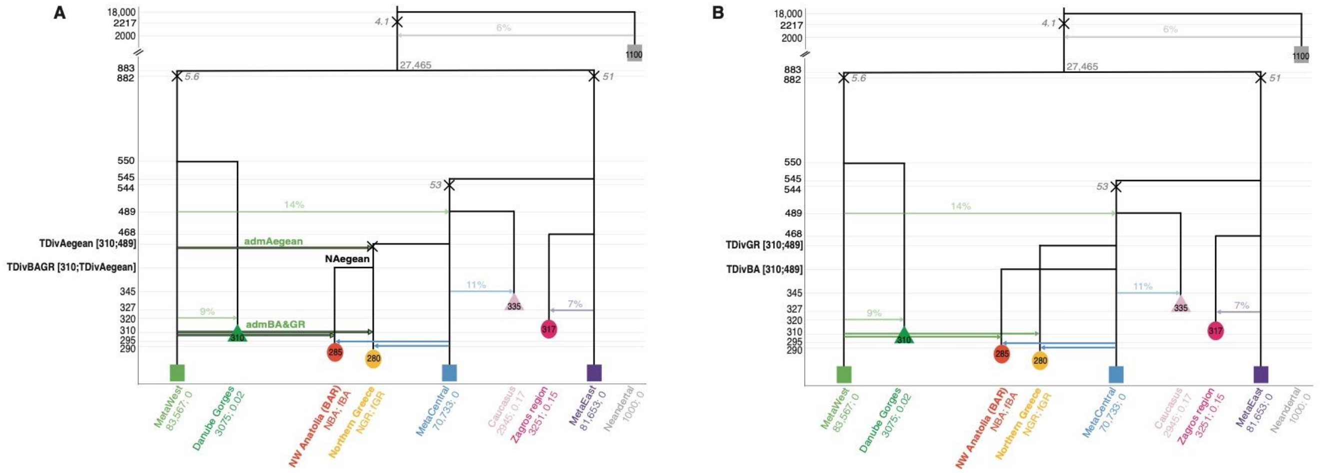


**Figure S32 - Schematic description of models tested for the *Aegean* panel.**

**(A) An ancestral Aegean structure.** Under this model, the Aegean ancestral population diverged TDivAegean generations ago from the *Central* metapopulation followed by an instantaneous bottleneck one generation after; then, the ancestral population split into Northern Greek and NW Anatolian populations TDivBAGR generations ago. We tested four scenarios: i) one allowing a single pulse of gene flow coming from the *Western* metapopulation to the Aegean branch one generation after its founding (*admAegean*), ii) one with independent admixture events into the Northern Greek and into NW Anatolia from the *Western* metapopulation 310 generations ago (*admBA&GR*), iii) one allowing both *admAegean* and *admBA&GR*, iv) one without any admixture.

**(B) Independent split events within the Aegean region.** Under this model, Northern Greece and NW Anatolia population independently split from the *Central* metapopulation TDivGR and TDivBA generations ago, and received independent pulses of gene flow from the *Western* metapopulation 310 generations ago.

As is the case for the Greek population, the NW Anatolia population received some gene flow from the *Central* metapopulation 10 generations before sampling. These models are derived from the best model inferred for *HGNeo*: parameter values estimated in the previous scenario were fixed in this analysis and shown in light fonts while newly estimated parameters are shown in bold fonts with search ranges shown within brackets. Further information about the parameters can be found in Supp. Table 4 and the properties of the sampled populations are listed in Table S4.

*Best model parameters estimates and SFS fit*

We find that all models of class A) very significantly outperformed model B), implying that the Neolithic populations from the Aegean region diverged from the same ancestral Aegean population (Fig. S33A, Supp. Table 4). This ancestral Aegean population received some gene flow from the *Western* metapopulation, as shown by the better fit of models *A_admAegean+BA&GR* and *A_admAegean* as compared to *A_admBA&GR*, and even more importantly than *A_noadm*.

Under model *A_admAegean+BA&GR,* which is the most complex and best supported scenario, the Barcın individual Bar25 from NW Anatolia seems drawn from a smaller population (2Ne = 1491 [358-5737]) than the Northern Greek population (2Ne = 4437 [1720-7689]), from which it would have diverged recently (320 generations ago, corresponding to ~9.3 kya [313-414]) (Fig. S33B, Supp. Table 4). The Aegean ancestral population is found to have been of small size (2Ne = 1242 [143-4300]), after a mild bottleneck (during one generation, 2Ne = 250 [9.5-333]) and to have diverged from the *Central* metapopulation 126 generations before (i.e. 12.3 kya [9.4-13.9]). Interestingly, this modeling allows us to refine the admixture from the *Western* metapopulation received by the Greek Neolithic found in *HGNeo*: it confirms that about 15% [6-17] of the gene pool of all Aegeans were received from the *Western* metapopulation (modeled here as a single pulse occurring when the Aegean ancestral population split from the *Central* metapopulation). There were also additional pulses of gene flow in the NW Anatolian Neolithic population (12% [6-16]) and in the Greeks (3% [1-11]) that might have contributed to their differentiation into distinct populations. Note that this approach of extending a former model with some fixed and newly estimated parameters leads to a very good fit between the observed and simulated SFS (Fig. S33C).

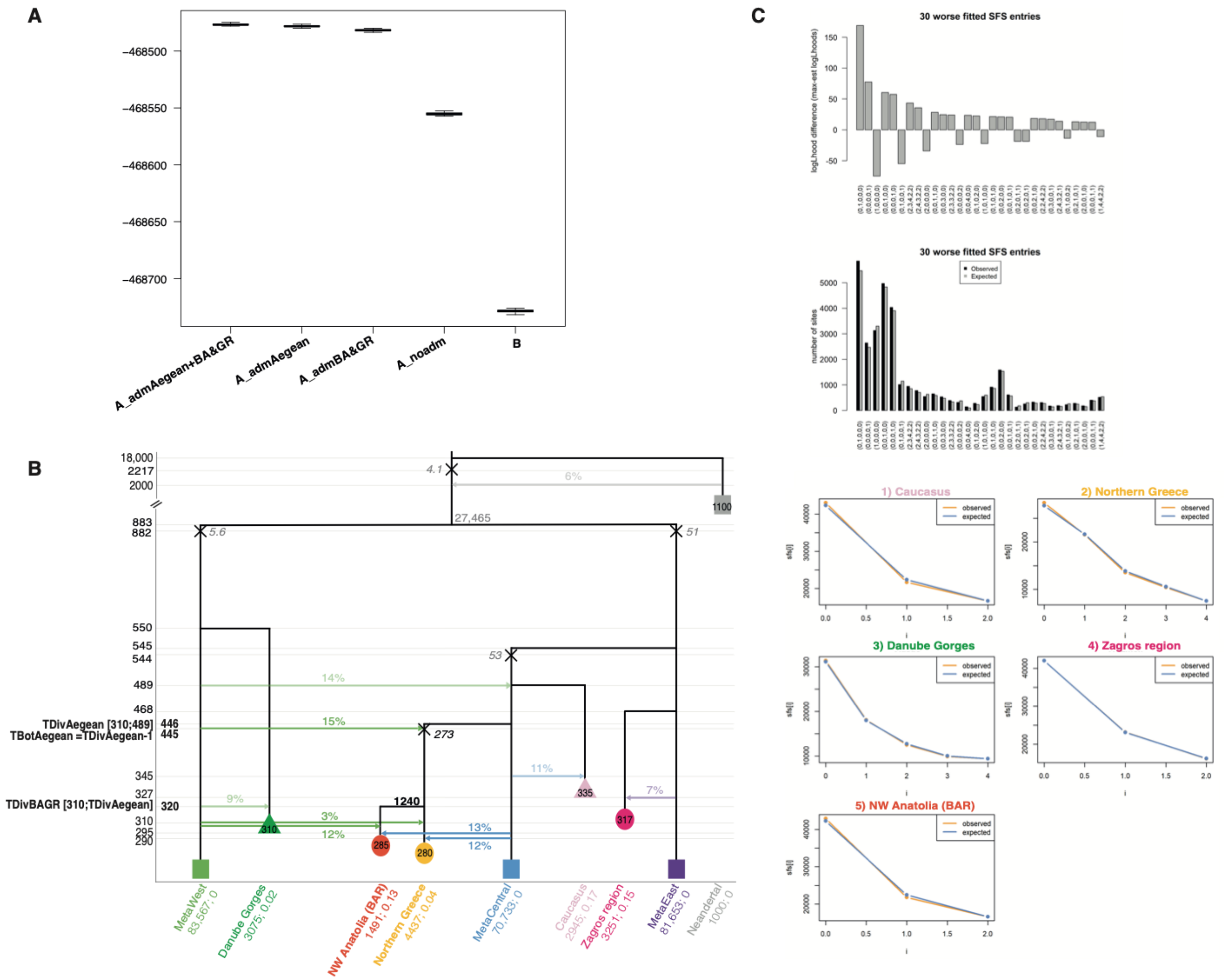


**Figure S33 - Exploration of models best explaining the observed multi-dimensional *Aegean* SFS**
(**A**) **Distribution of the log10-likelihoods** obtained by simulation from the maximum likelihood point estimates for the four tested models described in Fig. S32. Higher likelihoods (top of the plot) reflect better fit.
(**B**) **Schematic representation of the ML demographic model *A_admAegean+BA&GR*.** Values of various parameters are reported with their 95% CI estimated from 100 parametric bootstrap estimations. Close to the bottleneck symbol (X), we report the size of a bottleneck, conveniently modeled as a single generation bottleneck (*instbot fastsimcoal2* option) .
(**C**) **Fit between the observed and estimated SFS under the ML model** (multidimensional SFS on the upper panel; marginal SFS for each population on the bottom).

### NW Anatolia genetic structure

*Tested models*

In a third step, we used the *Aegean* best model parameters (Fig. S32B) as a backbone to examine more complex models involving an additional sample from the site of Aktopraklık, which in addition to being one of the oldest Neolithic sites in NW Anatolia, belongs to the local 'Fikirtepe culture', thought to have been influenced by both Mesolithic and Neolithic traditions (*20*). In order to investigate the possibly complex settlement history of this site, we tested four models allowing AKT16 to branch off from the NW Anatolian (BAR), from the Greek Neolithic population, from the ancestral Aegean population or from the *Central* metapopulation (Fig. S34).

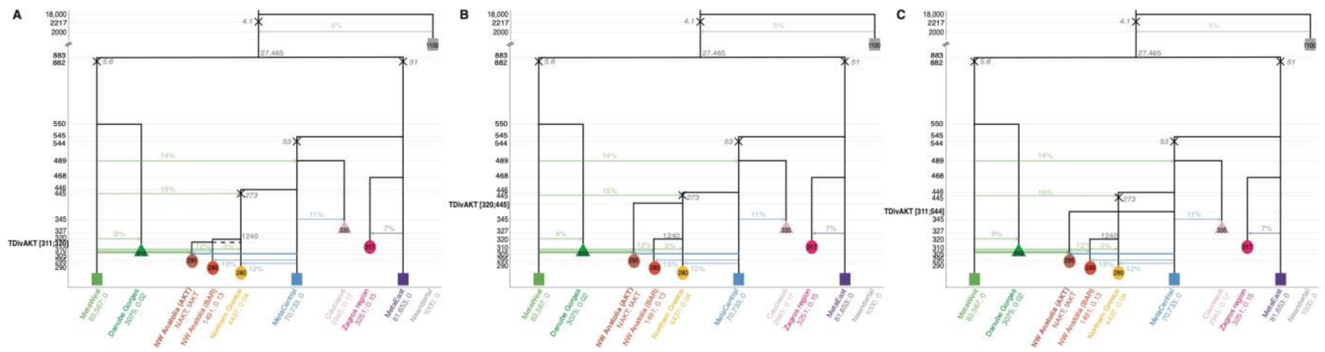


**Figure S34 - Schematic description of models tested for the *AKT* panel**

(**A**) **As a sister population of Barcın or of the Greek Neolithics.** Under this model, the AKT-related population has diverged TDivAKT generations ago from the Neolithic population from NW Anatolia or Greece, necessarily after their split 320 generations ago.

(**B**) **From the Aegeans.** Under this model, the AKT-related population has diverged TDivAKT generations ago from the ancestral Aegean population, i.e. before the split of the NW Anatolia and Greece Neolithics 320 generations ago but after the divergence of the branch from the *Central* metapopulation 445 generations ago.

(**C**) **From the *Central* metapopulation.** Under this model, the AKT-related population has diverged TDivAKT generations ago from the *Central* metapopulation as did the ancestral Aegean branch.

The AKT-related population received some gene flow from the *Western* metapopulation 310 generations ago and from the *Central* metapopulation 10 generations before sampling. These models are derived from the best model inferred for the Aegean dataset: parameter values estimated in the previous scenario were fixed in this analysis and shown in light fonts while newly estimated parameters are shown in bold fonts with search ranges shown between brackets. Further information about the parameters can be found in Supp. Table 4 and the properties of the sampled populations are listed in Table S4.

*Best model parameters estimates and SFS fit*

The model best explaining the *AKT* panel (Fig. S35, Supp. Table 4) is when the Aktopraklık individual is drawn from a small population (2Ne = 687 [145-4933]) emerging recently as a sister population from the Northern Greece population ~9.1 kya [9.1-9.2], 6 generations after the split between the Barcın and

Greek population, suggesting the existence of an Aegean cluster to which he Aktopraklık, Barcın and Northern Greece populations belong as they have diverged from each other at about the same time. Despite this recent split within the Aegeans, the Aktopraklık-related population has 17% [11-18] of its genome drawn from the *Western* metapopulation, and another 13% [5-17] from surrounding Neolithic populations (modeled by the *Central* metapopulation), i.e. a proportion similar to that received by the two other Aegean populations which might suggest a single ancestral pulse of gene flow before their split. The quite important *Western* metapopulation contribution to the Aktopraklık genome is in line with the admixture analysis (Fig. 3B). Note that we observed a good fit between the observed and simulated SFS using the backbone approach (Fig. S35C).

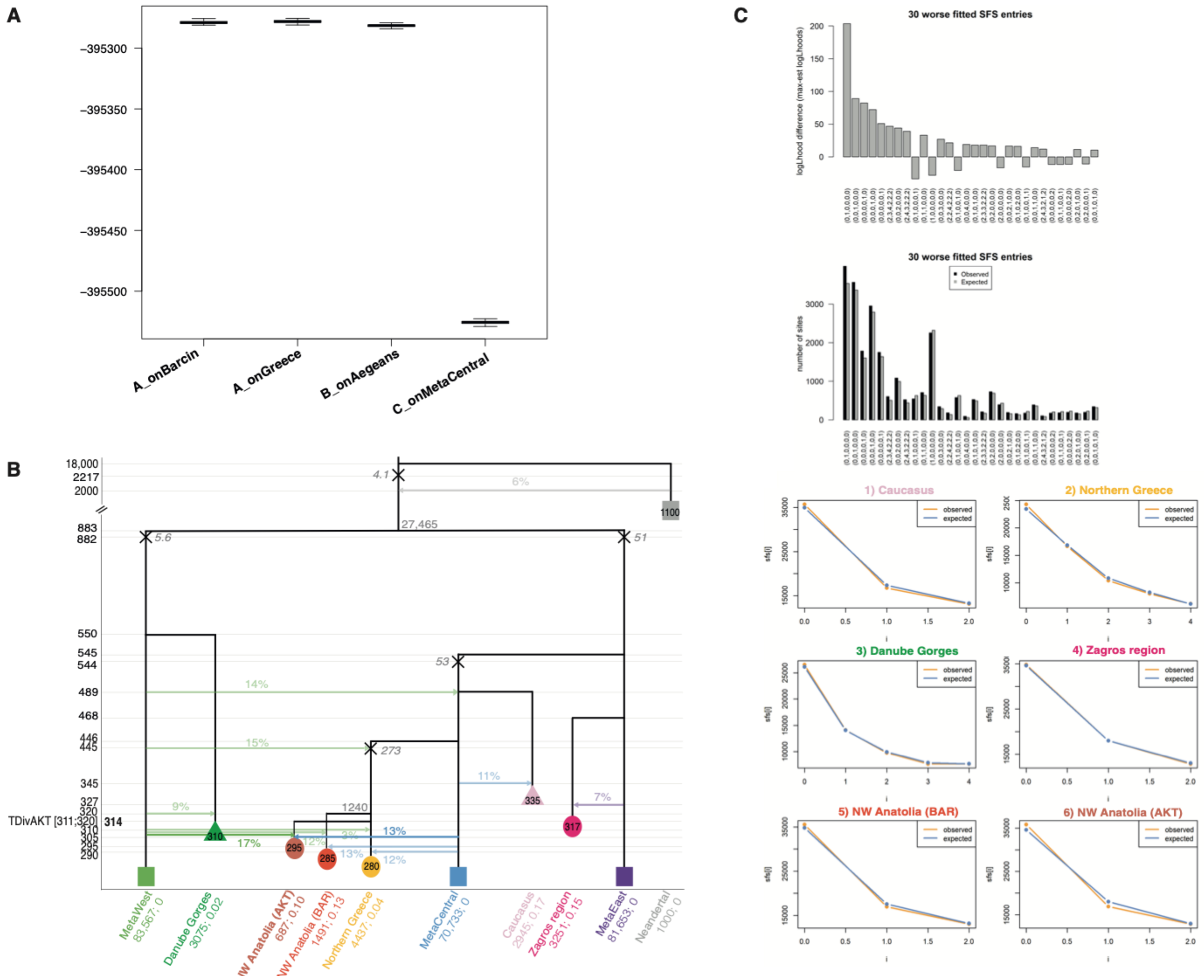


**Figure S35 - Exploration of models best explaining the observed multi-dimensional *AKT* SFS**
(**A**) **Distribution of the log10-likelihoods** obtained by simulation from the maximum likelihood point estimates for the four tested models. Higher likelihoods (top of the plot) reflect better simulations.

(**B**) **Schematic representation of the ML demographic model *A_onGR*.** Values of various parameters are reported with their 95% CI estimated from 100 parametric bootstrap estimations. Close to the bottleneck symbol (X), we report the size of a bottleneck, conveniently modeled as a single generation bottleneck (*instbot fastsimcoal2* option).
(**C**) **Fit between the observed and estimated SFS under the ML model** (multidimensional SFS on the upper panel; marginal SFS for each population on the bottom).

<u>Relationship between Aegean and Central Anatolian early farmers</u>

*Tested models*

The significant Mesolithic European-like contribution (that we model as the *Western* metapopulation) found in the ancestors of all European and NW Anatolian Neolithics as well as Caucasus HGs raises the question of the geographic location of this HG input, and whether it can also be observed in Central Anatolia too. We examined these questions by building on the best scenario found for the *Aegean* panel and including the Bon002 (Boncuklu) early Neolithic individual from the Konya Plain in Central Anatolia. In contrast to *AKT* panel, the lower number of genomic sites available in the Bon002 genome (due to its lower coverage) led us to reduce the dimensions of the SFS and only keep the Danube Gorges Mesolithic, Northern Greek and Iranian Neolithic samples in this panel (Table S4). Thus, we were only able to test two models: A) in which the Central Anatolian population branches from the Aegean ancestral population, or B) from the *Central* metapopulation (Fig. S36).

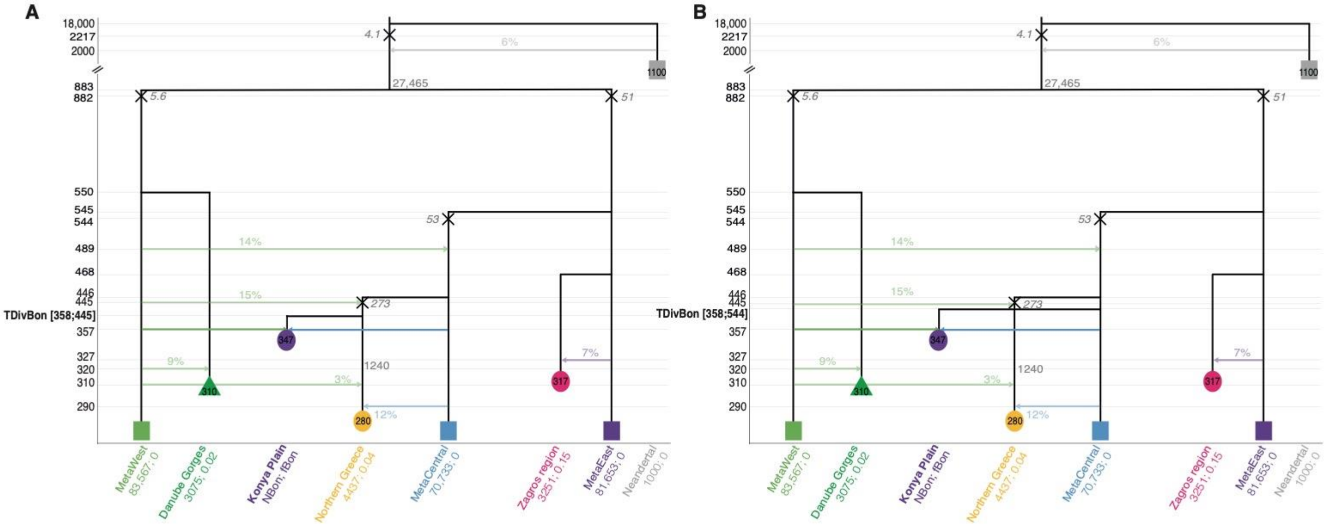


**Figure S36 - Schematic description of models tested for the *Bon* panel**
(**A**) **From the Aegeans.** Under this model, the Bon-related population has diverged TDivBon generations ago from the ancestral Aegean population, i.e. before the split of the NW Anatolia and Greece Neolithics 320 generations ago, but after the divergence of the *Central* metapopulation 445 generations ago.

(**B**) **From the *Central* metapopulation.** Under this model, the Bon-related population has diverged TDivBon generations ago from the *Central* metapopulation.
The Bon-related population received gene flow from the *Western* and *Central* metapopulations 10 generations before sampling. These models are derived from the best model inferred for the *Aegean* panel: parameter values estimated in the previous scenario were fixed and shown in light fonts, while newly estimated parameters are shown in bold fonts with search ranges shown between brackets. Further information about the parameters can be found in Supp. Table 4 and the properties of the sampled populations are listed in Table S4.

*Best model parameters estimates and SFS fit*

Our analysis reveals that the Boncuklu individual has diverged from the Aegean ancestral population some 362 [361-378] generations ago (~10.5 kya) during the cold Younger Dryas period, and after the admixture with *Western* metapopulation shared by all the Aegeans (Fig. S37, Supp. Table 4). Therefore it appears that Bon002 from Central Anatolia shares an ancestry with the NW Anatolian and European Neolithics, and that the Iranian Neolithic population seems the only sampled Neolithic population not to have received any European-like HG admixture. An additional *Western* metapopulation genetic input, forced to be received 10 generations before sampling, contributed to 10% [3-15] of the Boncuklu's genome. Furthermore, this population seems to be more isolated than other Aegean Neolithic populations since only 2% [1-11] of the Boncuklu gene pool comes from *Central* metapopulation. However, the results obtained for this dataset should be considered with caution due to the small number of polymorphic sites available for these analyses.

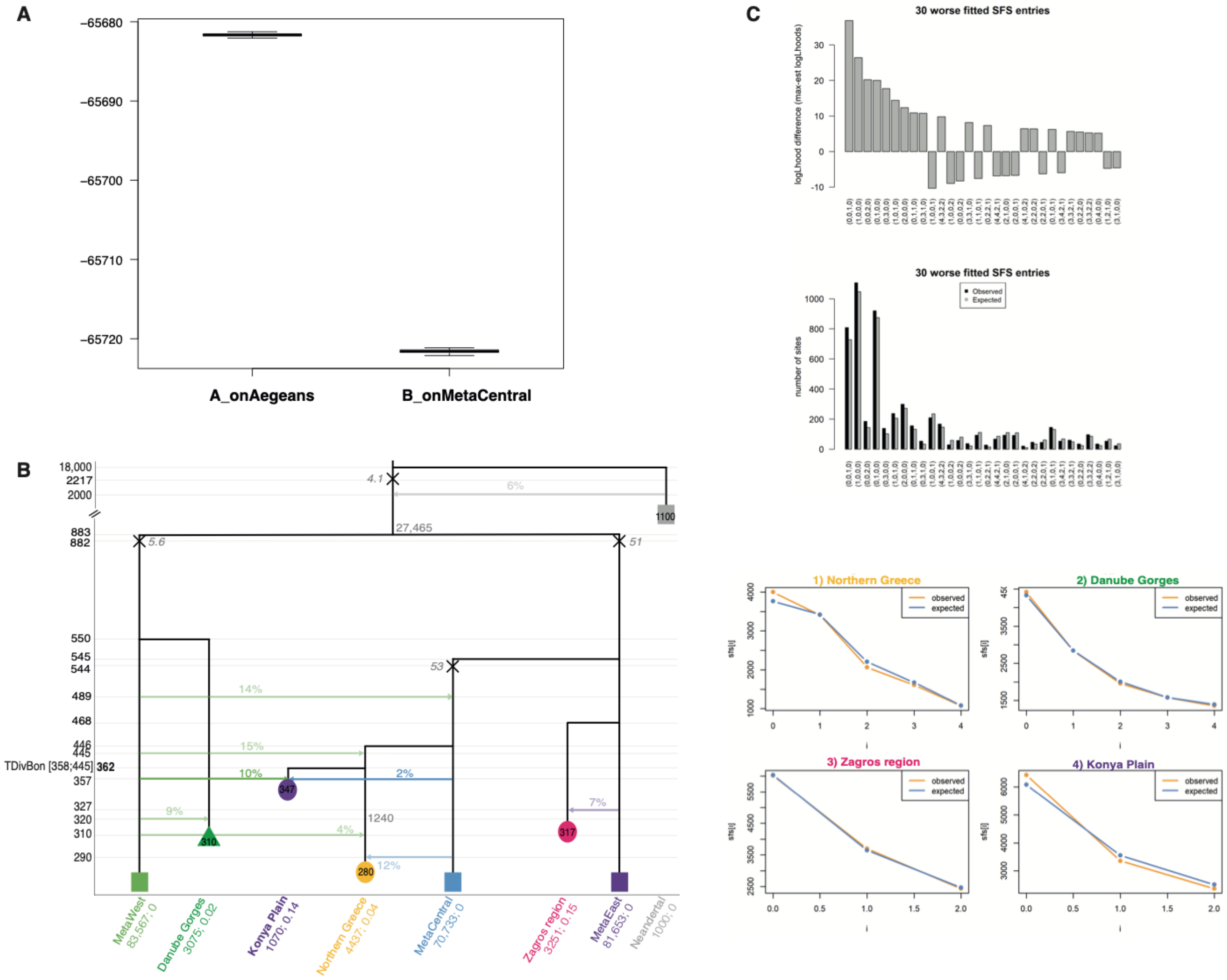


**Figure S37 - Exploration of models best explaining the observed multi-dimensional *Bon* SFS**

(**A**) **Distribution of the log10-likelihoods** obtained by simulation from the maximum likelihood point estimates for the two tested models described in Fig. S36. Higher likelihoods (top of the plot) reflect better fit.

(**B**) **Schematic representation of the ML demographic model *A_onAegeans*.** Values of various parameters are reported with their 95% CI estimated from 100 parametric bootstrap estimations. Close to the bottleneck symbol (X), we report the size of a bottleneck, conveniently modeled as a single generation bottleneck (*instbot fastsimcoal2* option).

(**C**) **Fit between the observed and estimated SFS under the ML model** (multidimensional SFS on the upper panel; marginal SFS for each population on the bottom).

Spread of Neolithic along the Danube

*Tested models*

In order to study the spread of Neolithic farmer populations from the Aegean-Marmaran basin to Central Europe along the Danube corridor, we considered samples from Central Serbia, Lower Austria, and Southern Germany in addition to those from Greece, Caucasus and Mesolithic individuals from the Danube Gorges (Table S4). In this analysis, we did not consider NE1 who is a geographic outlier nor the two individuals from Lepenski Vir, even though they are located in the Danube corridor, as individuals from this site show recent admixture with Iron Gates hunter-gatherers and display subsistence and burial practices more akin to the European Mesolithic (*142*).

We built two classes of models (Fig. S38): A) here we assume that the spread of farmer populations consisted in a series of stepping-stone (SS) migration events from Northern Greece to Central Serbia, then from Central Serbia to Lower Austria, and finally from Lower Austria to Southern Germany; B) here we explored a partial SS model, where Lower Austria could be directly settled from Northern Greece, thus bypassing Central Serbia. For both types of models, we allowed Neolithic populations to receive some input from local Mesolithic populations, modeled by admixture from the *Western* metapopulation occurring at the founding of the Neolithic population. We re-estimated here the parameters for the Neolithic population from Northern Greece too as this population is the source of the Danubian subsequent populations.

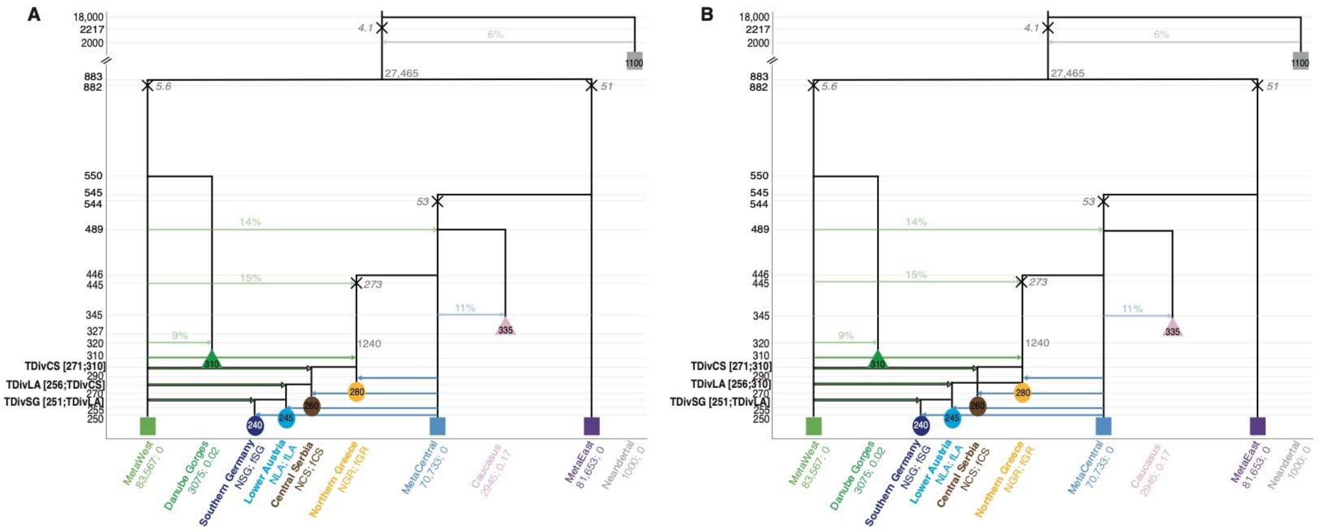


**Figure S38 - Schematic description of models tested for the *NeoEur* panel**

(**A**) **Pure stepping-stone model.** The Central Serbian Neolithic population derived from Northern Greece TDivCS generations ago, the Lower Austrian Neolithic from Central Serbia TDivLA generations ago and Southern German Neolithic population from Lower Austria TDivSG generations ago.

(**B**) **By-passing model.** Here the Central Serbian and Lower Austrian Neolithic populations are both settled from Northern Greece TDivCS and TDivLA generations ago, respectively, then the Southern German Neolithic population derived from Lower Austria TDivSG generations ago.

In the models with admixture, the Neolithic populations can receive some gene flow from the *Western* metapopulation at the time of their founding (TDivCS, LA or SG). They receive in all models some gene flow from the *Central* metapopulation 10 generations before sampling. These models are derived from the best model inferred for the *Aegean* panel: parameter values estimated in the previous scenario were fixed and shown in light fonts, while newly estimated parameters are shown in bold fonts with search ranges shown within brackets. Further information about the parameters can be found in Supp. Table 4 and the properties of the sampled populations are listed in Table S4.

*Best model parameters estimates and SFS fit*

We find that stepping-stone models are better supported than their corresponding bypassing model (Fig. S39A, Supp. Table 4), suggesting some stepwise expansion of the early farmers in Europe along the Danube between ~8 and 7.5 kya. Interestingly, admixture from the *Western* metapopulation always improves the fit, suggesting that the early Neolithic populations incorporated a few HG individuals (2-7%) at all stages of the dispersal along the Danubian corridor (Fig. S39B, Supp. Table 4). Finally, the early Neolithic populations were well connected to other farmer communities, as modeled by single pulses of gene flow coming from the *Central* metapopulation with intensity between 4 and 8%. Interestingly, this continuous gene flow from less-mixed neighbouring farmer populations could counterbalance the rate of *Western* metapopulation admixture entering the farmer gene pool during the expansion. This complex pattern of gene flow might explain the apparent lack of genetic structure among early farmers observed in the MDS plot (Fig. 2A) and the absence of increasing *Western* ancestry along the Danubian corridor in the admixture analysis (Fig. 2B).

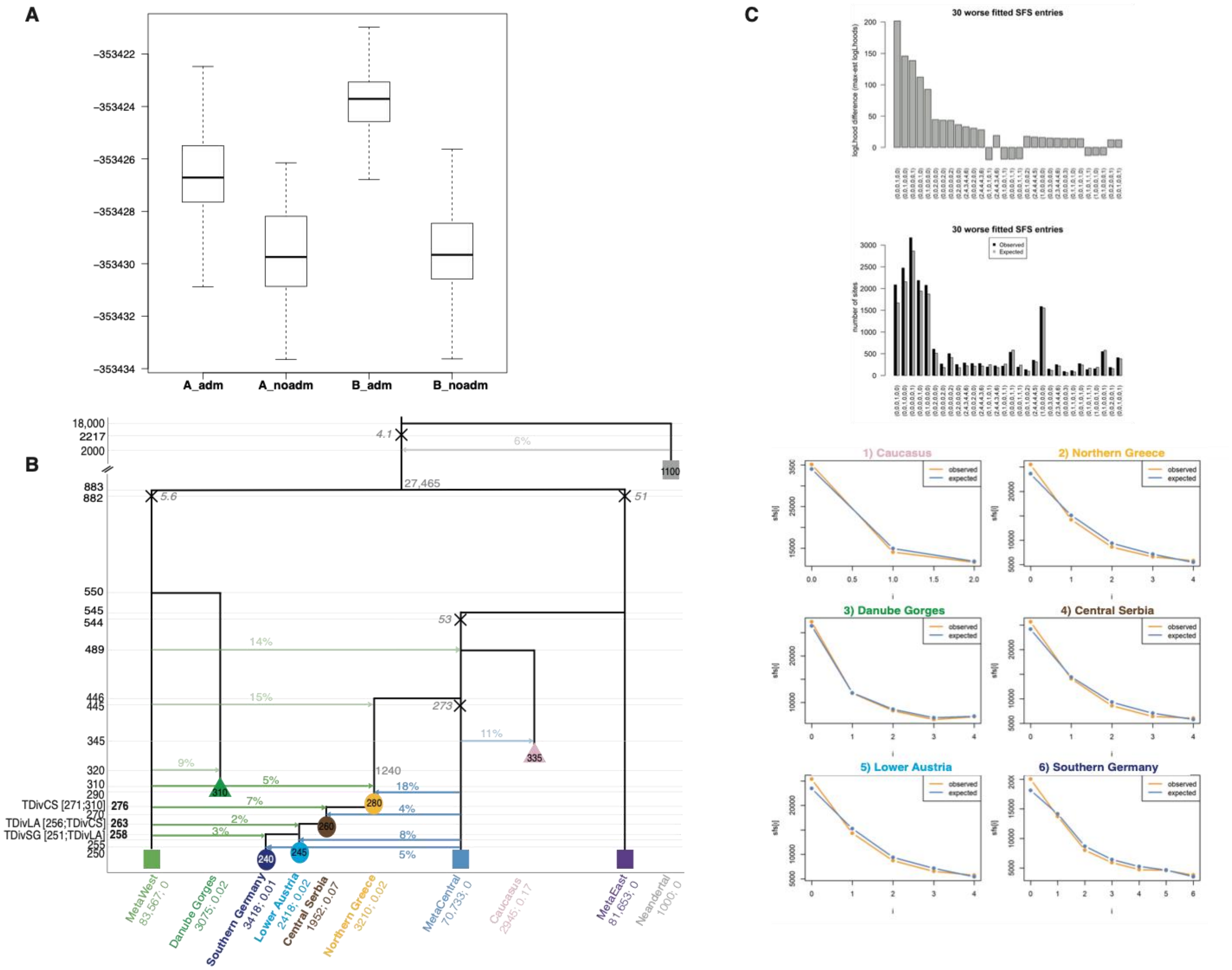


**Figure S39 - Exploration of models best explaining the observed multi-dimensional *NeoEur* SFS**

(**A**) **Distribution of the log10-likelihoods** obtained by simulation from the maximum likelihood point estimates for the four tested models described in Fig. S38. Higher likelihoods (top of the plot) reflect better simulations.

(**B**) **Schematic representation of the ML demographic model *A_adm*.** Values of various parameters are reported with their 95% CI estimated from 100 parametric bootstrap estimations. Close to the bottleneck symbol (X), we report the size of a bottleneck, conveniently modeled as a single generation bottleneck (*instbot fastsimcoal2* option) .

(**C**) **Fit between the observed and estimated SFS under the ML model** (total on the upper panel; marginal for each population on the bottom).

Relationship between Western and Central European HG populations

*Tested models*

In order to investigate the relationship between Western and Central European HG populations, we included in the panel *HG* two previously published good quality genomes: the Upper Palaeolithic individual from Bichon and the Mesolithic individual from Loschbour, the two Mesolithic genomes from the Danube Gorges as well as the genomes from the Mesolithic Caucasus and from Neolithic Iran. We considered two models: a shared ancestry for Lochbour and Bichon (Fig. S40A) with a Western European ancestral population that splitted from the *Western* metapopulation before diverging into the Bichon and Loschbour-related populations, or two independent splits from the *Western* metapopulation for the populations ancestral to Bichon and Loschbour respectively (Fig. S40B). In both models, we estimated the demographic parameters related to the three European populations (Danube Gorges, Bichon and Loschbour) and used for the other parameters the values obtained for the best *HGNeo* run.

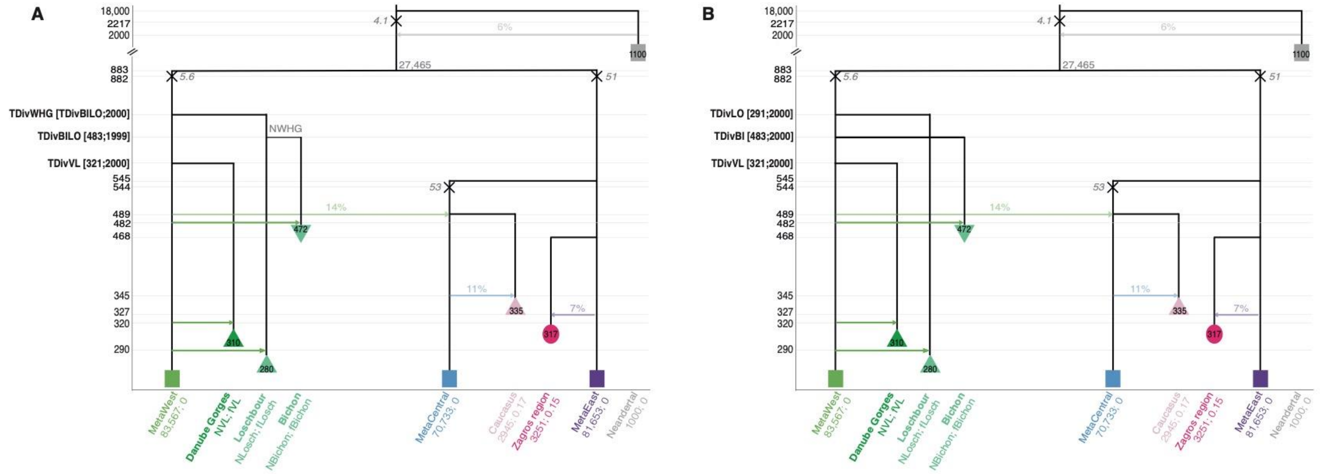


**Figure S40 - Schematic description of models tested for the *HG* panel.**
(**A**) **An ancestral Western European HG (WHG) structure.** Under this model, the WHG ancestral population has diverged TDivWHG generations ago from the *Western* metapopulation with a resize event; then, the ancestral population split into populations related to Loschbour and Bichon respectively, TDivBILO generations ago; then both sampled populations received some gene flow from the *Western* metapopulation 10 generations before their sampling time.
(**B**) **Independent split events within the WHGs.** Under this model, populations related to Bichon and Loschbour independently split from the *Western* metapopulation TDivBI and TDivLO generations ago respectively, before they received independent pulses of gene flow from the *Western* metapopulation ten generations before sampling time.
These models are derived from the best model inferred for *HGNeo*: parameter values estimated in the previous scenario were fixed and shown in light fonts, while newly estimated parameters are shown in bold fonts with search ranges shown within brackets. Note that all parameters of the HG population from the Danube Gorges (i.e. NVL, fVL, TDivVL, admMetaW.VL) are re-estimated here. Further information

about the parameters can be found in Supp. Table 4 and the properties of the sampled populations are listed in Table S4.

*Best model parameters estimates and SFS fit*

We find that the model where Bichon and Loschbour have a common ancestry is best supported (Fig. S40C, Supp. Table 4). In this scenario, the ancestral population of these two individuals would have diverged from the *Western* metapopulation during the LGM, 787 [579-851] generations ago (~22.8 kya [16.7-24.7]), 100 generations after its divergence from the *Eastern* metapopulation. Note that our modeling did not constrain them to have diverged from the *Western* branch as they could have diverged before the LGM from the ancestors of the two metapopulations. It implies that during the LGM, the HG population split into (at least) three components (Western European, Central European and Near-Eastern/Caucasus HGs). Interestingly, the divergence of these Western European HGs from the *Western* metapopulation has occurred after the very severe bottleneck linked to the divergence of the metapopulations (about two diploid individuals for one generation or 25 individuals for 10 generations). This bottleneck seems to have affected all European HGs and it explains their reduced diversity as compared to most Neolithics, e.g HGs clustering all together on MDS plots, low heterozygosity, numerous short and sometimes long ROHs (Fig. 2, Fig. S19). Then, the Loschbour and Bichon populations seem to have split during the LGM some 748 [501-748] generations ago (~21.7 kya [14.5-21.7]) and do not seem to have remained isolated after the LGM since they received about 8-14% [2-13] of admixture from the *Western* metapopulation. Contrastingly, the HG population from the Danube Gorges split more recently (~14.7 kya [11.3-22.9]) from the *Western* metapopulation, suggesting some closer affinities with the HG local populations that admixed with the Neolithic populations of our study.

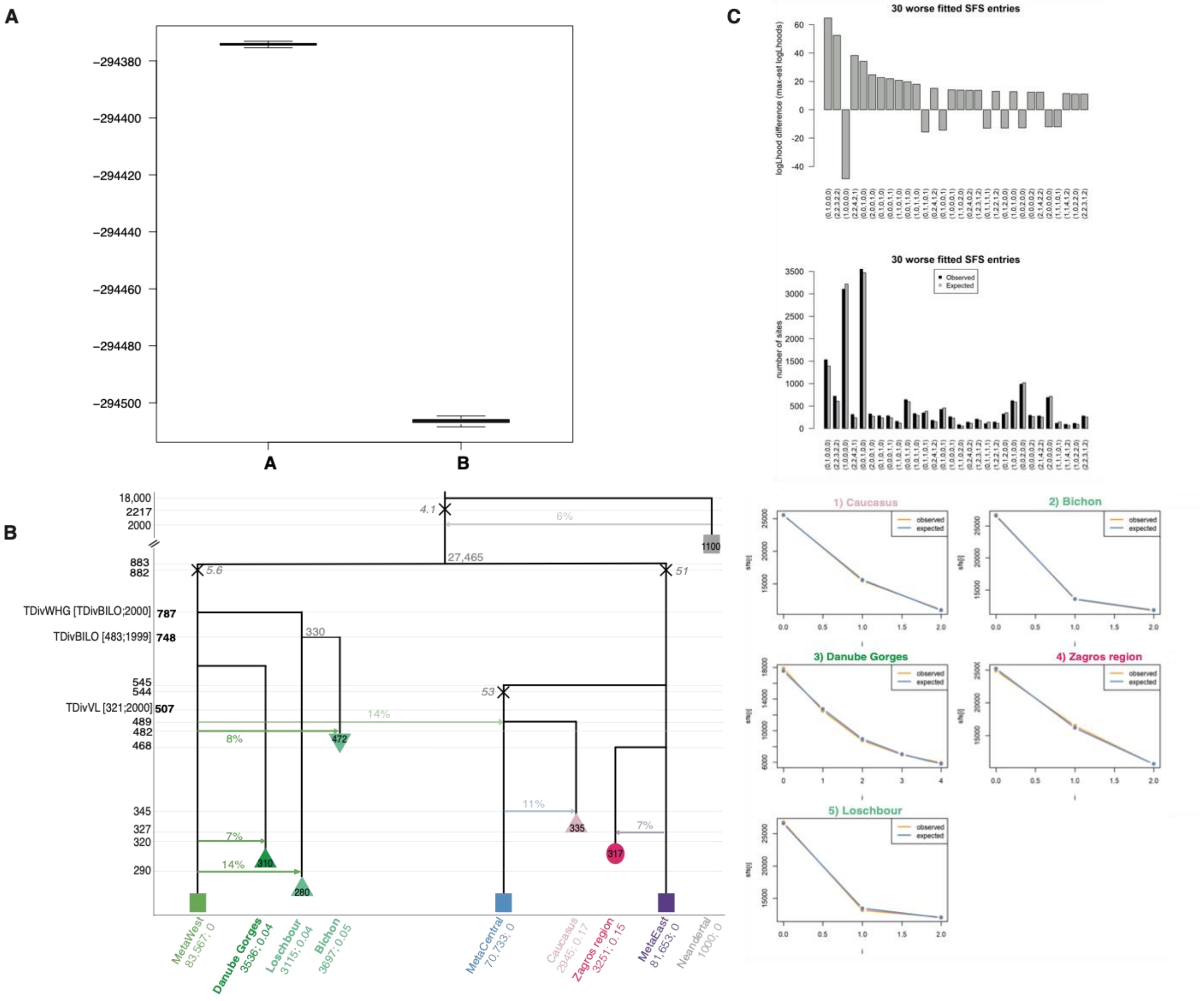

**Figure S41 - Exploration of models best explaining the observed multi-dimensional *HG* SFS**

(**A**) **Distribution of the log10-likelihoods** obtained by simulation from the maximum likelihood point estimates for the two tested models described in Fig. S40. Higher likelihoods (top of the plot) reflect better simulations.

(**B**) **Schematic representation of the ML demographic model *A*.** Values of various parameters are reported with their 95% CI estimated from 100 parametric bootstrap estimations. Close to the bottleneck symbol (X), we report the size of a bottleneck, conveniently modeled as a single generation bottleneck (instbot *fastsimcoal2* option) i.

(**C**) **Fit between the observed and estimated SFS under the ML model** (multidimensional SFS on the upper panel; marginal SFS for each population on the bottom).

**Final Model**

We assembled as a single *fastsimcoal2* input file (.par) a complex scenario including the maximum-likelihood parameter values of the best models for the 6 panels, corresponding to what is described in Fig. 3A. Thus, this model includes the 12 sampled populations (Table S4), the three metapopulations and a Neandertal population with parameters estimated in the different scenarios (in particular, Vlasac's parameters have been taken from the *HG* best scenario and the Greeks' from the *Aegean* best scenario). For later use (see section on *f*-statistics below), we added an African population for which we chose a haploid population size of 20,000, a null $F_{IS}$, and a split time from the European/SWAsian group some 97.1 kya (i.e. 3350 generations ago). Eventually, we sampled 11 ancestral populations at key moments of the demography we reconstructed: on the ancestral branch before the split between *Western* and *Eastern* metapopulations 25.6 kya (i.e. 883 generations ago), on the *Western* and *Central* metapopulations branches just before the admixture occurring 14.2 kya (i.e. 490 generations ago), on the *Central* metapopulation branch just after this event (489 generations ago); on the *Eastern* metapopulation branch at the time of split of the Konya Plain population; on the Aegean ancestral branch just after its founding and its admixture with the *Western* metapopulation (445 generations ago = 12.9 kya), then every 25 generations until the split into Aegean distinct populations (related to Northern Greece, Aktopraklık and Barcın; 320 generations ago = 9.3 kya).

***Validating the proposed demographic model***

**Validation of parameter estimates from parametric bootstraps**

In addition to the clear good fit found between the expected and observed SFS for every model, the parametric bootstrap approach used to compute parameters confidence intervals actually brings a different validation of our estimation procedure. The fact that we can infer confidence intervals around our estimated parameters shows that we can correctly recover the simulated parameters with our approach. Indeed, for every dataset, all but four point estimates (except admMetaW.LO in *HG*; admMetaC.BA in *Aegean*; NLA, f_CS in *NeoEur)* were found to lie within the 95% confidence intervals, showing that if the true model was that we estimated, we should be able to recover it with our maximum-likelihood estimation procedure.

**Predictive Simulations**

We validated the scenario from Fig. 3A using several complementary approaches, as outlined below. First, we tested if we were able to reproduce the actual genetic diversity observed within ancient samples under the scenario shown in Fig. 3A. We simulated some genetic data as 10 million independent segments of 100bp for 10 different replicates (we obtained between 1,829,466 and 1,834,676 polymorphic sites among the replicates) and a mutation rate of $7.5e^{-9}$ (chosen within the range of rates of the 6 panels).

We then computed the nucleotide divergence $\pi_{XY}$ between all diploid samples. We found a significant correlation between simulated and observed distances computed over the neutral portion of the genome (Mantel test: r = 0.8, p-value = 0.001, with similar values for the 10 replicates). We then performed a MDS analysis of the $\pi_{XY}$ genetic distances computed among the Europeans and SW Asians simulated samples (Fig. 3B). We see that the simulated MDS is very similar to the observed one (Fig. 2A), with a cluster of HGs, a cluster of Neolithics and the group of the Iranian and Caucasus samples, showing that our model can reproduce very well the observed genetic structure of the ancient samples. By simulating data for ancestral populations, we gain some interesting insight on the admixture process between the *Central* and the *Western* metapopulations.

Admixture analyses performed on simulated data are also really able to replicate observed patterns (compare Fig. S42A and B). It is possible to simulate data for ancestral populations too (Fig. S42C), an approach that offers the opportunity to better understand the biological processes causing the observed pattern. Indeed, we can see the increasing proportion of the green ancestry from *MetaCentral (490g)* to *MetaCentral (489g),* i.e. just before and one generation after its admixture with the *Western* metapopulation; this signal of admixture is also visible in the Aegean ancestral population (*445g*) that is modeled as receiving a second gene flow in our model. Furthermore, we can see that the red ancestry found in the Neolithic samples seems to appear progressively through time with its proportion increasing in the Aegean ancestral populations sampled every 25 generations. The Aegean population is changing only because of drift with a small population size estimated for this population ancestral to European and Anatolian early farmers (Ne = 620, 95% CI 72-2150). Thus, drift makes the Neolithic genomes look like they were from a completely independent gene pool and erases the signal of admixture from the *Western* metapopulation, explaining why the hybrid nature of early farmers had been previously unnoticed.

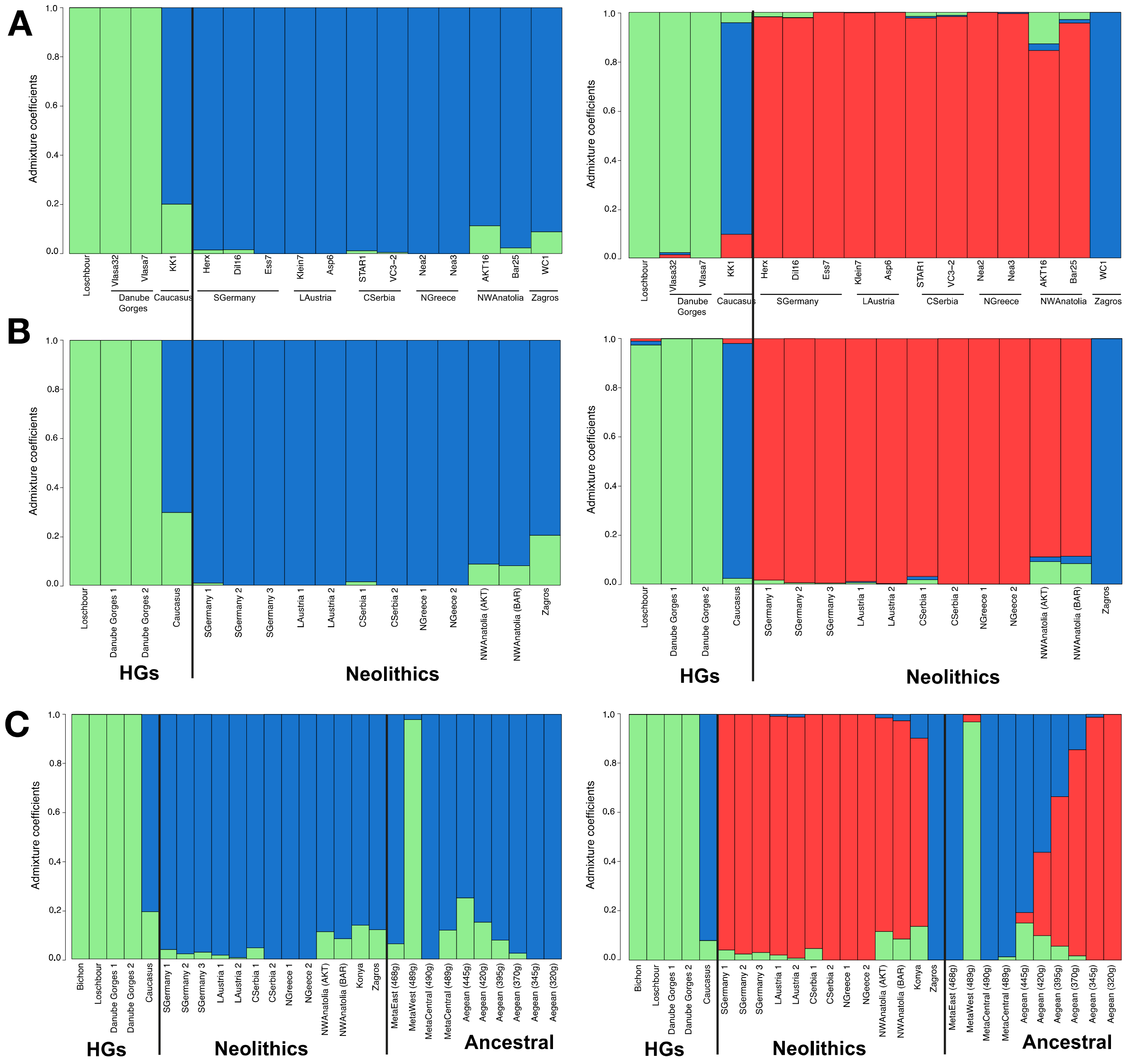

**Figure S42 - Admixture plot for K = 2 (left panel) and K = 3 (right panel)** carried on (**A**) observed non-missing data for 16 ancient genomes included into *fastsimcoal2* demographic analyses and best scenario (909,688 sites without missing data; Bichon and Bon002 were not included because of too low quality); (**B**) simulated data for the same individuals as in A); (**C**) simulated data for the 12 sampled populations represented by 18 individuals used in the demogenomic analyses, and the ancestral populations at different sampling times (1 individual each; see Fig. 3A and Final Model section for the details).

**f-statistics**

Confirming population assignment

As shown in Fig. S43, only very few comparisons violated our assignment. First, *AKT16* from the NW Anatolia region showed some WHG ancestry (represented by *Loschbour* and *Bichon*) not present in *Bar25*. This led us to model these two NW Anatolian individuals as independent populations in the *fastsimcoal2* demogenomic analysis. Second, more Central European HG ancestry (represented by the genomes from Vlasac VLASA7 and 32) was evidenced for *Asp6* than *Klein7* from Lower Austria. However, variation in HG ancestry is expected in early Neolithic populations due to the ongoing process of admixture. Since these samples did not show variation in their affinities to other Neolithic samples, modelling them as a single population seems justified.

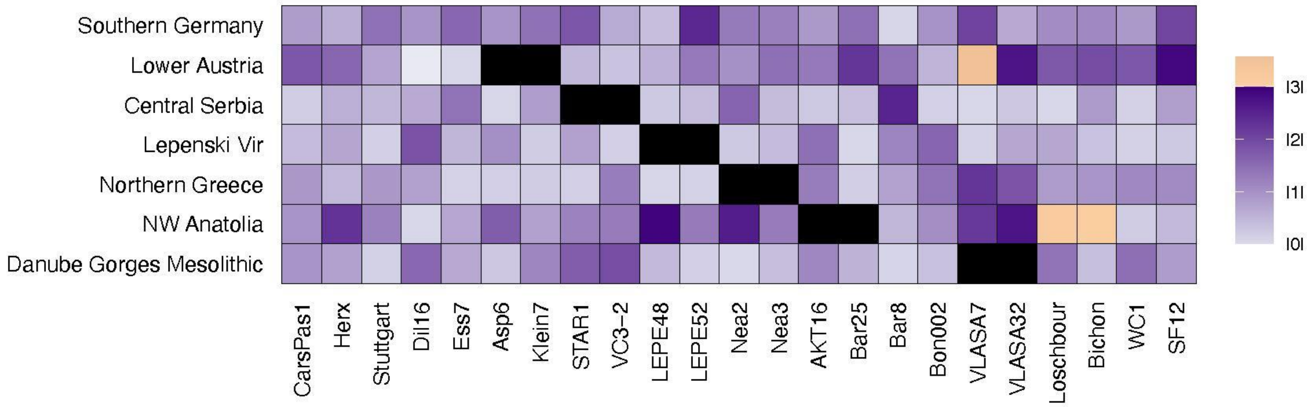


**Figure S43 - Heatmap of absolute Z-Scores of *f*-statistics of the form *D(Individual 1 from the population tested, Individual 2 from the population tested; Other samples, Outgroup)* for each population with two or more individuals and using Mbuti as the outgroup.** Salmon shades indicate significant absolute Z-scores above 3.0, purple shades indicate Z-scores below this threshold (black indicates comparisons of samples with their own population, for which *f*-statistics could not be calculated). For comparisons with the population of Germany that contained more than two samples, the highest Z-score of all possible two-way comparisons is shown.

Global model-fit using admixture graphs

To validate the model inferred by *fastsimcoal2*, we aimed at comparing *f*-statistics predicted under the *fastsimcoal2* model against those calculated from the data. Since the full graph could not be fit (see methods above), we created a simplified graph matching Fig. 3A model in structure, but containing fewer admixture edges. For this graph, no major differences between the *f*-statistics predicted under the model and those calculated from the simulated data were detected: For the best among 100 runs on this simplified graph (with the smallest final score), only one *f*-statistics showed a barely significant Z-score: *f4(NGreece, SGermany; Zagros, Caucasus), predicted 0.000265, observed 0.001644, Z-score 3.131*. But we note that the admixture graph analysis did not result in accurate estimates of all admixture proportions, as those were inferred rather variably among the 20 best runs (Fig. S44).

We next tested for discrepancies between the *f*-statistics predicted under this simplified graph and those calculated on the 12 sampled populations used in model fitting with *fastsimcoal2* (Table S4). For this purpose, we refitted the admixture graph 100 times with *qpGraph*. We first conducted this analysis on the set of neutral SNPs used to infer the SFS for model fitting (Fig. S45). On this data, no *f*-statistics was violated in the best graph, indicating that this graph is compatible with all *f*-statistics calculated on the real data. Since this graph was constructed to match the model inferred by *fastsimcoal2*, the *fastsimcoal2* model therefore is also fully compatible with these *f*-statistics.

To study the impact of using ascertained markers, we next repeated this analysis on the calls for the 1240K SNPs obtained as described in the Material & Methods section (Fig. S46). While the graph still generally holds for this set of markers, the best run resulted in twelve *f*-statistics that were significant, albeit mostly barely so:

1. *f4(CSerbia, NGreece; SGermany, NWA_AKT); pred. 0.002634, obs. -0.002462 Z-score -3.279*
2. *f4(NGreece, SGermany; NGreece, SGermany); pred. 0.260349, obs. 0.253955 Z-score -4.045*
3. *f4(NGreece, KonyaPlain; SGermany, NWA_AKT); pred. 0.002508, obs. 0.009600 Z-score 3.568*
4. *f4(NGreece, NWA_BAR; SGermany, Vlasac); pred. 0.000562, obs. 0.006496 Z-score 3.425*
5. *f4(NGreece, NWA_AKT; SGermany, KonyaPlain); pred. 0.003642, obs. 0.010112 Z-score 3.307*
6. *f4(NGreece, Vlasac; SGermany, NWA_BAR); pred. -0.005740, obs. 0.000405 Z-score 3.522*
7. *f4(NGreece, Vlasac; SGermany, NWA_AKT); pred. 0.004104, obs. 0.010444 Z-score 3.749*

8. *f4(NGreece, Loschb; SGermany, NWA_AKT); pred. 0.003970, obs. 0.010613 Z-score 3.139*
9. *f4(NGreece, Caucasus; SGermany, NWA_AKT); pred. 0.001200, obs. 0.007007 Z-score 3.004*
10. *f4(LAustria, KonyaPlain; NWA_AKT, Bichon); pred. -0.001021, obs. -0.008986, Z-score -3.237*
11. *f4(LAustria, NWA_BAR; Vlasac, Bichon); pred. 0.000162, obs. -0.006377, Z-score -3.641*
12. *f4(SGermany, NWA_BAR; Vlasac, Bichon); pred. 0.000230, obs. -0.006031 Z-score -3.796.*

This indicates that the marker choice is important and suggests that ascertained SNPs may hint at affinities not found when using neutral markers. However, we note that the graph used is likely too complex for the information contained in *f*-statistics and thus many branches and particularly admixture proportions were not reliably fitted with both sets of markers (large 90% quantiles across the 20 best runs, Fig. S45, Fig. S46). An important exception was the admixture event resulting in the source population of Anatolian and European farmers, which was inferred to have received 25% and 29% from the *Western* metapopulation on the neutral and 1024K set of markers, respectively, in agreement with a substantial contribution as inferred with *fastsimcoal2 (*8-26% 95% CI, Fig. S33).

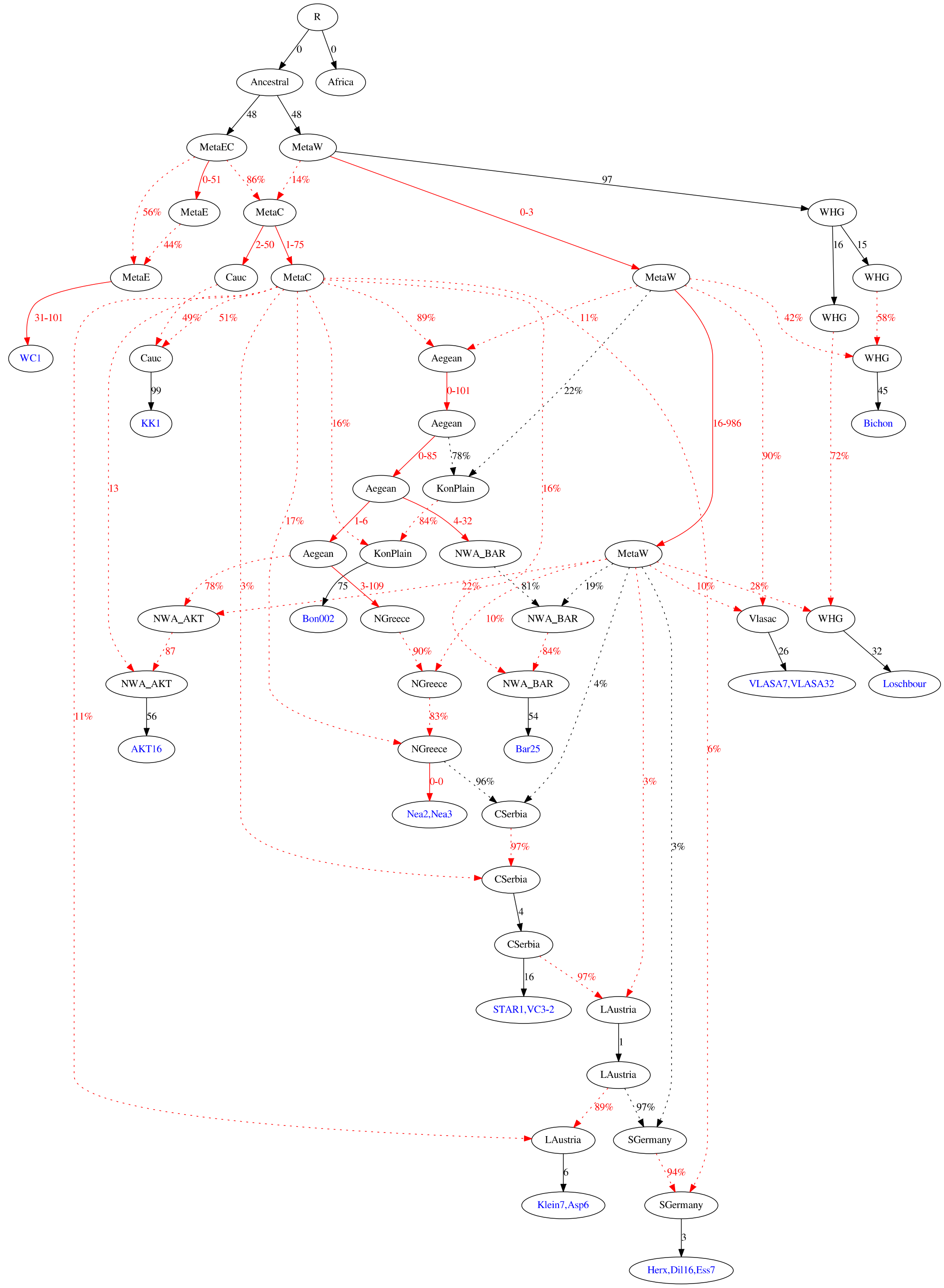


**Figure S44 - Admixture graph fitted to data simulated under the *fastsimcoal2* model.** Shown are the median branch lengths and mixture proportions among the best 20 graphs (lowest final score). Branches and mixture proportions are shown in red if the 90% quantile among the best 20 graphs was larger than given by half and twice the median (factor two) of the branch length or odds ratios, respectively. For outlier branches, the full range of estimates among the best 20 graphs are shown.

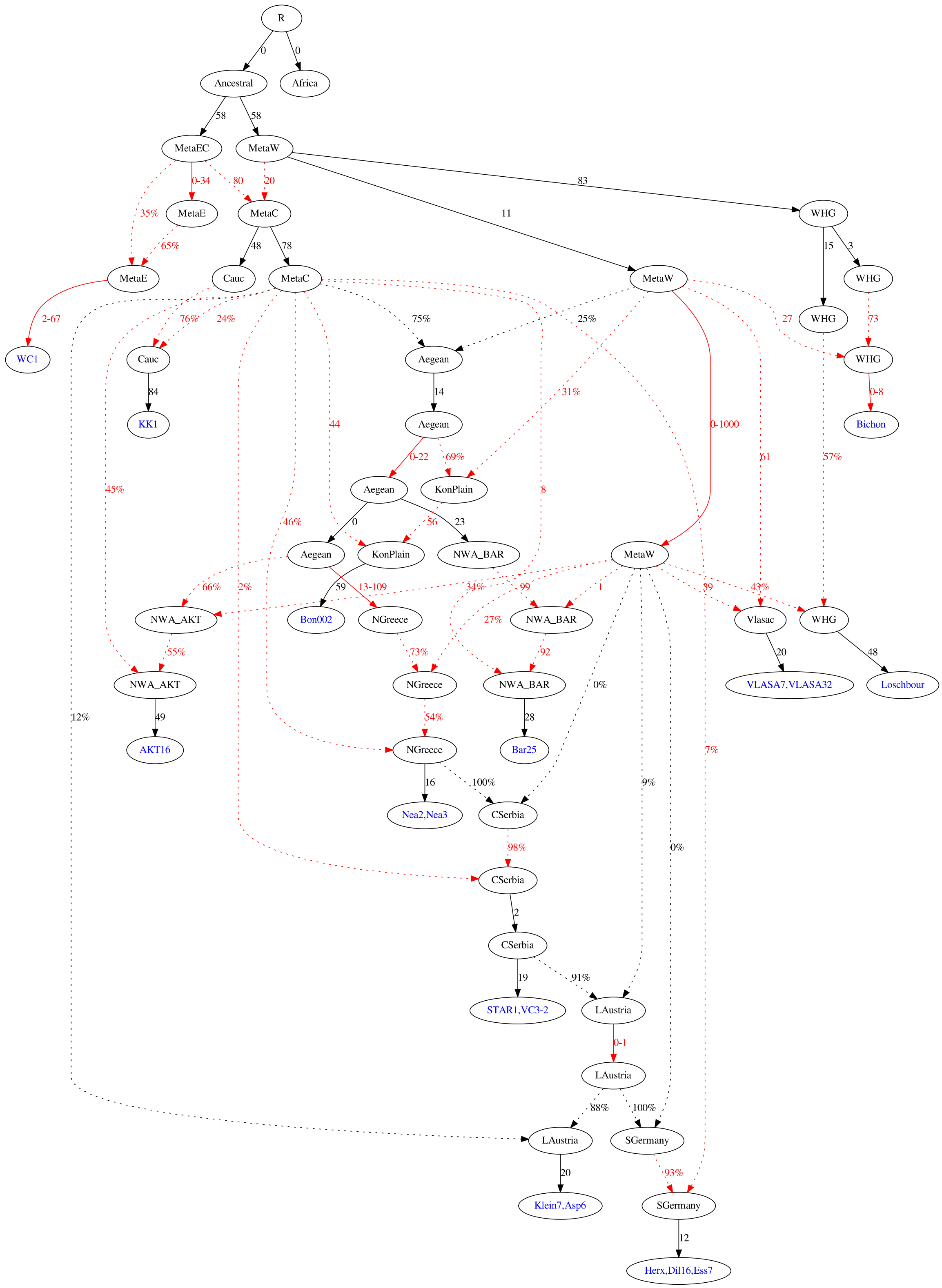


**Figure S45 - Admixture graph fitted to the *NeoEur* panel.** Color and labels as in Fig. S44.

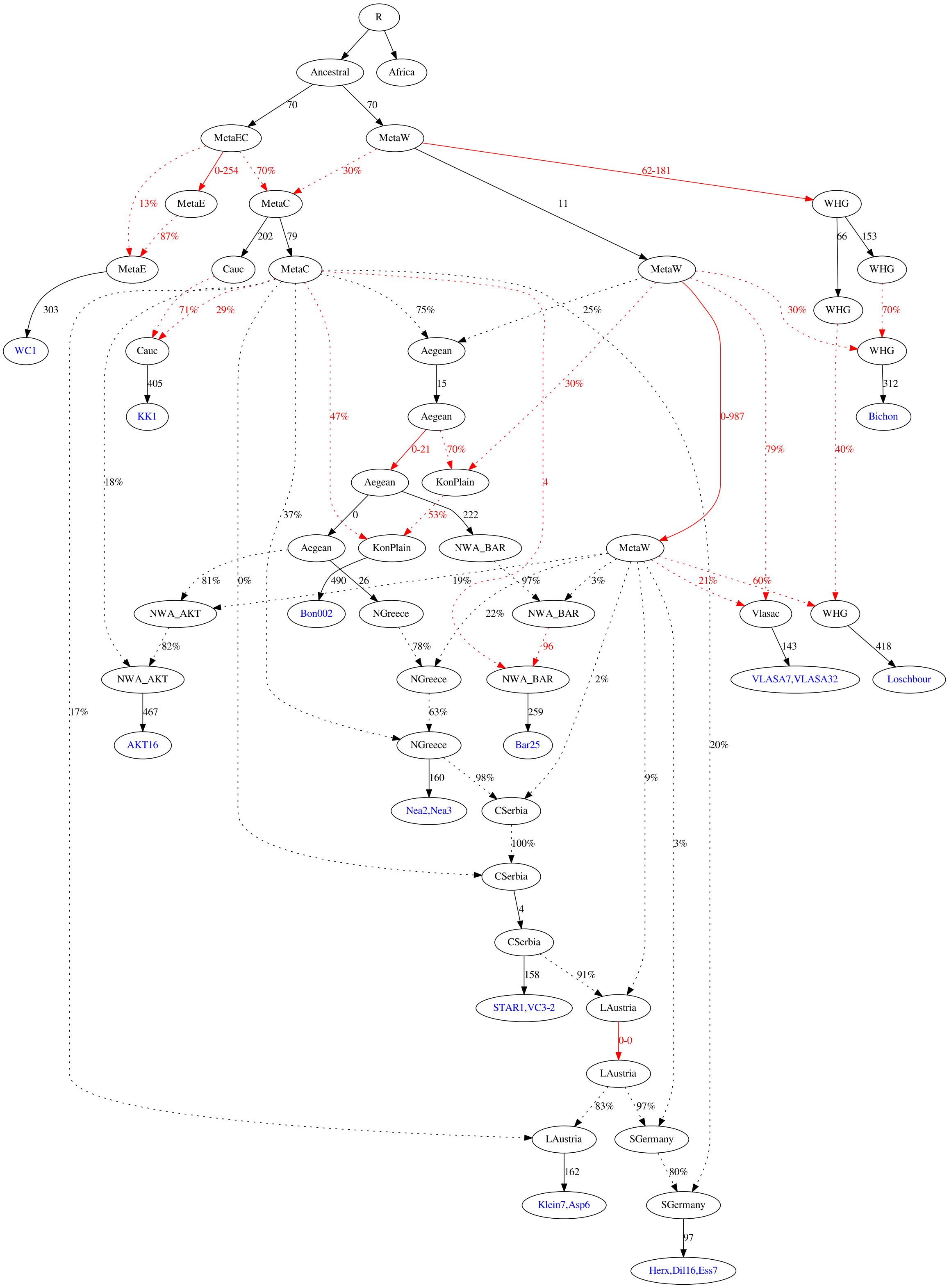


**Figure S46 - Admixture graph fitted to the 1240K dataset.** Color and labels as in Fig. S44**.**

Early separation of Konya Plain from Northern Greece

The *fastsimcoal2* analysis revealed an early split between the Konya Plain in Central Anatolia and the population leading to the Neolithic samples in the Aegean (Fig. S37). In the same analysis, admixture from the *Central* metapopulation was estimated to be substantially lower (2%) into Konya Plain than into Northern Greece (9%). In contrast, the estimates for admixture from the *Western* metapopulation into Konya Plain was estimated to be substantially larger (8%) than in Northern Greece (3%). Here we used two sets of *f*-statistics to support these findings.

*Excess Central/Western ancestry in the Konya Plain*

To confirm the excess HG ancestry in the Konya Plain, we used *f*-statistics of the form *D(NGreece/NWAnatolia, Konya Plain; HG_West, Outgroup)*. Here, *HG_West* is represented by the Vlasac individuals (*VLASA7* and *VLASA32*) from this study and *Bichon* and *Loschbour*. The Konya Plain is represented by 1) the four *Anatolia_Boncuklu* samples available in 1240k (*10*) consisting of the *Bon002* sample used in the *fastsimcoal2* analysis and three additional low-depth samples, and 2) the additional sample *Pınarbaşı*, an Epipalaeolithic HG from the Konya Plain dating to 15,592-15,023 cal BP (*12*). As shown in Fig. S47, we observed a significant excess of shared drift between *HG_West* and *Konya Plain* compared to *NGreece/NWAnatolia*. This signal was particularly strong for *Pınarbaşı* (all comparisons involving > 800,000 SNPs, Supp. Table 5). For *NWAnatolia_Boncuklu*, the signal was only present when compared to the late Neolithic Greek samples (*Pal7* and *Klei10*).

A notable exception from the general pattern is *AKT16* that shows excess shared drift with *HG_West* compared to *Konya Plain* (Fig. S47), in line with elevated *Western* metapopulation admixture estimated by *fastsimcoal2* (Fig. 3A) and also evidenced by the admixture analysis (Fig. 2B).

We conducted similar comparisons using *Neolithic Iranian* and *HG_East* samples from 1240K of the form *D(NGreece/NWAnatolia, Konya Plain; Neolithic Iranian/HG_East, Outgroup).* Here, *Neolithic Iranian* are represented by *WC1*, *Iran_TepeAbdulHosein_N* and *Iran_GanjDareh_N (243)* and *HG_East* is represented by *Georgia_Kotias* and *Georgia_Satsurblia (14)*. None of these comparisons revealed a significant difference between the samples from Northern Greece and *NW Anatolia* and those from the *Konya Plain* in their level of shared drift with neither *Neolithic Iranian* nor *HG_East* (Supp. Table 5), in line with the *fastsimcoal2* model that places these populations as sister groups.

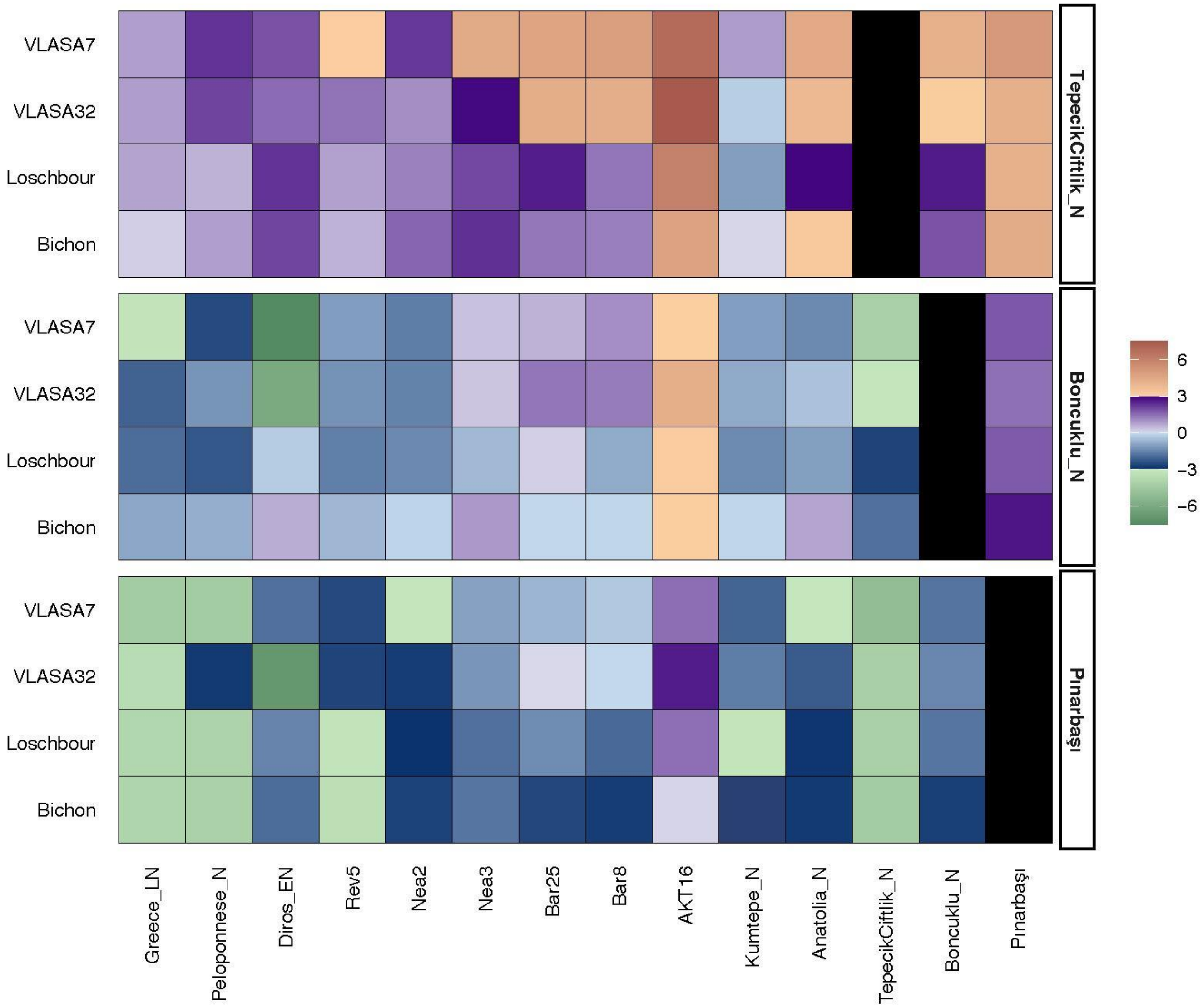


**Figure S47 - Heatmap of Z-Scores of f-statistics of the form *D(NGreece/NWAnatolia, Test; HG_West, Outgroup)*** using Mbuti as the outgroup and where Test is (listed from the top): Tepecik-Çiftlik, NW Anatolia_Boncuklu and Pınarbaşı. Red shades indicate significant positive Z-scores above 3, purple shades are for Z-scores below 3 and above 0, blue shades for Z-scores below 0 and above -3 and green shades indicate significant negative Z-scores below -3.

*Scarcity of Eastern metapopulation ancestry in the Konya Plain*

To confirm the scarcity of *Eastern* metapopulation ancestry in the Konya Plain, we used individuals from Israel from the Chalcolithic period (*Israel_C*), the PPNB period (*Israel_PPNB*) and the Natufian culture (*Israel_Natufian;* all Israel individuals were from *(9)*) as Eastern proxies and we performed *f*-statistics of the form *D(NGreece/NW Anatolia, Konya Plain; Israel, Outgroup)*. Most of the comparisons involving *NWAnatolia_Boncuklu* showed a strongly significant excess of shared drift between Israel and *NGreece/NWAnatolia* compared to *NWAnatolia_Boncuklu* (Fig. S48), in line with the scarcity of *Eastern* metapopulation admixture in the *Konya Plain* inferred by *fastsimcoal2*. Interestingly, however, this signal was not replicated when representing the *Konya Plain* by *Pınarbaşı* (Fig. S48): in that case, none of the comparisons was significant. Similarly, no significant comparisons were found when using either *Neolithic Iranian* or *HG_East* instead of *Israel* (Supp. Table 5).

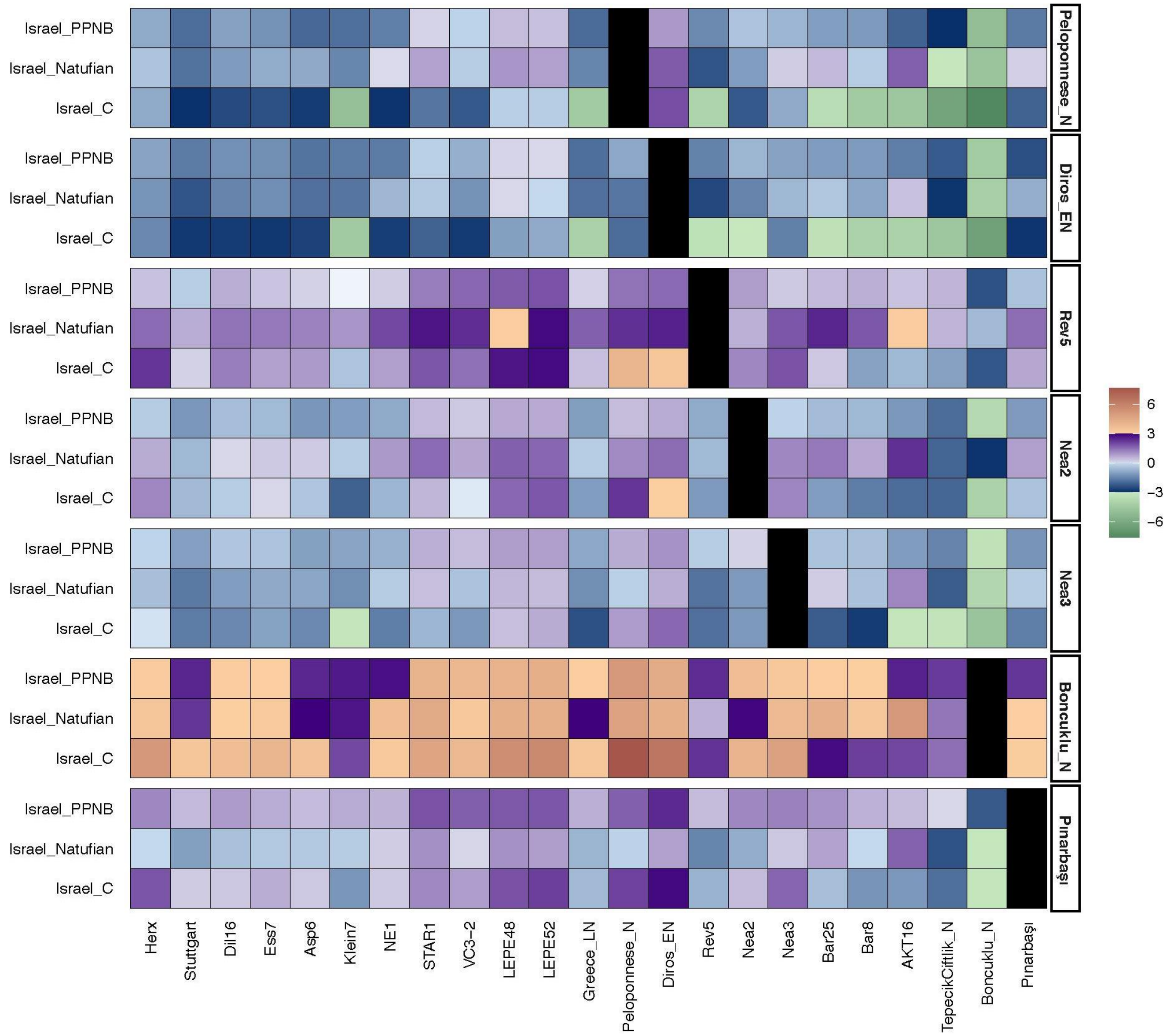


**Figure S48 - Heatmap of Z-Scores of f-statistics of the form *D(NGreece/NWAnatolia/early European Farmers, Test; Israel_PPNB/Natufian/C, Outgroup)*** using Mbuti as the Outgroup and Test as (listed from the top): Peloponnese_N, Diros_EN, Rev5, Nea2, Nea3, NW Boncuklu and Pınarbaşı. Red shades indicate significant positive Z-scores above 3, purple shades are for Z-scores below 3 and above 0, blue shades for Z-scores below 0 and above -3 and green shades indicate significant negative Z-scores below -3.

Structure among HG individuals

The model inferred by *fastsimcoal2* predicts a deep split between *Loschbour, Bichon* and the *Western* metapopulation. To validate these findings, we investigated the relationships among HG samples using *f*-statistics. First, we calculated *f*-statistics of the form *D(Loschbour, Bichon; HG_Central, Outgroup)*, representing *HG_Central* by the two Danube Gorges Mesolithic individuals *VLASA7* and *VLASA32*. In line with their deep split inferred by *fastsimcoal2*, none of these comparisons was significant.

We next investigated if the HG ancestry in *NWAnatolian* individuals can be attributed more to *HG_Central* than to *Bichon*, as predicted by the *fastsimcoal2* model. In line with these predictions, *f*-statistics of the form *D(Bichon, HG_Central; AKT16/Bar25, Outgroup)* resulted in significant, negative values (Z-score from -3.218 to -4.826).

Interestingly, the comparisons of the form *D(Loschbour, HG_Central; AKT16/Bar25, Outgroup)* were not significant and additional comparisons of the form *D(Loschbour, Bichon; AKT16/Bar25, Outgroup*) resulted in significant positive values. We interpret this as a result of sample age (*Bichon* is much older than *Loschbour*) and the admixture from *HG_Central* that was consequently estimated at a much older time for *Bichon* than *Loschbour*, allowing for shared drift in the *HG_Central* component in both *Loschbour* and *AKT16/Bar25*.

Noteworthy are further the many significant and positive *f*-statistics of the form *D(Bichon/Loschbour, HG_Central; HG_Adriatic, Outgroup)*, where *HG_Adriatic* is represented by Italian and Croatian Mesolithic and Epipalaeolithic HGs (see Supp. Table 5). This may point to an enhanced level of drift in the Danube Gorges Mesolithic samples (representing *HG_Central* here) and their relative uniqueness when compared to other HG populations in the region.

Differentiation within Greece and Konya Plain

**Peloponnese:** Based on differences between Neolithic samples from the Peloponnese to those from other regions, it was suggested that the Peloponnese might constitute a wave of Neolithization other than the Balkan route (*16*). Specifically, it was shown that Neolithic samples from the Peloponnese are shifted towards CHG-like ancestry and away from WHG-like ancestry, potentially due to contacts with populations like Kumtepe or Tepecik-Çiftlik that show similar patterns. We benefited from the additional samples generated in this study to shed additional light on the relationship between Neolithic samples from the Peloponnese and Northern Greece. We confirm that *Greece_Peloponnese* (four individuals from Diros Cave and one from Franchthi Cave labelled as *Greece_Peloponnese_N*, all Middle or Late Neolithic) are indeed more CHG-like than *Rev5*. However, this pattern is stronger when CHG is substituted with Iran_GanjDareh individuals (interestingly not WC1): *D(NGreece, Greece_Peloponnese; Iran_GanjDareh, Mbuti)* is significant and positive for all Greek Neolithic samples (Supp. Table 5). However, this signal is not obviously connected to Kumtepe or Tepecik-Çiftlik, as Greece Peloponnese samples also show more influence from the Levant (with Israel samples as proxies) than NW Anatolia, Greece (Late Greece) and Central European samples (Klein7), see Fig. S48. In addition, the same analyses conducted with the Peloponnese early Neolithic sample I5427 (labelled here *Diros_EN*) showed that higher amounts of Levantine ancestry was already present in the early Neolithic, at least compared to the Barcın samples (*Bar8* and *Bar25*). The sample *Diros_EN* also shows more shared drift with *Iran_GanjDareh* than *Nea2* and more influence from *Pınarbaşı* than the later Peloponnese group (*Greece_Peloponnese_N)*.

**Northern Greece and NW Anatolia:** In general, samples from Northern Greece and NW Anatolia show a relatively high level of heterogeneity in shared drift with samples from other populations (Fig. S49, Fig. S48). Particularly interesting differences include the excess shared drift with Levantine samples found for the Nea Nikomedeia sample *Nea3* (the older) but not *Nea2* (the younger) when comparing these to other samples from the region (Fig. S48). Rev5, which is of similar age as *Nea3*, also shows a bit more of the Levantine ancestry (Fig. S48), suggesting a decrease in that ancestry over time. While there is no significant *f*-statistics when directly comparing *Rev5* to *Nea2* or *Nea3*, it does seem that all comparisons with the 4 *HG_Central* and *HG_West* genomes in our dataset show a trend for an excess shared drift with HG samples for *Rev5* and *Nea3* (Supp. Table 5).

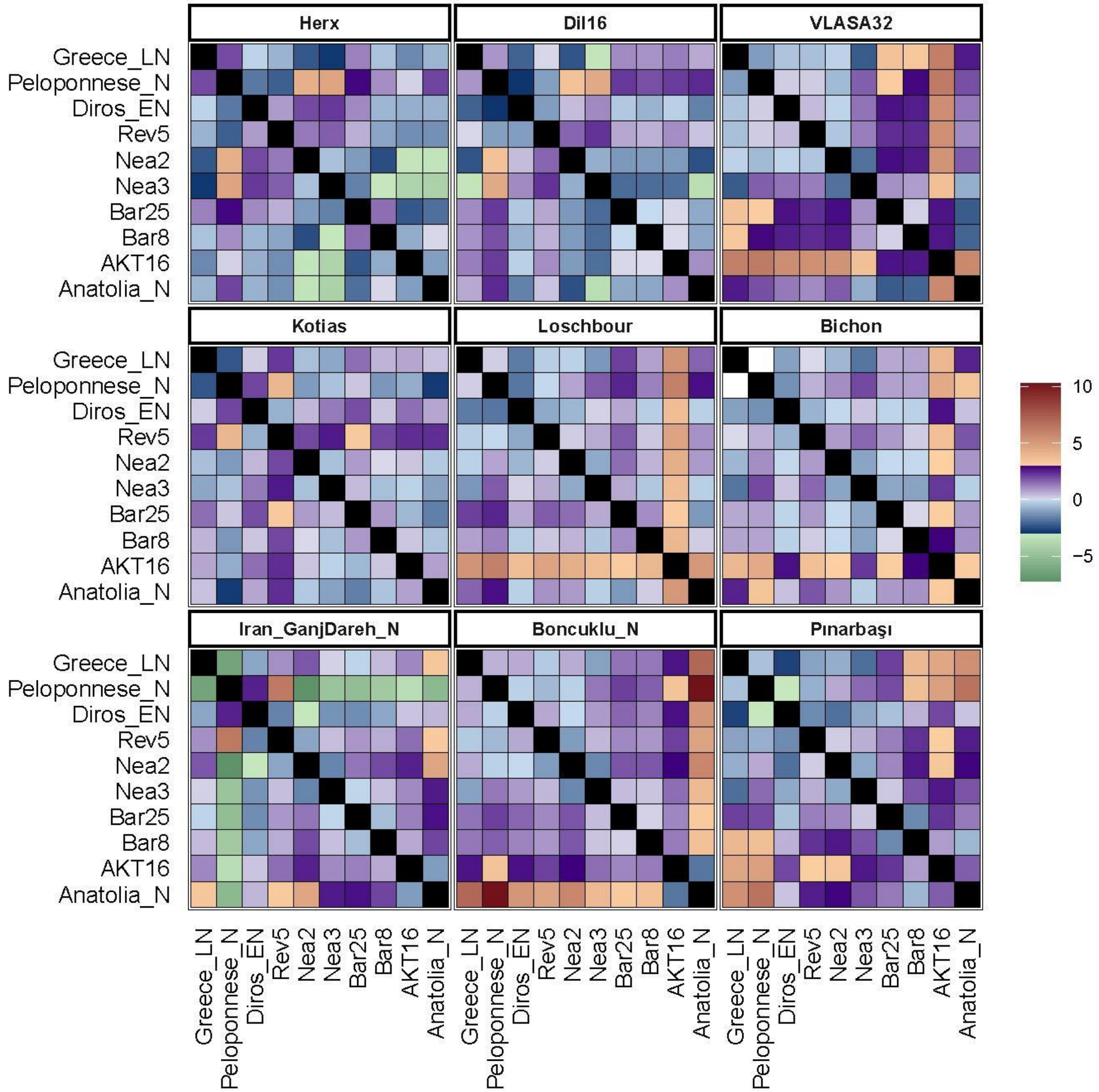


**Figure S49 - Heatmap of Z-scores of f-statistics of the form *D(Greece/NWAnatolia1; Greece/NWAnatolia2; Selected sample; Outgroup)*** with Mbuti as an outgroup and selected sample marked in the graph. Red shades indicate significant positive Z-scores above 3, purple shades are for Z-scores below 3 and above 0, blue shades for Z-scores below 0 and above -3 and green shades indicate significant negative Z-scores below -3.

**Konya Plain:** We investigated the relationships inside Anatolia by comparing our samples from Northern Greece and NW Anatolia to Pınarbaşı (Epipalaeolithic), Boncuklu (Neolithic), Kumtepe (Chalcolithic) and Tepecik-Çiftlik (Neolithic). A first result was that Tepecik-Çiftlik is cladding outside the branch represented by Boncuklu and Pınarbaşı as *D(Boncuklu, Pınarbaşı, Tepecik-Çiftlik, Outgroup)* is not significant (Z-score 0.76), while the two alternative *f*-statistics with either *Boncuklu* or *Pınarbaşı* as a sister group were highly significant (Z-score < -6.29, Supp. Table 5). *Tepecik-Çiftlik* also seems to clad outside a branch leading to *Boncuklu*, *Pınarbaşı* and all Neolithic Aegean samples as all *f*-statistics of the form *D(Aegean, Boncuklu / Pınarbaşı, Tepecik-Çiftlik, Outgroup)* were not significant (Z-scores < |2.051|), but 8 of 24 *f*-statistics of the form *D(Aegean, Tepecik-Çiftlik, Boncuklu / Pınarbaşı, Outgroup)* and *D(Boncuklu / Pınarbaşı, Tepecik-Çiftlik, Aegean, Outgroup)* were significant, in particular where *Aegean* is represented by the NW Anatolian samples and an early Neolithic sample from Peloponnese (*AKT16*, *Bar8*, *Anatolia_N* and *Diros_EN*, Supp. Table 5).
