## Supplementary material for "Demogenomic modeling of the timing and the processes of early European farmers differentiation": Supp. Tables Legend

**Supplementary Table 1 -** **Sample processing** including detailed information on library preparation, pipeline statistics and read stats for sequenced libraries and ancient reference samples.

**Supplementary Table 2** [**-**](https://docs.google.com/document/d/1kk9eEShr-ipZJppXdpyUG6cizK29AjQFknjfCoCXF9k/edit#table_AllSamples) **Genomic and sampling information about all 102 genomes used in our study**, including newly sequenced ancient genomes, ancient genomes from the literature and modern genomes selected within the SGDP panel.

**Supplementary Table 3** [**-**](https://docs.google.com/document/d/1kk9eEShr-ipZJppXdpyUG6cizK29AjQFknjfCoCXF9k/edit#table_HaplogroupsHirisPlex) **Detailed haplotype information for mtDNA and Y haplogroups**, including quality information and defining markers, for newly-sequenced individuals (in black) and already published individuals (in grey); and **results as reported by the HirisPlex webtool**, including all input files created for each newly-sequenced individual.

**Supplementary Table 4** [**-**](https://docs.google.com/document/d/1kk9eEShr-ipZJppXdpyUG6cizK29AjQFknjfCoCXF9k/edit#table_DemoResults) **Description of the parameters inferred from the tested models under the six different datasets.** For each dataset, we provide the maximum estimated log-likelihood of all models with the best supported model highlighted in grey, their relative likelihood derived from the AIC, the values obtained for the different parameters and the 95% confidence intervals under the best supported model.

**Supplementary Table 5** [**-**](https://docs.google.com/document/d/1kk9eEShr-ipZJppXdpyUG6cizK29AjQFknjfCoCXF9k/edit#table_Fstats) **Results of D-statistics in the form of D(Population1, Population2, Population3, Outgroup)** where we used Mbuti as Outgroup and Population1-3 were samples analysed in the study and relevant reference samples and populations.
